# Single-cell DNA replication dynamics in genomically unstable cancers

**DOI:** 10.1101/2023.04.10.536250

**Authors:** Adam C. Weiner, Marc J. Williams, Hongyu Shi, Ignacio Vázquez-García, Sohrab Salehi, Nicole Rusk, Samuel Aparicio, Sohrab P. Shah, Andrew McPherson

**Author notes:** These authors co-supervised this work (Sohrab P. Shah, Andrew McPherson). Contributing authors.

## Abstract

Dysregulated DNA replication is both a cause and a consequence of aneuploidy, yet the dynamics of DNA replication in aneuploid cell populations remains understudied. We developed a new method, PERT, for inferring cell-specific DNA replication states from single-cell whole genome sequencing, and investigated clone-specific DNA replication dynamics in >50,000 cells obtained from a collection of aneuploid and clonally heterogeneous cell lines, xenografts and primary cancer tissues. Clone replication timing (RT) profiles correlated with future copy number changes in serially passaged cell lines. Cell type was the strongest determinant of RT heterogeneity, while whole genome doubling and mutational process were associated with accumulation of late S-phase cells and weaker RT associations. Copy number changes affecting chromosome X had striking impact on RT, with loss of the inactive X allele shifting replication earlier, and loss of inactive Xq resulting in reactivation of Xp. Finally, analysis of time series xenografts illustrate how cell cycle distributions approximate clone proliferation, recapitulating expected relationships between proliferation and fitness in treatment-naive and chemotherapeutic contexts.

## 1 Introduction

DNA replication and cell cycle regulation are frequently disrupted as part of a cancer’s progression toward uncontrolled proliferation [1–3]. The resulting dysregulation increases replication stress and genomic instability, generating somatic copy number alterations (CNAs) and producing intratumoral heterogeneity that drives subsequent evolution [4, 5]. At a more granular level, the relative timing at which different regions of the genome replicate during sythesis (S)-phase of the cell cycle, known as replication timing (RT), is strongly associated with epigenomic features including 3D nuclear organization, chromatin state, and transcription, and cellular phenotype [6–10]. Structural variation and CNAs have been shown to impact epigenomic and chromatin state, and may also impact RT [11–14]. Additionally, specific genomic alterations confer fitness advantages, producing genetically distinct subclones with unique proliferation rates and thus more rapid progression through the cell cycle. Single-cell whole genome sequencing (scWGS) is a powerful method for studying clonal heterogeneity and CNAs, and has the potential to provide greater insight into DNA replication dynamics in aneuploid populations [15–20]. However, computational identification of S-phase cells and distinguishing replicating from non-replicating loci remains challenging due to the difficulty of distinguishing inherited somatic CNAs from transient DNA replication changes. Disentangling these two signals would improve the ability to study replication timing and proliferation rate of individual genetic subclones, leading to better understanding of how DNA replication drives and is further modulated by genomic instability.

We present a new method, Probabilistic Estimation of single-cell Replication Timing (PERT), to jointly infer single-cell copy number and replication states from scWGS data. PERT uses a Bayesian framework that models observed read depth as a combination of somatic copy number (CN), replication, and sequencing bias, enabling estimation of DNA replication profiles and cell cycle phase for individual cells. Unlike previous approaches for estimating single-cell replication timing (scRT) that assume the same CN profile for all cells in a sample [21–24], PERT is capable of modelling the clone- and cell-specific CNAs that are a common feature of genomicaly unstable cancers. Additionally, unlike scWGS cell cycle phase classifiers which rely on training data and existing RT information [16, 25], PERT provides unbiased estimates of RT and cell cycle phase which allows for analysis of previously uncharacterized cell types using any scWGS platform. These unique properties enable PERT to perform novel analysis such as estimating clone-specific proliferation rates and studying the interplay between RT and somatic CNAs during tumor evolution.

We used PERT to study DNA replication dynamics of genomically unstable cell lines and a collection of high-grade serous ovarian cancer (HGSOC) and triple negative breast cancer (TNBC) human tumors. First, since early and late RT loci are known to have different DNA damage and repair rates [26–28], we investigated the relationship between ancestral RT and the emergence of CNAs. Second, we modeled the relative impact of cell type, mutational signature, and ploidy on RT and the distribution of early vs late S-phase cells because these features have been shown to correlate with replication origin placement, replication stress response, perturbed epigenetic state, and 3D nuclear organization [7, 11–14, 29–31]. Third, we leveraged the fact that the inactive chromosome X allele (Xi) replicates very late within S-phase [22, 32] to identify recurrent patterns of Xi selection in TNBC and HGSOC tumors. Finally, since enrichment for S-phase cells is a marker for increased proliferation in histologic or transcriptional data modalities [33–36], we investigated the effect of chemotherapy and whole genome doubling (WGD) on the relationship between proliferation and evolutionary fitness at clone resolution.

## 2 Results

### Accurate estimation of single cell replication timing with PERT

The main methodological objective of PERT is to infer transient replication states and inherited somatic CN states from scWGS data. To do so, PERT implements a hierarchical Bayesian probabilistic graphical model and uses stochastic variational inference. Observed read depth (*Z*) is modelled as dependent on both latent CN (*X*) and replication (*Y*) states across all genomic loci (*N* cells x *M* bins) where the replication state depends on each cell’s time within S-phase (*τ*) and each locus’s average RT (*ρ*) (Fig. 1a, Extended Data Fig. 1a-e). Additional parameters govern the likelihood of the observed per-cell read depth (*Z*) given somatic CN and replication states (*X* + *Y*). After learning replication states in all cells, PERT then predicts S, G1/2, and low quality (LQ) phases based on the fraction of replicated loci and the quality of the cell’s predicted replication state. PERT is implemented using Pyro [37] and is freely available online with user tutorials. All terms in the graphical model as well as additional mathematical, inference, and implementation details can be found in the Methods.

**Fig. 1.**
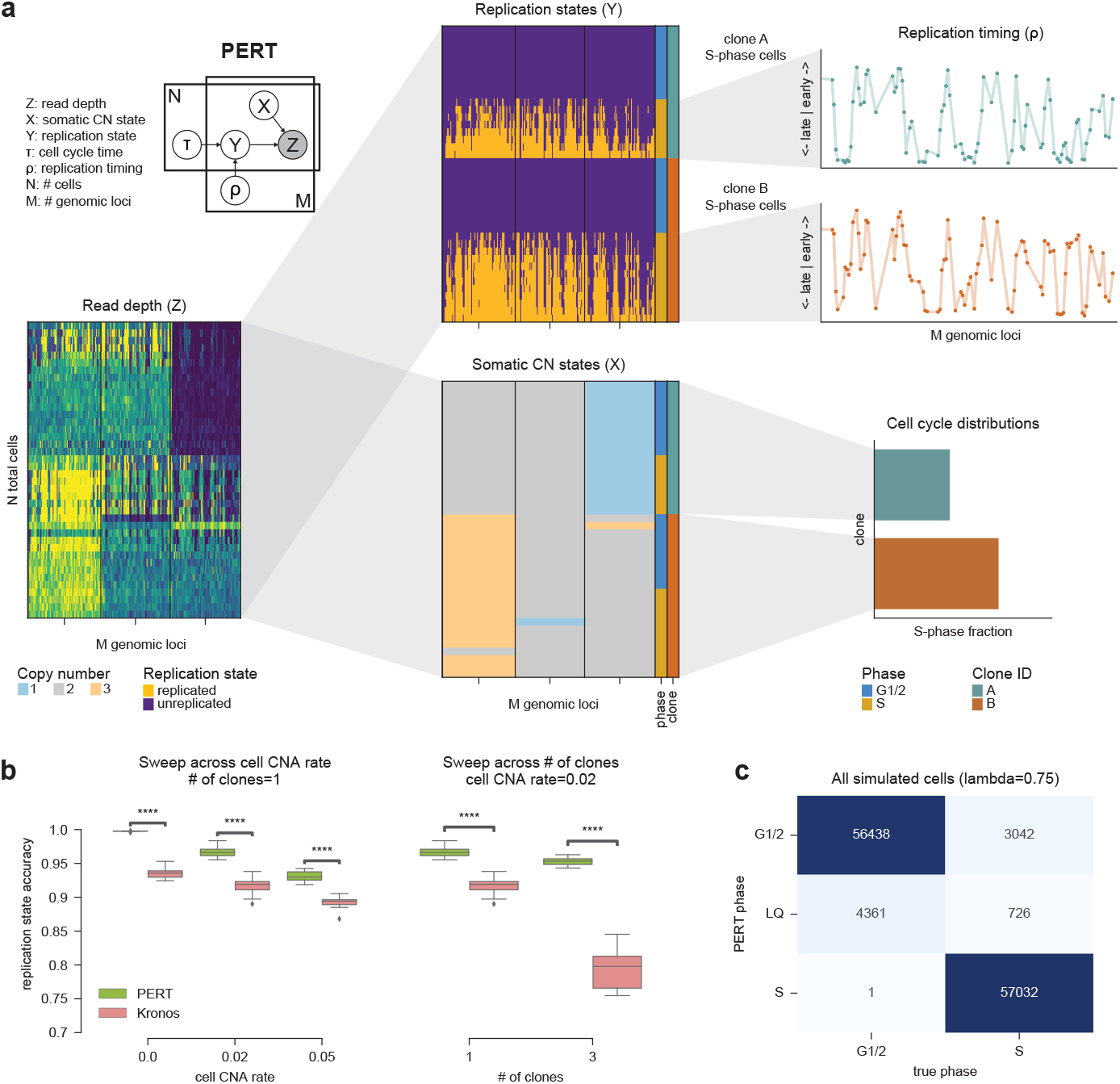
Overview of PERT. **a)** ERT takes scWGS binned read count as input (bottom left) and decomposes this signal into replication and somatic copy number states (center column). A simplified version of the PERT graphical model is shown in the top left. Downstream analysis tasks such as computing clone RT profiles and cell cycle distributions are shown in the right column. **b)** Perbin replication accuracy in S-phase cells for Kronos and PERT with clone and composite CN prior concentrations across varying numbers of clones and cell-specific CNA rates. **c)** Confusion matrix of true vs PERT cell cycle phase for simulated datasets with *λ* = 0.75.

We benchmarked PERT’s accuracy at inferring somatic CN, replication states, and cell cycle phase through quantitative simulation experiments. PERT outperformed the Laks *et al* classifier [16] for cell cycle phase prediction and Kronos [24] for scRT estimation in all simulated datasets. The performance gap between PERT and Kronos was significant (*p*_*adj*_ *<* 10^*-*4^) for all parameter combinations and increased as a function of cell CNA rate, number of clones, and noise *λ* (Fig. 1b). The agreement between each cell’s true and inferred fraction of replicated bins was used to classify cell cycle phase with 93% overall accuracy (97% accuracy excluding LQ cells) (Fig. 1c). Additional benchmarking information can be found in the Supplementary Notes 1-2. In summary, PERT significantly improves inference of scRT and phase, particularly in cases where CNAs arise with subclonal structure, allowing for exploration of replication dynamics in heterogeneous aneuploid populations.

### Validation of PERT in sorted diploid and aneuploid cell lines

Next we sought to validate whether PERT was sensitive to distinct clone-specific RT profiles in clonally heterogeneous samples. To do so, we performed *in silico* mixing experiments of scWGS data of two unrelated cell lines with ground truth cell cycle labels from fluorescence-activated cell sorting (FACS) based on DNA content. We combined lymphoblastoid GM18507 cells with diploid genomes (657 G1 cells, 585 S cells, 337 G2 cells, 1 clone) and breast cancer T47D cells with aneuploid genomes (703 G1 cells, 623 S cells, 522 G2 cells, 5 clones) [16] into one merged sample for analysis with PERT (Extended Data Fig. 2a). PERT found distinct CN profiles for S-phase cells of each line and its predicted phases were highly concordant with FACS (Fig. 2a,b). Both GM18507 and T47D samples were enriched for mid-S-phase cells (Extended Data Fig. 2b). Cell line ‘pseudobulk’ RT profiles showed that 15% (794/5258) of loci had an absolute RT difference *>*0.25 between GM18507 and T47D, consistent with each cell line having a unique RT program (Fig. 2c). RT has been shown to be influenced by nuclear organization, with genomic loci in inactive chromatin being late replicating and active chromatin being early replicating [6–8, 22]. Consistent with this, we found cell line specific correlations between the inferred RT profiles and Hi-C A/B (active/inactive) compartments of T47D and other lymphoblastoid cell lines [38] (Fig. 2d), highlighting the biological relevance of the RT differences identified by PERT. To ensure that the latent RT variable (*ρ*) did not prevent inference of cell line RT profiles, we ran PERT independently for each cell line and found that both cell lines had RT correlations of 0.99 between their merged and split PERT runs (Fig. 2e, Extended Data Fig. 2c). Similarly, PERT inferred accurate clone-specific RT profiles for simulated data in which each clone had a unique ENCODE cell line RT profile [38] (Extended Data Fig. 2d,e). These experiments demonstrate that PERT can accurately identify clone-specific RT profiles within the same sample.

**Fig. 2.**
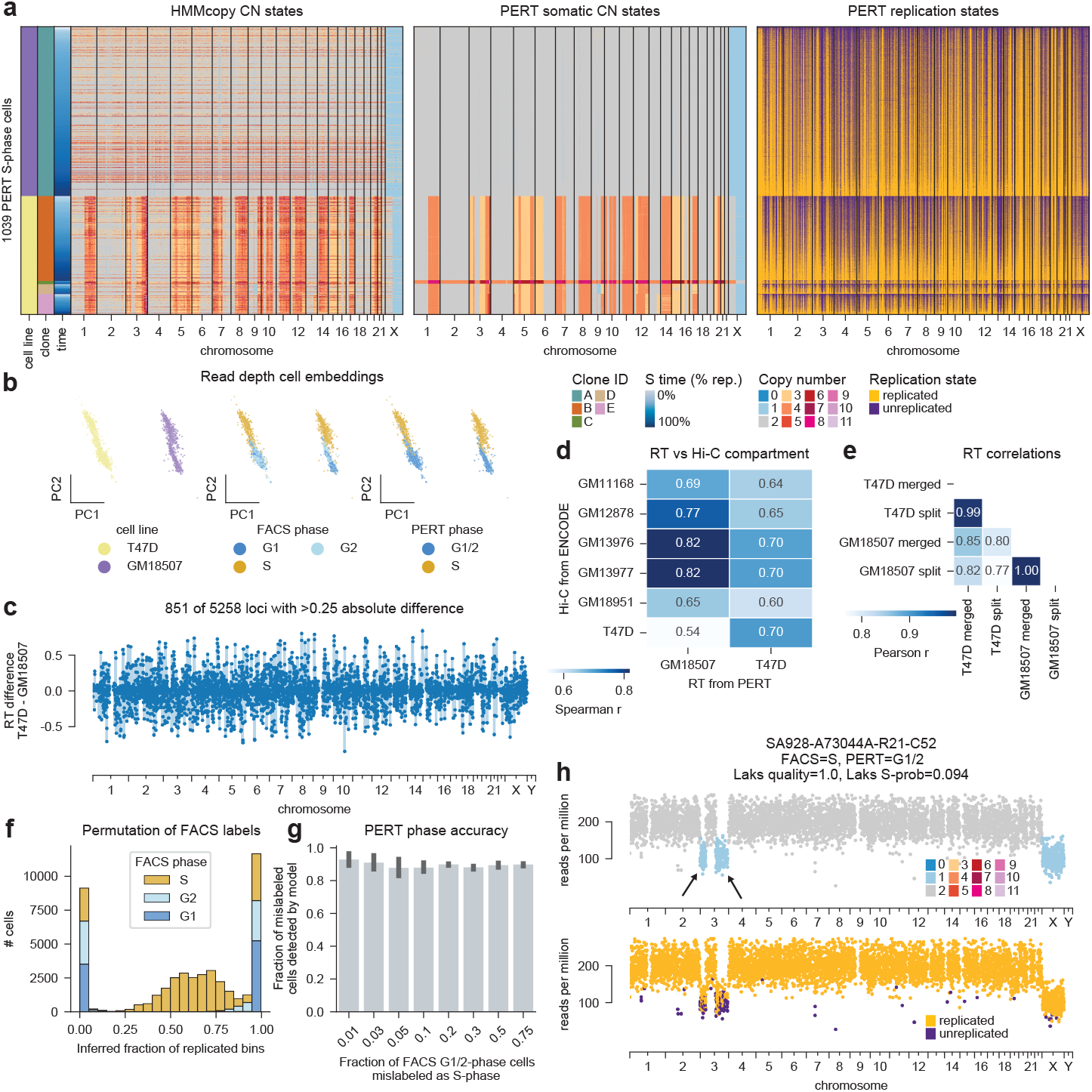
PERT identifies cell line-specific RT profiles in cell cycle sorted scWGS data. **a)** HMMcopy (left), PERT copy number (middle), and PERT replication (right) states for GM18507 and T47D cells predicted as S-phase by PERT. **b)** PCA cell embeddings of read depth where cells are colored by cell line (left), FACS cell cycle phase (middle), and PERT predicted phase (right) **c)** Difference between inferred T47D and GM18507 RT profiles. Positive values have earlier RT in T47D than GM18507. **d)** Spearman correlation between inferred T47D and GM18507 RT profiles and Hi-C A/B compartment scores from ENCODE. All cell lines with the ‘GM’ prefix are lymphoblastoids. **e)** Pearson correlation between inferred T47D and GM18507 RT profiles when the cell lines are merged into one sample or split when running PERT. **f)** Histogram of PERT’s inferred fraction of replicated bins per cell for all cells in permuted datasets, colored by true FACS phase. **g)** Fraction of FACS=G1/2 cells mislabeled as S-phase predicted as PERT=G1/2 across all permutation rates. **h)** Example of a FACS=S PERT=G1/2 GM18507 cell. Bins are colored by HMMcopy (top) and PERT replication (bottom) states. Arrows point to cell-specific CNAs. Title includes quality score and S-phase probability from Laks *et al* classifiers.

With this same data, we investigated PERT’s robustness to poor initialization of preliminary G1/2 vs candidate S-phase cells. Given the high per-cell failure rates of sequencing and CN calling in scWGS, we were concerned that the inclusion of too many true G1/2-phase cells during initialization would bias RT and phase estimates. We thus devised a permutation experiment in which a subset of GM18507 and T47D FACS G1/2-phase cells were mislabeled as candidate S-phase cells during initialization to examine whether PERT successfully recovered them as G1/2-phase. We found that *>*90% of all mislabeled cells were accurately recovered (predicted G1/2) across all permutation datasets without compromising identification of S-phase cells and cell line-specific RT profiles (Fig. 2f,g, Extended Data Fig. 2f). Mislabeled cells which were predicted to be in S-phase were disproportionately G2 by FACS and 80-95% replicated with orthogonal per-cell features concordant with late S-phase (Fig. 2f, Extended Data Fig. 2g). Additionally, we found many cell-specific CNAs in the set of FACS S-phase cells predicted to be G1/2-phase (Fig. 2h). We hypothesize that many discrepancies between FACS and PERT phases were FACS errors, which is known to have S-phase purities ranging from 73-93% [39], since cells with unique CNAs possess higher or lower DNA content than other cells in the same phase. Our evidence suggests that prediction of cell cycle phase using PERT is non-inferior, if not superior, to experimental sorting, especially in heterogeneous aneuploid populations.

### Replication timing predicts future CNAs

Next, we investigated the relationship between CNAs and RT by applying PERT to previously published scWGS data of mammary epithelial 184-hTERT cell lines (hereafter referred to as hTERTs) [40]. The hTERT samples were engineered using CRISPR/Cas9 to ablate TP53, TP53/BRCA1, and TP53/BRCA2, and passaged *∼* 60 times with intermediate scWGS sampling to capture accrual of aneuploidies (Fig. 3a). The initial investigation of this dataset revealed clonal expansions of cell populations with increasing levels of CNAs but excluded analysis of S-phase cells. Here we analyzed all cells with PERT and found that, unlike the FACS cell lines which were artificially enriched for mid-S-phase cells, the unsorted hTERTs had more late than early S-phase cells which agrees with reports that most loci replicate during early S-phase while very late RT loci take much longer to replicate [41, 42] (Fig. 3b).

**Fig. 3.**
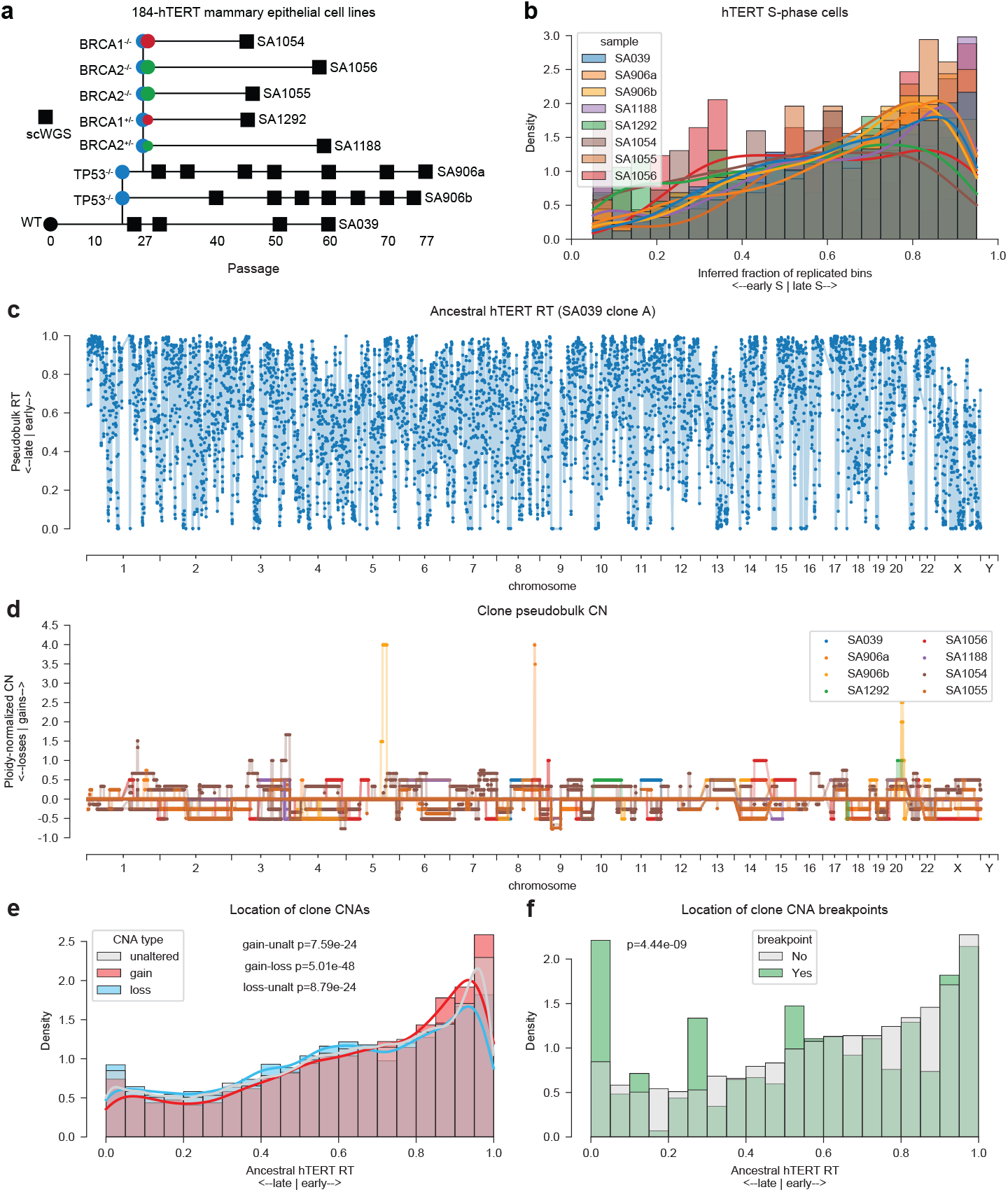
Relationship between copy number alterations and replication timing. **a)** hTERT genotype lineage diagram showing progression from WT to mutant alleles. Boxes represent passages in which scWGS libraries were generated. **b)** Distribution of inferred fraction of replicated bins across all S-phase cells in hTERT cell lines. **c)** RT profile of hTERT SA039 clone A (diploid WT). **d)** CN profiles for all hTERT clones, normalized by ploidy. Values *>* 0 are gains, *<* 0 are losses, and = 0 are unaltered. **e-f)** Distribution of hTERT SA039 clone A (diploid WT) RT values split by **e)** clone pseudobulk CNA types and **f)** the presence of clone pseudobulk CNA breakpoints.

We then used PERT results to interrogate whether RT influences CNA acquisition. We computed a reference RT profile from the ancestral hTERT WT (TP53 and BRCA1/2 WT) population with no CNAs (SA039 clone A) (Fig. 3c, Extended Data Fig. 3a-c). Counting gain, loss and unaltered bins in clone pseudobulk CN profiles that descended from this ancestral WT population, we found that gains preferentially arise from early RT loci, losses from late RT loci, and CNA breakpoints from late RT loci (*p*_*adj*_ *<* 10^*-*4^) (Fig. 3d-f). These results support previous reports of common fragile sites being enriched in late RT loci [43, 44] and were reproduced when using sample pseudobulk CN profiles (Extended Data Fig. 3d-f). The association between RT and emergence of clone- and sample-specific CNAs recapitulates existing evidence these two phenomena are governed by the same underlying processes.

### Global model for replication timing variability between clones

To understand the relative impact of cell type and other covariates on clone-specific RT, we applied PERT to a wider cohort of scWGS datasets with diverse genomic properties. The assembled metacohort comprised 6 TNBC tumors, 13 HGSOC tumors, 3 cancer cell lines, and the previously described hTERTs [16, 40, 45]. Samples had been labelled as homologous recombination deficiency (HRD), fold-back inversion (FBI), and tandem duplicators (TD) by previous mutation signature analysis, and contained both whole-genome doubled (WGD) and non-genome doubled (NGD) clones. We focused our analysis on the 102 unique clones with *>*20 S-phase cells (Fig. 4a). We first investigated the time distribution of S-phase cells across signature and ploidy, finding that both WGD and mutation signatures consistent with replication stress exhibit higher fractions of late S-phase cells [29–31] (Fig. 4b,c). Clustering the pairwise Pearson correlations between clone RT profiles revealed striking sample and cell type specificity (Fig. 4d). We then implemented a factor model to jointly learn weights representing relative covariate importance to clone level RT, and latent profiles representing covariate-specific RT differences across the genome (Methods). The constant term representing global RT was estimated to have the highest importance, as expected given that RT is largely conserved across cell types. Estimated covariate importance from most to least important was sample, cell type, ploidy, and signature (Fig. 4e). Ploidy and signature were both an order of magnitude lower in importance than sample or cell type, suggesting that the higher proportion of late S-phase cells identified for clones with WGD and replication stress associated signatures did not necessarily coincide with large RT changes. Finally, we computed the mean RT of each chromosome across all cell types, and did the same for ENCODE bulk RT (RepliSeq) data [38]. We found high agreement in cell type RT on most chromosomes with chromosome X having the highest variability (Fig. 4f, Extended Data Fig. 4), prompting further analysis on the relationship between X-inactivation and RT.

**Fig. 4.**
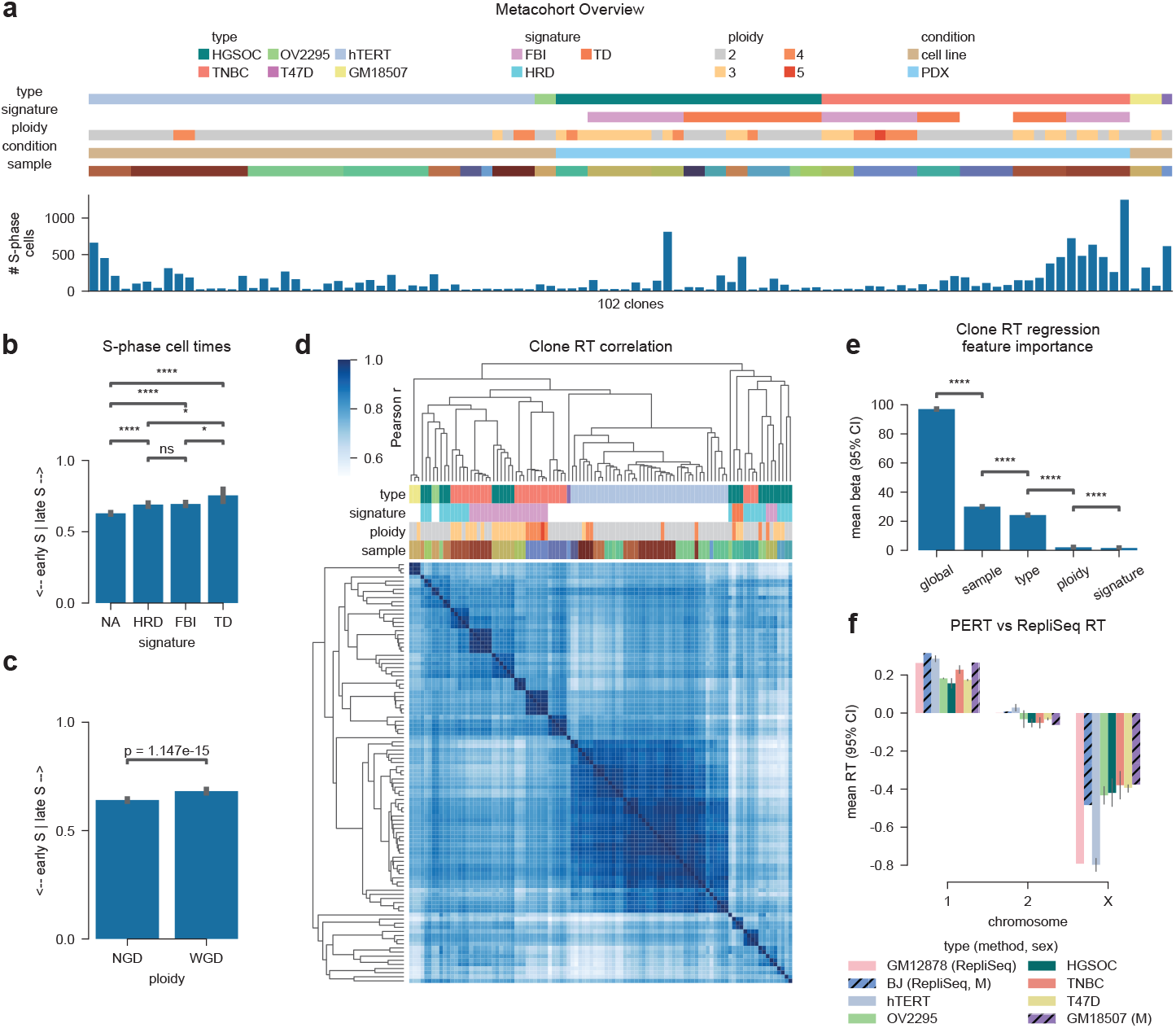
Replication timing variability across cell type, signature, and ploidy. **a)** Overview of metacohort containing various cell lines and human tumors. Each clone is annotated based on its cell/tumor type, mutational signature, condition (cell line, PDX), sample, and number of S-phase cells. **b-c)** Mean S-phase time for all S-phase cells belonging to each **b)** mutational signature or **c)** ploidy group. S-phase time is defined as the fraction of replicated bins per cell. Bars represent mean values across all cells with error bars showing 95% confidence intervals. **d)** Pairwise Pearson correlations between all clone RT profiles in the metacohort. Rows and columns are clustered in the same order. Color bars for each column match the legneds shown in **a. e)** Posterior distribution of covariate importance terms (*β*s) from a model which jointly infers covariate-specific RT profiles and importance terms directly from the matrix of clone RT profiles. **f)** Mean RT across cell types and chromosomes 1, 2, and X. GM12878 and BJ cell type RT data is derived from ENCODE RepliSeq data; all other cell type RT data is derived from scWGS PERT analysis. Male cell lines are noted with black crosshatches. Error bars represent the 95% confidence intervals over the per-chromosome mean RT when multiple clones are present.

### Chromosome X replication timing shifts measure the ratio of active to inactive alleles and reveal selection bias in HGSOC and TNBC tumors

Given that X-inactivation produces a late replicating inactive allele (Xi) and an early replicating active allele (Xa) [22, 32], we hypothesized that the greatest RT shifts would occur from CNAs which disrupted the 1:1 balance of Xa to Xi alleles. To study the relationship between RT and X-inactivation we ran PERT and SIGNALS [40], a single-cell allele-specific copy number caller, on the previously described hTERT cell lines and compared the SIGNALS B-allele frequencies (BAF) to the RT difference between chrX and all autosomes. ChrX RT and DNA BAF were negatively correlated at both sample and clone resolution, with delayed RT associated with balanced allelic copy number (BAF=0.5), and loss of the B-allele (BAF*<*0.5) shifting RT earlier (Fig. 5a-c). A decrease in chrX BAF for S-phase cells compared to G1/2-phase cells provides further evidence that A=Xa and B=Xi in all hTERT samples (Extended Data Fig 5a-c). These results are concordant with the loss of Xi and retention of Xa producing a shift towards earlier RT, and highlight PERT’s ability to associate the SIGNALS A- and B-allele labels with Xa and Xi epigenetic states.

**Fig. 5.**
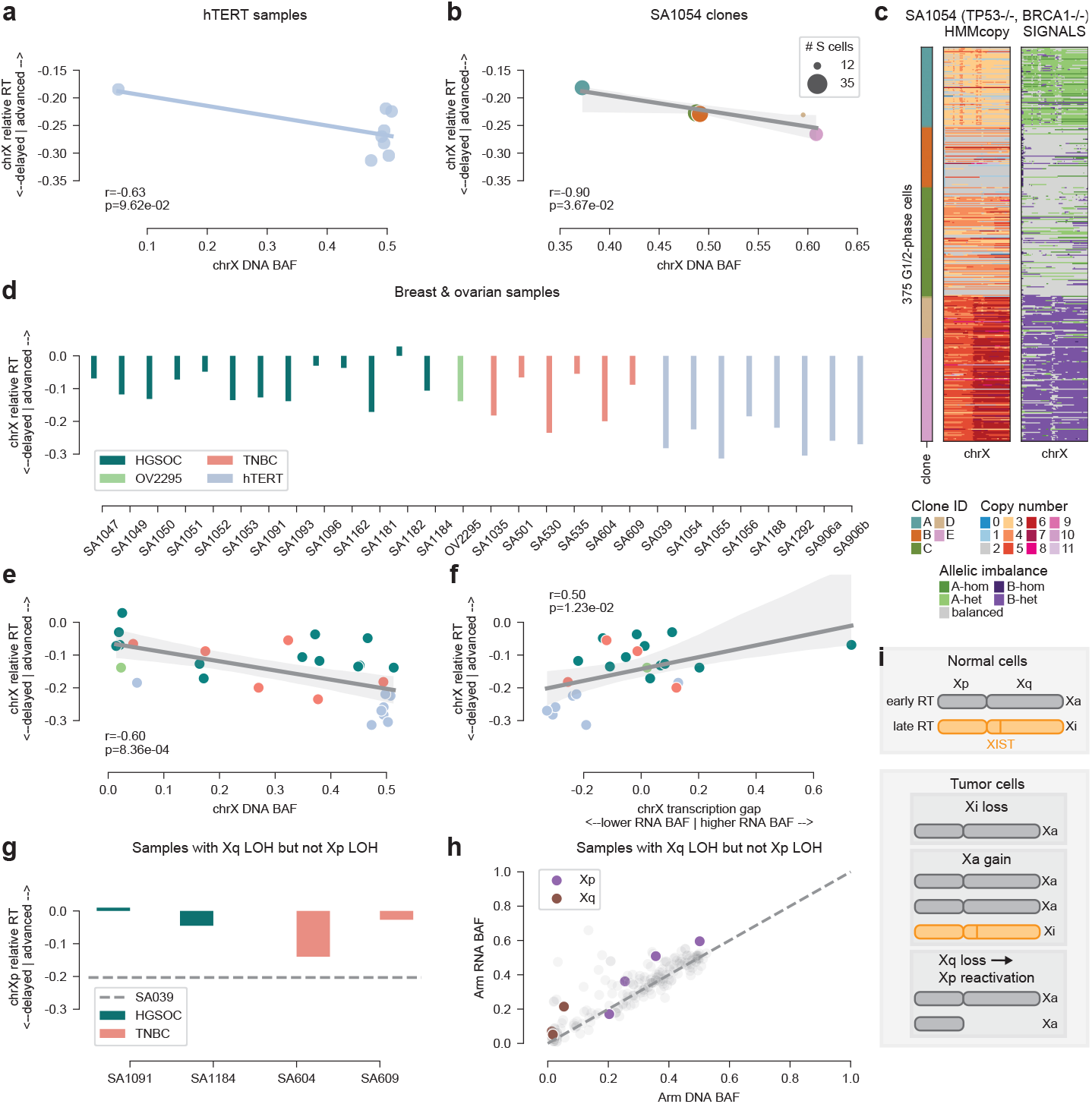
Replication timing shifts allow for phasing of chrX inactivation and reveals Xa*>*Xi selection in HGSOC and TNBC. **a)** ChrX B-allele frequency from SIGNALS analysis of scDNA data vs the relative RT of chrX compared to autosomes in the same sample. All points are unique hTERT samples which share the same SIGNALS phasing. chrX relative RT values of 0 represent cases in which chrX replicates at the same time as all autosomes and negative values imply that chrX replicates later than the autosomes. SIGNALS assigns the major (more prevalent) allele as A and minor allele as B at each haplotype block. **b)** chrX B-allele frequency vs relative RT for clones in hTERT sample SA1054. **c)** Total copy number and allelic imbalance states in chrX for all G1/2-phase cells in sample SA1054. Clone IDs are annotated in the left-hand column. **d)** ChrX relative RT for all samples, colored by cell type. hTERT and OV2295 samples are cell lines and HGSOC and TNBC samples are PDXs. Note that OV2295 is a cell line derived from an HGSOC tumor. **e)** chrX B-allele frequency vs relative RT for all samples shown in **d**. SIGNALS phasing was performed independently for each sample. **f)** Comparison of the chrX RNA BAF - DNA BAF “transcription gap” (x-axis) vs relative RT (y-axis) of a given sample. Positive transcription gap means a sample has more transcription of the B-allele than one would expect from looking at the DNA BAF of said sample. **g)** Xp relative RT for subset of samples with LOH (BAF=0) on the Xq arm (contains *XIST* locus) but not on the Xp arm. The horizontal line represents Xp relative RT for hTERT WT sample SA039 which is balanced with the B-allele being inactive on both arms. **h)** Mean DNA vs RNA BAF for each chromosome arm for samples with Xq LOH and balanced Xp. All autosomes arms are colored light grey, chrX arms are colored to illustrate their 1:1 relationship between gene dosage and transcription. **i)** Schematic demonstrating how tumors achieve Xa*>*Xi ratios through Xi loss, Xa gain, and X-reactivation.

We then assessed the degree of chrX RT allelic imbalance in 19 TNBC and HGSOC samples. All tumors had earlier chrX RT than the allelically balanced (BAF=0.5) hTERT samples, with a negative correlation between chrX RT and DNA BAF, suggesting that many of these tumors had more copies of Xa than Xi (Fig. 5d,e, Extended Data Fig. 5d). Many of these allelic imbalances were fully clonal events and arose from both loss of Xi (4/19 clonal full chrX LOH, 4/19 clonal Xq LOH, 2/19 clonal partial LOH, 1/19 subclonal LOH) and gain of Xa (1/19 clonal full chrX, 4/19 subclonal and/or partial) (Additional File 1), suggesting that Xa*>*Xi imbalances emerge early in tumor evolution and that both Xa gain and Xi loss are independently favorable events. For samples with matching scRNA-seq, we compared SIGNALS RNA BAF with DNA BAF to investigate the relationship between allelic dose and allele-specific transcription of each chromosome. While autosomes maintained a 1:1 relationship between RNA and DNA BAF, samples with Xi alleles present had lower RNA BAF than DNA BAF, indicating lower Xi expression (Extended Data Fig. 5e). We termed the difference between chrX RNA and DNA BAF as the “transcription gap” and found that it positively correlated with chrX RT for each sample (Fig. 5f), confirming that the unexpected expression from the B-allele on X coincided with earlier X replication.

Next we sought to identify mechanisms that would explain unexpected expression of the inactive X allele. We identified loss of heterozygosity (LOH, BAF=0) on Xq but not Xp in 4 of 19 tumors (2 HGSOC, 2 TNBC, Extended Data Fig. 5f). Given that X-inactivation proceeds *in cis* through the transcription of *XIST* on the Xq arm, we hypothesized that loss of the Xq B-allele enabled re-activation of the Xp B-allele. Using PERT, we found that the Xp arm of these samples replicated much earlier than the corresponding locus in the hTERT WT sample (SA039) which had intact *XIST* transcription on the B-allele (Fig. 5g). We then compared the DNA and RNA BAFs at chromosome-arm level resolution for these samples and found that the Xp arm maintained a 1:1 ratio between gene dosage and transcription, confirming our hypothesis that these Xp B-alleles were reactivated after loss of the Xq B-alleles (Fig. 5h). These results demonstrate that loss of Xi, gain of Xa and, in some cases, reactivation of Xi are evolutionarily favorable events in HGSOC and TNBC tumors (Fig. 5i).

### Clone cell cycle distributions reflect proliferation rate and cisplatin sensitivity

Intratumoral evolution and clonal expansions are driven by high proliferation rates, providing a relative fitness advantage to highly proliferative cells in the treatmentnaive setting and greater sensitivity to platinum-based chemotherapies [46]. We thus leveraged PERT’s ability to estimate the cell cycle phase distributions to examine on- and off-treatment fitness of individual clones. We confirmed that cell cycle distributions correctly approximate proliferation rate by observing that high PERT G1/2-phase fractions correlate with low proliferation and high scRNA G1-phase fractions in three published gastric cancer cell lines with co-registered doubling times, 10X scWGS, and 10X scRNA measurements [17] (Extended Data Fig. 6, Additional File 2). We then analyzed time-series scWGS generated from serially propagated TNBC patient derived xenografts (PDXs) with and without cisplatin treatment [45] (Fig. 6a) to investigate whether PERT can assess proliferative fitness of tumor clones under therapeutic selective pressure. Previous analysis of these data had revealed an inversion of the clonal fitness landscape upon cisplatin exposure but had not identified any genotypic or phenotypic features to explain such an inversion. We used the relative abundance of each clone within each cell cycle phase to compute continuous S-phase enrichment (SPE) scores for all clone x timepoint combinations (Fig. 6b-e, Extended Data Fig. 7, Methods). Clone SPE scores were positively correlated with clone expansion between adjacent time points in untreated samples, but negatively correlated in treated samples, consistent with increased cell death for S-phase cells and fitness advantages of slow proliferation in the presence of platinum chemotherapy (Fig. 6f).

**Fig. 6.**
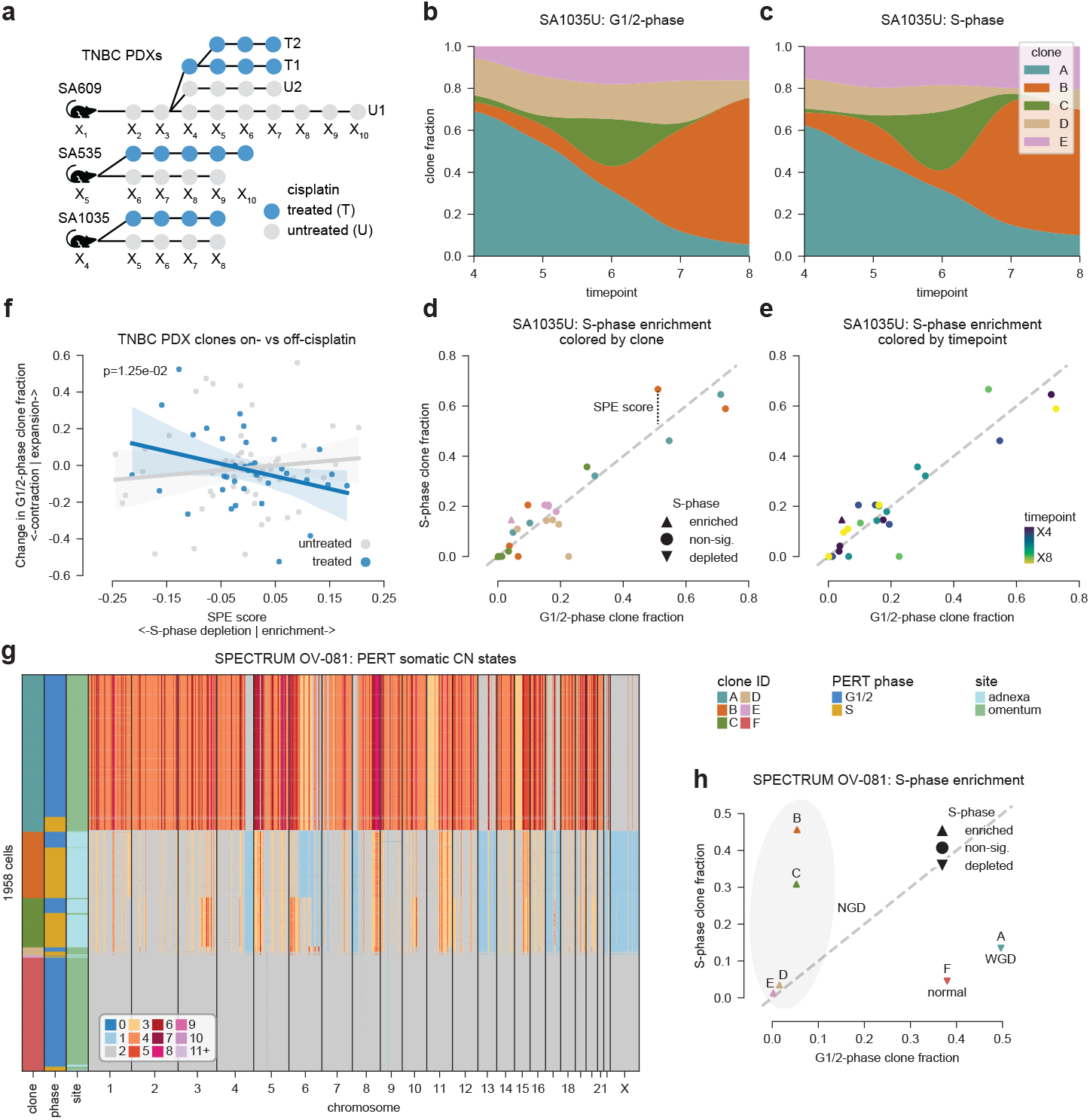
Relationship between clone cell cycle distribution and evolutionary fitness. **a)** Schematic of time-series scWGS sampling for untreated and cisplatin-treated TNBC PDXs. **b-e)** Representative SA1035 untreated sample. **b-c)** Relative fraction of each clone within G1/2- and S-phase cells. **d-e)** Comparison of each clone’s fraction in S-vs G1/2-phase populations at each timepoint. Dashed gray line represents equal prevalence in both phases. Triangles denote clone and timepoint combinations significantly (*p*_*adj*_ *<* 0.01) enriched or depleted for S-phase cells via hypergeometric test. Distance from the dashed gray line represents each point’s continuous SPE score. **f)** Relationship between SPE score and clone expansion between timepoints *t* and *t* + 1 for all TNBC PDX clone and timepoint combinations with *>* 10 G1/2-phase cells, split by cisplatin status. Lines represent linear regression fits with shaded areas representing 95% confidence intervals. **g)** PERT somatic copy number states from multi-site scWGS sequencing of HGSOC patient OV-081. Rows are annotated by their clone ID, PERT predicted cell cycle phase, and site of collection from the primary debulking surgery. Contaminating normal cells are included as clone F for reference **i)** Clone fraction in S-vs G1/2-phase populations for each OV-081 clone. Each clone is annotated by tumor/normal and WGD/NGD status.

To better understand the impact between WGD and proliferation rate in human tumors, we ran PERT on scWGS data from HGSOC patient OV-081 in the MSK SPECTRUM cohort [47]. Patient OV-081 presented with a primary tumor in the left adnexa consisting of mostly NGD tumor cells and a metastasis in the omentum consisting of mostly WGD tumor cells (Fig. 6g, Extended Data Fig. 8a). The NGD tumor clones (B-E) were significantly enriched for S-phase cells (positive SPE) while WGD (A) and normal (F) clones were significantly depleted for S-phase cells (negative SPE) (Fig. 6h). The discrepancy in clone SPE was validated with scRNA-seq data showing that 53% of tumor cells are cycling in the NGD left adnexa (22% S-phase, 31% G2/M-phase) but only 33% of tumor cells are cycling in the WGD omentum (16% S-phase, 17% G2/M-phase, Extended Data Fig. 8b). The relative ordering of SPE scores between NGD, WGD, and normal clones was preserved within the omentum alone (Extended Data Fig. 8c), confirming that this is unlikely to be a site-specific batch effect. This data suggests that WGD cells proliferate slower than NGD cells in this treatment-naive tumor.

## 3 Discussion

Here we demonstrate that somatic copy number change and DNA replication states can be jointly inferred from single cell whole genome sequence data using PERT. We show PERT’s compatibility with scWGS data produced by both the direct library preparation (DLP+) and 10X Chromium platforms, in addition to its flexibility to handle both lower resolution (500kb) and higher resolution (20kb) bin sizes. Although there is no limit to the size of clones that may be analyzed by PERT, accurate estimation of RT and cell cycle phase requires a sufficient number of S-phase cells within a sample as PERT learns RT *de novo*. For this reason, certain samples with *<* 100 S-phase cells or *<* 300 total cells were removed from further analysis. Finally, PERT is unsuitable for scWGS data generated from sorted G1/2 populations [19, 20].

PERT analysis of scWGS data from cell lines, xenografts, and tumor samples highlighted the complex relationship between somatic CNAs and RT. Analysis of RT in relation to subsequent CNAs revealed that copy number losses and breakpoints preferentially emerge from late RT loci while gains from early RT loci. These results are in agreement with findings from the PCAWG consortium [28] and could be explained by mechanisms of over- and under-replication or reflect differential fitness of gains vs losses in gene-rich early vs gene-poor late RT loci, respectively [2, 14, 41, 42]. In all samples, we found more late S-phase cells than early S-phase cells; however, this effect was more pronounced in clones with properties associated with replication stress such as the tandem duplication mutation signature or whole genome doubling. This result could be a consequence of how long it takes different DNA repair mechanisms to repair double strand breaks and stalled replication forks that arise in cells with increased genomic instability [29–31]. Additionally, this result implies a shorter time window between the end of replication (S/G2-phase boundary) and the start of mitosis (G2/M boundary), increasing the likelihood for missegregations. We found that RT is highly conserved within cell types across our metacohort of 102 clones despite highly variable copy number, suggesting that heritable RT may be helpful for identifying the cell-of-origin in tumors. We also found that CNAs on chromosome X recurrently produced Xa*>*Xi allelic imbalances in HGSOC and TNBC tumors, impacting RT and allele-specific expression, with evidence of Xp-reactivation via Xq LOH . These findings agree with similar reports of Xi loss, Xa gain, and Xp reactivation in breast, ovarian, and other female-specific or -enriched cancer types [48–50]. We observed some form of Xi loss in 8 of 19 HGSOC and TNBC cases and postulate that delayed Xi replication may increase the likelihood that a cell undergoes mitosis before replication. Our results implicate a role of chromosome X reactivation in female reproductive cancers.

Quantification of clone-specific cell cycle distributions allowed us to study the relative proliferation rate of tumor subclones. In serially propagated, drug-treated TNBC PDXs, we found that highly proliferative clones expanded at the next timepoint in the untreated context and contracted in the cisplatin-treated context. This suggests that accurate prediction of subclonal cell cycle phase distributions may be helpful for identifying senescent or hyperproliferative clones [51–53]. Furthermore, we believe that using cell cycle distributions to study which clones will respond to chemotherapy can provide complementary information to other genomic features such as gain of oncogenes, loss of tumor suppressors, and WGD which can have variable phenotypic impacts [47, 54, 55]. Finally, we found that in an HGSOC patient, a metastatic WGD clone had a slower proliferation rate than the NGD clones found in both primary and metastatic sites. This finding agrees with observations that the selective advantages of WGD can be conferred through slower but more robust growth (potentially via evasion of cell cycle checkpoints or immune surveillance) [56], prompting further study on the phenotypic consequences of WGD.

In summary, PERT offers a statistical framework with which to study copy number driven evolution and replication dynamics in cancer cells. Combining PERT with future scWGS generated for larger, more diverse cohorts will allow investigation into the relationship between DNA replication and genomic instability, providing insights into each tumor subclone’s etiology, evolutionary fitness, and drug sensitivities.

## 4 Methods

### PERT model

The input for PERT is binned read depth (*Z*) and called CN states for all scWGS cells. Input CN states are obtained through single-cell CN callers such as HMMcopy [15, 57] or 10X CellRanger-DNA [17, 18]. PERT first identifies a set of high-confidence G1/2-phase cells where the input CN states reflect accurate somatic CN. All remaining cells have their input CN states dropped as they are initially considered to have unknown CN states and cell cycle phase. Most S-phase cells should be present in the unknown initial set. High-confidence G1/2-phase cells are phylogenetically clustered into clones based on CN using methods such as sitka [58] or MEDICC2 [59]. Optionally, users can provide their own sets of clustered high-confidence G1/2-phase and unknown cells. These sets of cells are passed into a probabilistic model which infers somatic CN (*X*) and replication states (*Y*) through three distinct learning steps. In Step 1, PERT learns parameters associated with library-level GC bias (*β*_*µ*_, *β*_*σ*_) and sequencing overdispersion (*λ*) by training on high-confidence G1/2 cells (Extended Data Fig. 1c). Step 1 conditions on CN (*X*), replication (*Y*), and coverage/ploidy scaling terms (*µ*) because input CN states are assumed to accurately reflect somatic CN states and all bins are unreplicated (*Y* = 0) in high-confidence G1/2 cells. Once *β*_*µ*_, *β*_*σ*_ and *µ* have been learned in Step 1, we can condition on them in Step 2 (Extended Data Fig. 1d). Step 2 learns latent parameters representing each cell’s time in S-phase (*τ*_*n*_), each locus’s replication timing (*ρ*_*m*_), and global replication stochasticity (*α*) to compute the probability that a given bin is replicated (*Y*_*n,m*_ = 1) or unreplicated (*Y*_*n,m*_ = 0). Only unknown cells are included in Step 2. Prior belief on each unknown cell’s CN state is encapsulated using a prior distribution (*π*) which has concentration parameters (*η*) conditioned on the input CN of the most similar high-confidence G1/2 cells. CN prior concentrations are set for each cell by using the consensus CN profile of the most similar G1/2 clone or a composite scoring of the most similar G1/2 clone and cell CN profiles (Extended Data Fig. 1f) A full list of model parameters, domains, and distributions can be found in Extended Data Fig. 1b. Step 3 is an optional final step which learns CN and replication states for high-confidence G1/2 cells (Extended Data Fig. 1e). This step is necessary to determine if any S-phase cells are present in the initial set of high-confidence G1/2 cells. Step 3 conditions on replication timing (*ρ*) and stochasticity (*α*) values learned in Step 2 to ensure that such properties are conserved between both sets of cells.

PERT is designed for scWGS data with coverage depths on the order of 0.01-0.1x and thus 500kb bin sizes are used by default in this manuscript; however, the model can be run on count data of any bin size as long as sufficient memory and runtime are allocated. We demonstrate PERT’s ability to run on 10X scWGS data at 20kb resolution in Additional File 2.

### Equations for Step 1

Given that we have accurate CN caller results for high-confidence G1/2 cells, we can solve for each cell’s coverage/ploidy scaling term *µ*_*n*_ and condition on it,

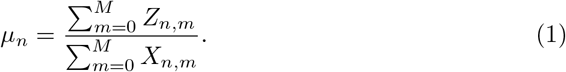

The latent variables are arranged together in function block *f* through the following equations to produce the bin-specific negative binomial event counts *δ*_*n,m*_. The GC bias rate of each individual bin (*ω*_*n,m*_) depends on the GC content of the locus (*γ*_*m*_) and the GC bias coefficients (*β*_*n,k*_) for the cell,

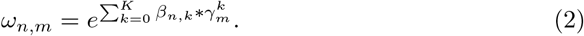

The expected read count per bin is computed as follows:

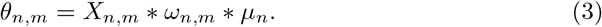

The expected read count per bin is then used in conjunction with the negative binomial event success probability term (*λ*) to produce a number of negative binomial event count for each bin,

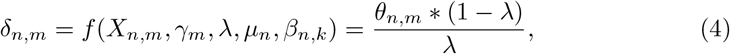

where we place the constraint *δ*_*n,m*_ *≥* 1 to avoid sampling errors in bins with *θ*_*n,m*_ *≈* 0. Finally, the read count at a bin is sampled from an overdispersed negative binomial distribution *Z*_*n,m*_ *≥* NB(*d*_*n,m*_, *λ*) where the expected read count for *Z*_*n,m*_ is *θ*_*n,m*_ and the variance is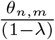.

### Equations for Steps 2-3

Steps 2-3 have equations which differ from Step 1 since it must account for replicated bins and cannot solve for *µ*_*n*_ analytically. The probability of each bin being replicated (*ϕ*_*n,m*_) is a function of the cell’s time in S-phase (*τ*_*n*_), the locus’s replication timing (*ρ*_*m*_), and the replication stochasticity term (*α*). Replication stochasticity (*α*) controls how closely cells follow the global RT profile by adjusting the temperature of a sigmoid function. The following equation corresponds to function block *g*:

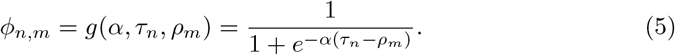

Equations corresponding to function block *f* differ from those in Step 1. The total CN (*χ*_*n,m*_) is double the somatic CN (*X*_*n,m*_) when a bin is replicated (*Y*_*n,m*_ = 1),

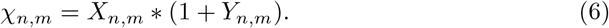

The GC rates (*ω*_*n,m*_) and negative binomial event counts (*δ*_*n,m*_) are computed the same as in Step 1 (Eq 2, Eq 4). However, the expected read count uses total instead of somatic CN,

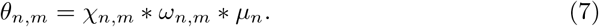

Since CN is learned in Steps 2-3, the coverage/ploidy scaling term (*µ*_*n*_) must also be learned. We use a normal prior 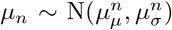 where the approximate total ploidy and total read counts are used to estimate the mean hyperparameters 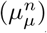. Total ploidies for each cell are approximated using the CN prior concentrations (*η*) and times within S-phase (*τ*) to account for both somatic and replicated copies of DNA that are present. We fixed the standard deviation hyperparameters 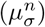 to always be 10x smaller than the means to ensure that *µ*_*n*_ *≥* 0 despite use of a normal distribution (used for computational expediency),

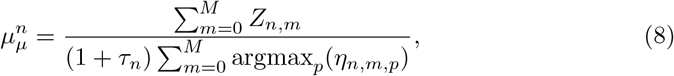

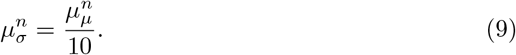

### Constructing the CN prior concentrations

There are two ways to construct the CN prior concentrations within PERT. The first is to use the most similar high-confidence G1/2 clone to define the concentrations for each unknown cell (clone method). We assign each unknown cell its clone (*c*_*n*_) via Pearson correlation between the cell read depth profile (*Z*_*n*_) and the clone pseudobulk read depth profile (*Z*_*c*_),

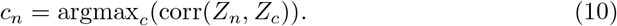

Clone pseudobulk CN and read depth profiles represent the median profile across all high-confidence G1/2 cells in a given clone *c*. Once we have clone assignments for each unknown cell, the CN concentration of all possible states *P* at each genomic bin (*η*_*n,m,p*_) is constructed to be *w* times larger for the state *p* that matches the clone pseudobulk CN state 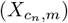 for that same bin compared to all other states. The default setting is *w* = 10^6^:

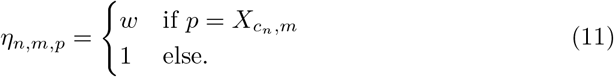

The second way to construct the prior is to leverage additional information from the most similar high-confidence G1/2 cells when constructing *η*_*n,m,p*_ (composite method). The rationale for the composite method is that there might be rare CNAs within a clone which only appear in a handful of cells but do not appear in the clone pseudobulk CN profile *X*_*c*_. To find the most similar high-confidence G1/2 cells, we compute the read depth correlation between the unknown cell 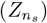 and the high-confidence G1/2 cells from the best matching clone 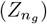,

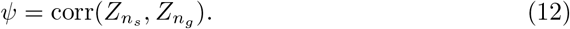

The consensus clone CN profile and top *J* matches for each unknown cell are then used to construct the CN prior (*η*_*n,m,p*_). Each row of *ψ* is sorted to obtain the top *J* highconfidence G1/2 matches *n*_*g*_(0), .., *n*_*g*_(*J*–1). All entries are initialized to 1 (*η*_*n,m,k*_ = 1) before adding varying levels of weight (*w*) to states where the CN matches a G1/2-phase cell or clone pseudobulk CN profile. The default settings are *w* = 10^5^ and *J* = 5:

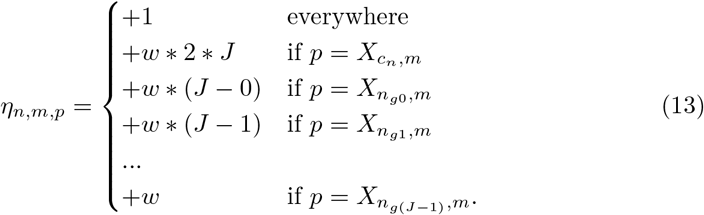

By default, the composite method is used during Step 2 and the clone method is used during Step 3; however, the user may select between both methods during Step 2. Using the clone method during Step 2 should be seen as a ‘vanilla’ version of PERT which should be used when very few cell-specific CNAs are present. The clone method is used for Step 3 since the composite method would produce many self-matching cells. A comparison of the two methods can be seen when benchmarking PERT on simulated data (Supplementary Note).

### Model initialization and hyperparameters

Splitting cells into initial sets of high-confidence G1/2-phase and unknown cells is performed by thresholding heuristic per-cell features known to correlate with cell cycle phase. PERT uses clone-normalized number of input CN breakpoints between neighboring genomic bins (BKnorm) and clone-normalized median absolute deviation in read depth between neighboring genomic bins (MADNnorm). These features are referred to as ‘HMMcopy breakpoints’ and ‘MADN RPM’, respectively, in the main text and figures. Note that breakpoints between chromosome boundaries are not counted.

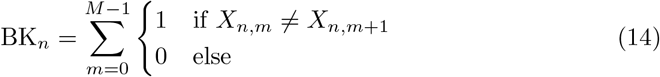

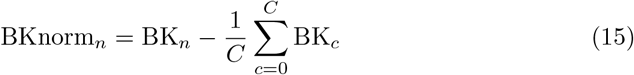

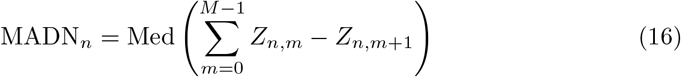

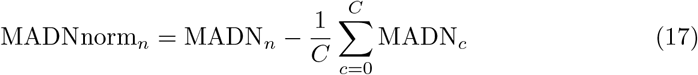

Under default settings, PERT initializes cells with MADNnorm*<*0 and BKnorm*<*0 as high-confidence G1/2-phase with all other cells as unknown phase. Initial cell phases can also be input by users based on experimental measurements or alternative metrics such as 10X CellRanger-DNA’s ‘dimapd’ score (used in [17, 23, 24]), the Laks *et al* classifiers’ S-phase probability and quality scores [16], or read depth correlation with a reference RT profile [25].

To improve convergence speed, each cell’s time in S-phase (*τ*_*n*_) is initialized using scRT results from a clone-aware adaptation of Dileep *et al* [21] which thresholds the clone-normalized read depth profiles into replicated and unreplicated bins. Each unknown cell *n* is assigned to clone *c* with the highest correlation between cell and clone pseudobulk read depth profiles (Eq 10). The read depth of each cell is then normalized by the CN state with highest probability within the CN prior (*η*_*n,m,p*_),

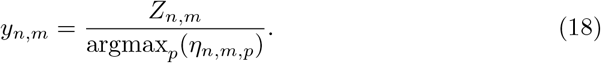

The clone-normalized read depth profiles (*y*_*n*_) are then binarized into replication state profiles (*Y*_*n*_) using a per-cell threshold (*t*_*n*_ ∈ [0, 1]) that minimizes the Manhattan distance between the real data and its binarized counterpart.

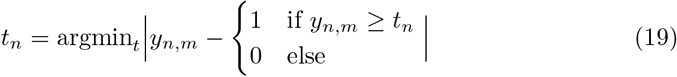

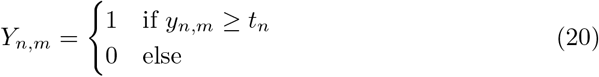

The fraction of replicated bins per cell from the deterministic replication states *Y*_*n,m*_ are then used to initialize the parameter representing each cell’s time in S-phase (*τ*_*n*_) within PERT’s probabilistic model.

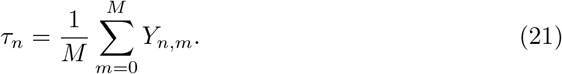

Initialization of *τ*_*n*_ is particularly important because the model might mistake an early S-phase cell (*<*20% replicated) for a late S-phase cell (*>*80% replicated), or vice versa, as both have relatively ‘flat’ read depth profiles compared to mid-S-phase cells. Thus *τ*_*n*_ will rarely traverse mid-S-phase values during inference when its initial and true values lie far apart. Additional parameter initializations include *λ* = 0.5 for negative binomial overdispersion and *β*_*σ,k*_ = 10^*-k*^ for the standard deviation of each GC bias polynomial coefficient *k*. Unlike *τ*_*n*_, the model is unlikely to get stuck at local minima with these parameters so they are initialized to the same values globally.

The latent variables *β*_*µ*_, *ρ*, and *α* are sampled from prior distributions with fixed hyperparameters. The mean of all GC bias polynomial coefficients (*β*_*µ*_) are drawn from the prior N(0, 1). Each locus’s replication timing (*ρ*) is drawn from the prior Beta(1, 1) to create a uniform distribution on the domain [0, 1]. The replication stochasticity parameter (*α*) is drawn from the prior distribution Γ(shape = 2, rate = 0.2) which has a mean of 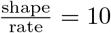 = 10 and penalizes extreme values on a positive real domain.

### PERT phase predictions

We used the PERT model output to predict ‘G1/2’, ‘S’, and ‘low quality’ (LQ) phases for each cell. G1/2-phase cells were defined by having *<*5% or *>*95% replicated bins. Of the remaining cells with 5-95% replicated bins, those with high read depth autocorrelation (*>*0.5), replication state autocorrelation (*>*0.2), or fraction of homozygous deletions (*X* = 0, *>*0.05) were deemed to be low quality. All other cells were deemed to be in S-phase. Using 500kb bins, autocorrelation scores were the average of all autocorrelations ranging from 10 to 50 bin lag size. Thresholds used for splitting S and LQ phases can be adjusted by users should the default settings produce unexpected output.

### Model construction and inference

PERT is written using Pyro which is a probabilistic programming language written in Python and supported by PyTorch backend [37]. PERT uses Pyro’s implementation of Black Box Variational Inference (BBVI) which enables the use of biologically-informed priors instead of being limited to conjugate priors [60]. Specifically, we use the AutoDelta function which uses a Taylor approximation around the *maximum a posteriori* (MAP) to approximate the posterior. Optimization is performed using the Adam optimizer. By default, we set a learning rate of 0.05 and convergence is determined when the relative change in the evidence lower bound (ELBO) is *<* 10^*-*6^ or the maximum number of iterations (2000 for step 2, 1000 for steps 1 and 3) is reached.

### Simulated datasets

To benchmark PERT’s ability to accurately infer single-cell replication states, somatic CN states, and cell cycle phase against Kronos and the Laks *et al* cell cycle classifier, we simulated datasets with varying clonal structures and cell-specific CNA rates. Somatic CN states are simulated by first drawing clone CN profiles and then drawing cellspecific CNAs that deviated from said clone CN profile. All CNAs are drawn at the chromosome-arm level. 400 S- and 400 G1/2-phase cells are simulated in each dataset.

Once CN states have been simulated, we simulate the read depth using PERT as a generative model. We condition the model on the provided *β*_*µ*_, *β*_*σ*_, *λ, α, ρ, γ*, and *X* parameters when generating cell read depth profiles. All read depth values (*Z*) are in units of reads per million. RepliSeq data for various ENCODE cell lines are used to set *ρ* values for each clone [38]. G1/2-phase cells were conditioned to have all bins as unreplicated *Y* = 0. S-phase cells had their cell cycle times *τ* sampled from a Uniform(0, 1) distribution. A table of all the parameters used in each simulated dataset can be found in Supplementary Table 1.

We called CN on simulated binned read count data using HMMcopy. Given that Kronos was designed as an end-to-end pipeline that takes in raw BAM files, we forked off the Kronos repository and edited their ‘Kronos RT’ module to accept binned read count and CN states as input. Cells were split into S- and G1/2-phase Kronos input populations according to their true phase. Code to our forked repository can be found at https://github.com/adamcweiner/Kronos scRT. Similarly, we removed features from the Laks *et al* cell cycle classifier that used alignment information such as the percentage of overalapping reads per cell. The Laks classifier was retrained with said features removed prior to deployment on simulated data (Supplementary Fig. 1).

### Experimental methods

Detailed descriptions of the data generation methods are described in Laks *et al*, Funnell *et al*, and Salehi *et al* [16, 40, 45]. Such descriptions include generation of the cell cycle FACS datasets, generation of engineered hTERT cell lines, xenografting, time series passaging, and scWGS with direct library preparation (DLP+) sequencing.

### scWGS data processing

Unless otherwise noted, all scWGS data was generated via DLP+. All DLP+ data was passed through https://github.com/shahcompbio/single cell pipeline before downstream analysis. This pipeline aligned reads to the hg19 reference genome using BWA-MEM. Each cell was then passed through HMMcopy using default arguments for single-cell sequencing. HMMcopy’s output provided read count and gc-corrected integer CN states for each 500kb genomic bin across all cells and loci. Loci with low mappability (*<*0.95) and cells with low read count (*<*500,000 reads) were removed. Cells were also filtered for contamination using the FastQ Screen which tags reads as matching human, mouse, or salmon reference genomes. If *>*5% of reads in a cell are tagged as non-human the cell is flagged as contaminated and subsequently removed.

Cells were only passed into phylogenetic trees if they were called as G1/2-phase and high quality by classifiers described in Laks *et al* [16]. In certain cases, cells might be manually excluded from the phylogenetic tree if they pass the cell cycle and quality filters but have an abnormally high number of HMMcopy breakpoints. All cells included in the phylogenetic tree are initialized in PERT as the set of high-confidence G1/2 cells; all cells outside the tree are initialized as unknown cells.

### Phylogenetic clustering based on CN profiles

We used the clone IDs from Funnell *et al* for high-confidence G1/2 cells [40]. These single-cell phylogenetic trees were generated using sitka [58]. Sitka uses CN breakpoints (also referred to as changepoints) across the genome as binary input characters to construct the evolutionary relationships between cells. Sitka was run for 3,000 chains and a consensus tree was computed for downstream analysis. The consensus tree was then cut at an optimized height to assign all cells into clones (clusters). For datasets with no sitka trees provided or select datasets, cells were clustered into clones using K-means where the number of clones was selected through Akaike information criterion. We performed a K-means reclustering for the Salehi *et al* TNBC PDX data [45] as sitka produced small clusters which inhibited robust tracking of S-phase clone fractions across multiple timepoints.

### Pseudobulk profiles

Many times in the text we describe “pseudobulk” replication timing, copy number, or read depth profiles within a subset of interest (i.e. cells belonging to the same clone or sample). To compute pseudobulk profiles, we group all the cells of interest and then take the median values for all loci in the genome. When computing pseudobulk CN profiles, we only include the cells of the modal (most common) ploidy state before computing median values for all loci.

### S-phase times

When we refer to the “time” of individual S-phase cells, such a time is calculated as the fraction of replicated bins per cell. Thus, S-phase times near 1 are in late S-phase cells and S-phase times near 0 are early S-phase cells.

### Comparison of RT profiles to Hi-C A/B compartments

Hi-C compartment data were downloaded from ENCODE for T47D and B-lymphoblast (GM-prefix) cell lines using the accession codes ENCFF713FCA, ENCFF220LEI, ENCFF733ZUJ, ENCFF907MWF, ENCFF522SPQ, and ENCFF4-11JKH [38]. Genomic coordinates were lifted to human reference hg19 for comparison. Due to varying quality and sequencing platforms of each Hi-C library, we used Spearman instead of Pearson correlation.

### Bespoke factor model which learns feature importance and RT profiles directly from clone RT profiles

We built a multivariate regression model which learned importance terms and RT profiles for each feature directly from the matrix of clone RT profiles. This model has the following terms and equations:

*RT*_*c,m*_: the observed replication timing of clone *c* at locus *m* on the domain of [0,1]. This represents the fraction of replicated bins at locus *m* across all S-phase cells *n* in clone *c*.

*ρ*_*k,m*_: the latent replication timing of feature *k* at locus *m*.

*I*_*c,k*_: indicator mask representing which features *k* are present for clone *c*.

*β*_*k*_: importance term for feature *k*.

*σ*: standard deviation term when going from expected to observed replication timing; sampled from a uniform distribution on the domain (0,1).

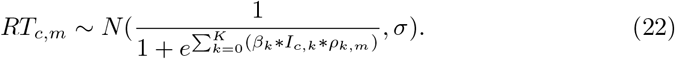

All latent replication timing terms *ρ*_*k*_ are normalized to have mean of 0 and variance of 1 and there is only *β* value per class of features

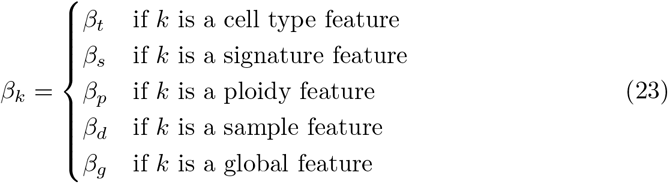

This model is implemented in pyro and fit using BBVI [37, 60]. We use the AutoNormal function which uses Normal distributions to approximate the posterior. Optimization is performed using the Adam optimizer with a learning rate of 0.02. Convergence is determined when the relative change in ELBO is *<* 10^*-*3^ of the total ELBO change between first and current iteration.

### Using SIGNALS to quantify allelic ratios from scDNA- and scRNA-seq

In brief, SIGNALS uses haplotype blocks genotyped in single cells and implements an hidden Markov model (HMM) based on a Beta-Binomial likelihood to infer the most probable haplotype-specific state. SHAPEIT was used to generate the haplotype blocks for SIGNALS input [61]. A full description of SIGNALS can be found in Funnell *et al* [40]. Within each haplotype block for each sample, the major (most common) allele is labeled as the A-allele with the minor (less common) allele labeled as the B-allele. The B-allele frequency (BAF) is computed as the fraction of B-allele heterozygouos single nucleotide polymorphisms (SNPs) out of all heterozygous SNPs present in a given bin. SIGNALS is run on scDNA data by default but when scRNA data is also available, the haplotype blocks derived from the scDNA data can be used to extract A- and B-allele counts in the scRNA data too (albeit with much fewer counts as there are fewer SNPs sequenced in scRNA data).

### Gastric cancer cell line data

10X Chromium single-cell DNA (10X scWGS) data of gastric cancer cell lines NCI-N87, HGC-27, and SNU-668 were downloaded from SRA (PRJNA498809). Copy number calling was performed using the CellRanger-DNA pipeline using default parameters. Data was aggregated from 20kb to 500kb bins for analysis with PERT. Each cell line’s doubling time and fraction of G1-phase scRNA cells were extracted from Andor *et al* [17].

### MSK SPECTRUM data

We obtained matched scRNA and scWGS from HGSOC patient OV-081 from the MSK SPECTRUM cohort. Samples were collected under Memorial Sloan Kettering Cancer Center’s institutional IRB protocol 15-200 and 06-107. Single cell suspensions from surgically excised tissues were generated and flow sorted on CD45 to separate the immune component as previously described. CD45 negative fractions were then sequenced using the DLP+ platform as previously described. Detailed generation of scRNA data can be found in [47].

### Clone S-phase enrichment scores

To test whether a clone (*c*) is significantly enriched or depleted for S-phase cells at a given timepoint (*t*), we must compare that clone’s fraction in both S- and G1/2-phases. We first define the following variables as such:

*N*_*s,c,t*_: Number of S-phase cells belonging to clone *c* at time *t*

*N*_*g,c,t*_: Number of G1/2-phase cells belonging to clone *c* at time *t*

*N*_*s,t*_: Total number of S-phase cells across all clones at time *t*

*N*_*g,t*_: Total number of G1/2-phase cells across all clones at time *t*

*N*_*t*_: Total number of cells in a population at time *t* (all clones, all phases)

We can then define the fractions of S- and G1/2-phase cells assigned to clone *c* at time *t* (*f*_*s,c,t*_, *f*_*g,c,t*_):

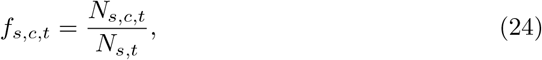

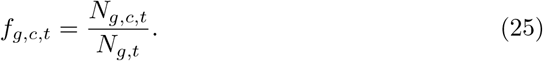

Each clone’s continuous S-phase enrichment (SPE) score (*ξ*_*c,t*_) is the difference between the S- and G1/2-phase fractions. Positive values indicate the clone is enriched for S-phase cells,

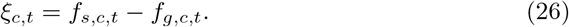

Using the fraction of G1/2-cells belonging to clone *c*, we can compute the expected total number of cells in clone *c* and time *t* across all cell cycle phases,

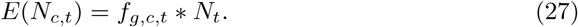

We produce a p-value for enrichment of S-phase cells using a hypergeometric test scipy.stats.hypergeom(M=*N*_*t*_, n=*N*_*s,t*_, N=*E*(*N*_*c,t*_)).sf(*N*_*s,c,t*_). To produce a p-value for S-phase depletion we subtract this enrichment p-value from 1. All p-values are Bonferroni-corrected by dividing by the total number of statistical tests. p-adjusted thresholds of 10^*-*2^ are used for saying a clone is significantly enriched or depleted for S-phase cells within a given library.

### Clone expansion scores

For time-series scWGS experiments, we computed clone expansion scores for each clone *c* at time *t* (*S*_*c,t*_) by examining the fraction of G1/2-phase cells belonging to clone *c* at timepoint *t* (*f*_*g,c,t*_) and the subsequent timepoint (*f*_*g,c,t*+1_). Positive values indicate the clone expands by the next timepoint,

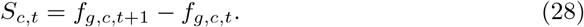

### Comparing SPE to expansion in treated vs untreated data

To test that treated clones had a significant difference in their relationship between SPE scores (*ξ*_*c,t*_) and expansion scores (*S*_*c,t*_) in treated (*T*) vs untreated (*U*) data, we first fit a linear regression curve to the untreated data,

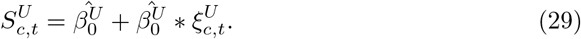

We then computed the residuals between the treated data and this line of best fit,

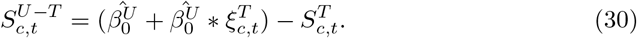

We then computed a second linear regression curve to the residuals 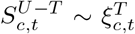 and computed the p-value for a hypothesis test whose null hypothesis is that the slope is zero, using Wald Test with t-distribution of the test statistic. Having a *p <* 0.05 indicated that the slope of the treated and untreated lines are significantly different. All clone and time points with *<* 10 G1/2-phase cells were excluded from such analysis.

### Cell cycle analysis of scRNA data

When available, we validated PERT cell cycle distributions using the cell cycle distributions estimated through scRNA sequencing. We determined the cell cycle phase of each scRNA cell using the Seurat CellCycleScoring() function [62] which uses a set of S- and G2M-phase markers derived from Tirosh *et al* [36].

### Statistical tests

When boxplots are presented in the figures, hinges represent the 25% and 75% quantiles and whiskers represent the *±* 1.5x interquartile range. Statistical significance is tested using independent t-tests from scipy.stats unless otherwise noted. Bonferroni correction is implemented for all statistical tests to limit false discovery. The number of stars is a shorthand for the adjusted p-value of a given statistical test (*<* 10^*-*4^: ****, *<* 10^*-*3^: ***, *<* 10^*-*2^: **, *<* 0.05: *, *≥* 0.05: ns). Shaded areas surrounding linear regression lines of best fit represent 95% confidence intervals obtained via boostrapping (n=1000 boostrap resamples). Unless otherwise noted, linear regressions are annotated with Pearson correlation coefficients (*r*) amd the p-value for a hypothesis test whose null hypothesis is that the slope is zero, using the Wald Test with t-distribution of the test statistic.

## 5 Declarations

### Ethics approval and consent to participate

Not applicable

### Competing interests

S.P.S. and S.A. are shareholders of Imagia Canexia Health Inc. S.P.S. has an advisory role to AstraZeneca Inc. All relationships are outside the scope of this work.

### Funding

This project was generously supported by the Cycle for Survival, by the Marie-Josée and Henry R. Kravis Center for Molecular Oncology and the NCI Cancer Center Core grant P30-CA008748 supporting Memorial Sloan Kettering Cancer Center. S.P.S. holds the Nicholls Biondi Chair in Computational Oncology and is a Susan G. Komen Scholar (#GC233085). This work was also funded in part by awards to S.P.S.: the Cancer Research UK Grand Challenge Program (GC-243330), and an NIH RM1 award (RM1-HG011014). A.C.W. is supported by NCI Ruth L. Kirschstein National Research Service Award for Predoctoral Fellows F31-CA271673. M.J.W. is supported by NCI Pathway to Independence award K99-CA256508. S.S. is supported by NCI Pathway to Independence award K99-CA277562 I.V.-G. is supported by a Mentored Investigator Award from the Ovarian Cancer Research Alliance (650687).

### Data availability

Data from Funnell *et al* [40] can be found on zenodo https://zenodo.org/record/6998936#.Y0h3luzMLzc. Raw scWGS data from Laks *et al* and Salehi *et al* [16, 45] are available from the European Genome-Phenome under study IDs EGAS00001004448 and EGAS00001003190, respectively. Raw scRNA data for SPECTRUM patient OV-081 is available at https://www.synapse.org/msk_spectrum. All other data will be made available for controlled access upon final publication.

### Code availability

The following code repositories are publicly available and contain tutorials for installation and use.

- Package containing PERT model and other tools for scRT analysis: https://github.com/shahcompbio/scdna_replication_tools
- DLP+ single-cell whole genome sequencing pipeline: https://github.com/shahcompbio/single_cell_pipeline

The following repositories will be made available upon final publication.

- Analysis scripts and figure generation: https://github.com/shahcompbio/scdna_replication_paper
- LaTeX files and figures for manuscript generation: https://github.com/adamcweiner/pert_manuscript

## Authors’ contributions

A.C.W., S.P.S., and A.M. designed the methodology and wrote the manuscript. N.R. helped with writing and editing. All authors helped to design experiments and/or analyze the data.

## Authors’ information

S.P.S. and A.M. jointly supervised this work.

## Extended Data Figures

**Fig. ED1.**
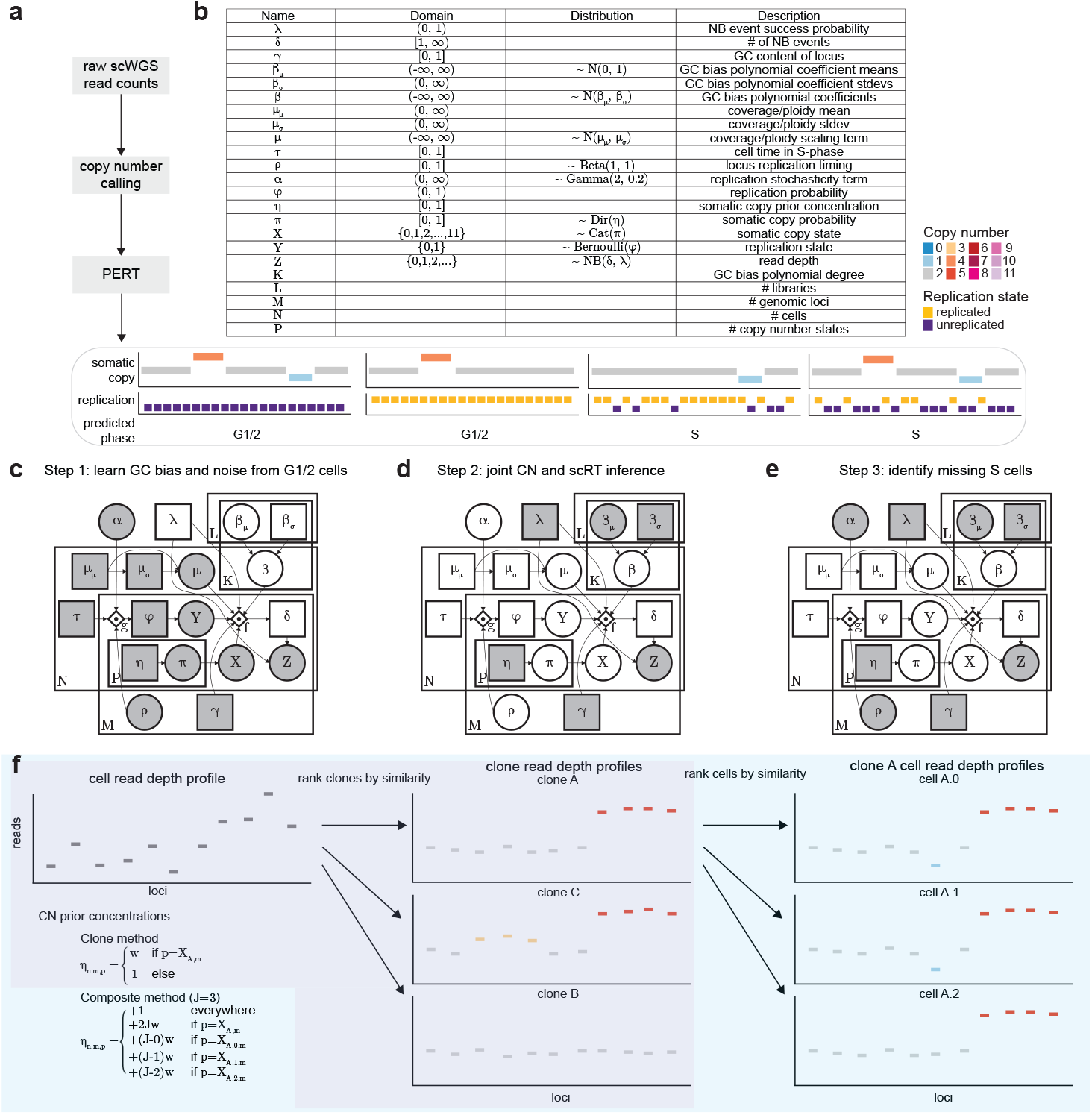
Details of PERT inference. **a)** PERT takes scWGS binned read count and CN calls as input and learns somatic copy number, replication states, and cell cycle phase predictions for all cells. **b)** Table of all parameters, domains and distributions used in PERT. **c-e)** Full graphical model for 3-step learning procedure. **c)** PERT first learns overdispersion (*λ*) and GC parameters (*β*_*µ*_, *β*_*σ*_) from high-confidence G1/2 cells where we condition all bins as unreplicated (*Y* = 0) and CN states (*X*) according to CN caller results. **d)** PERT conditions the parameters learned in Step 1 to learn latent replication and somatic CN states in unknown cells. **e)** Replication timing (*ρ*) and stochasticity (*α*) terms learned in Step 2 are conditioned as Step 3 learns latent replication and somatic CN states in high-confidence G1/2-phase cells to search for any missing S-phase cells. **f)** Overview of clone and composite methods to set copy number prior concentrations (*η*). Composite method is used by default. Pearson correlation is used to determine similarity.

**Fig. ED2.**
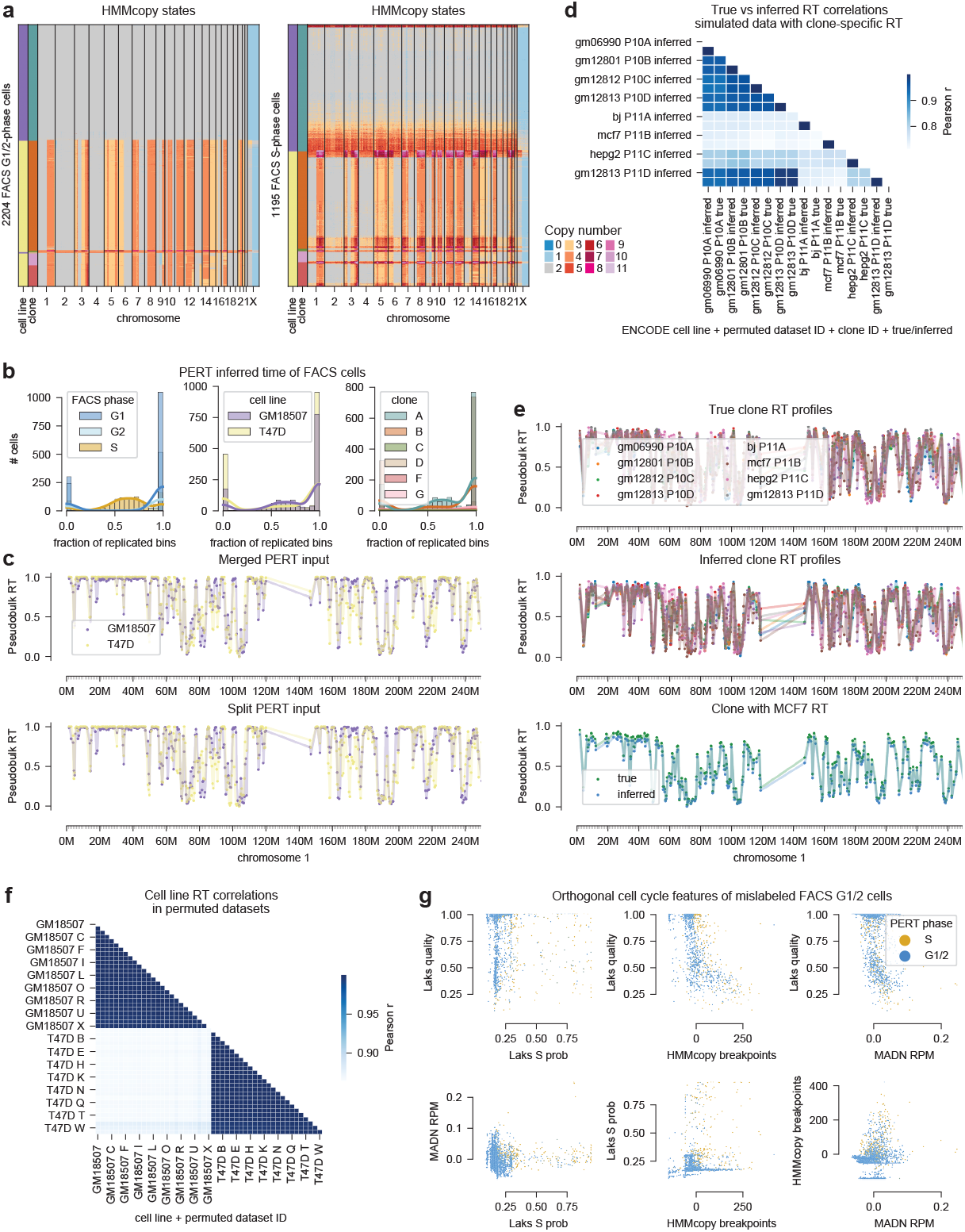
Assessing PERT’s robustness to clone-specific RT profiles and poor phase initialization. **a)** HMMcopy states for GM18507 and T47D cell lines FACS isolated into G1/2-(left) and S-phase (right) populations. **b)** Histogram of inferred fraction of replicated bins for FACS cell lines where colors represent FACS phase (left), cell line (middle), and clone ID (right). **c)** Inferred cell line RT profiles for chromosome 1 when the two cell lines are merged into one sample (top) or split into separate samples (bottom) before passing into PERT. **d)** Pairwise Pearson correlation between true and inferred clone RT profiles in two simulated datasets (P10 and P11) where each clone has a unique CN profile and unique ENCODE cell line RT profile. Cell lines with the ‘GM’ prefix are derived from B-lymphocytes, BJ from human foreskin fibroblasts, MCF7 from breast cancer, and HepG2 from liver cancer. Rows and columns are sorted in the same order. **e)** True (top) and inferred (middle) clone RT profiles across chr1 for simulated datasets with clone-specific RT. True and inferred RT of the clone emulating the ENCODE MCF7 RT is shown at the bottom. **f)** Pairwise Pearson correlation between inferred cell line RT profiles across all permuted datasets. Datasets A-C have the lowest permutation rate (0.01); U-W have the highest permutation rate (0.75). **g)** Pairwise scatterplots of orthogonal cell cycle phase features for FACS=G1/2 cells mislabeled as S-phase. Cells are colored by their predicted PERT phase. MADN RPM: median absolute deviation between neighboring bins of reads per million, normalized to 0 within each clone (Methods). Laks S prob: S-phase probability according to the Laks cell cycle classifier. Laks quality: Probability of a cell being high quality according to the Laks cell quality classifier. HMMcopy breakpoints: the number of adjacent bins per cell that do not share the same HMMcopy state, normalized to 0 within ea3ch4 clone (Methods).

**Fig. ED3.**
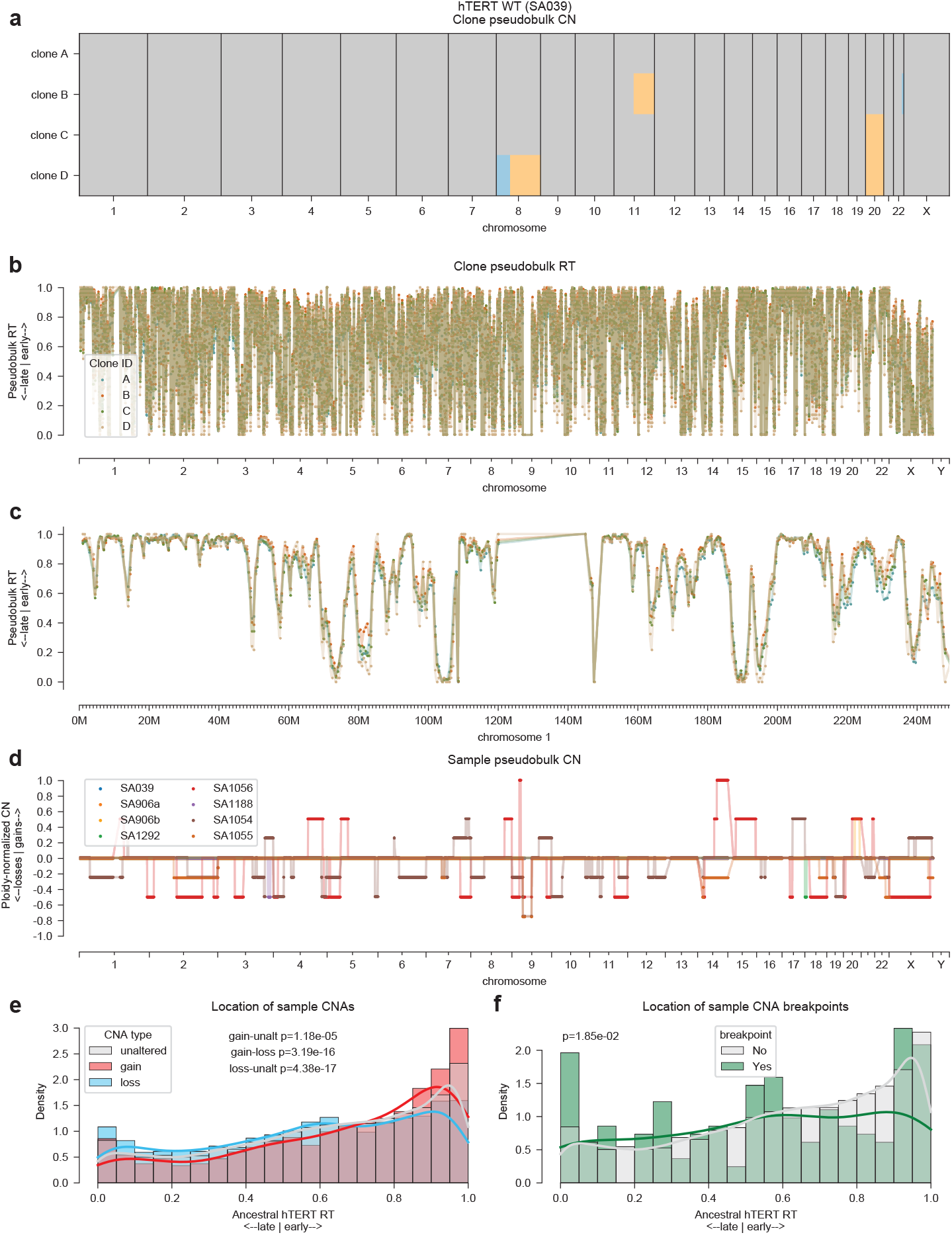
PERT identifies RT profiles of ancestral WT clone prior to emergence of CNAs. **a-c)** Clone CN and RT profiles for hTERT WT sample SA039. **d)** CN profiles for all hTERT clones, normalized by ploidy. Values *>* 0 are gains, *<* 0 are losses, and = 0 are unaltered. Distribution of hTERT WT SA039 clone A RT values split by whether a locus contains a clonal CNA breakpoint across all hTERT samples. **e-f)** Distribution of hTERT SA039 clone A (diploid WT) RT values split by **e)** sample pseudobulk CNA types and **f)** the presence of sample pseudobulk CNA breakpoints.

**Fig. ED4.**
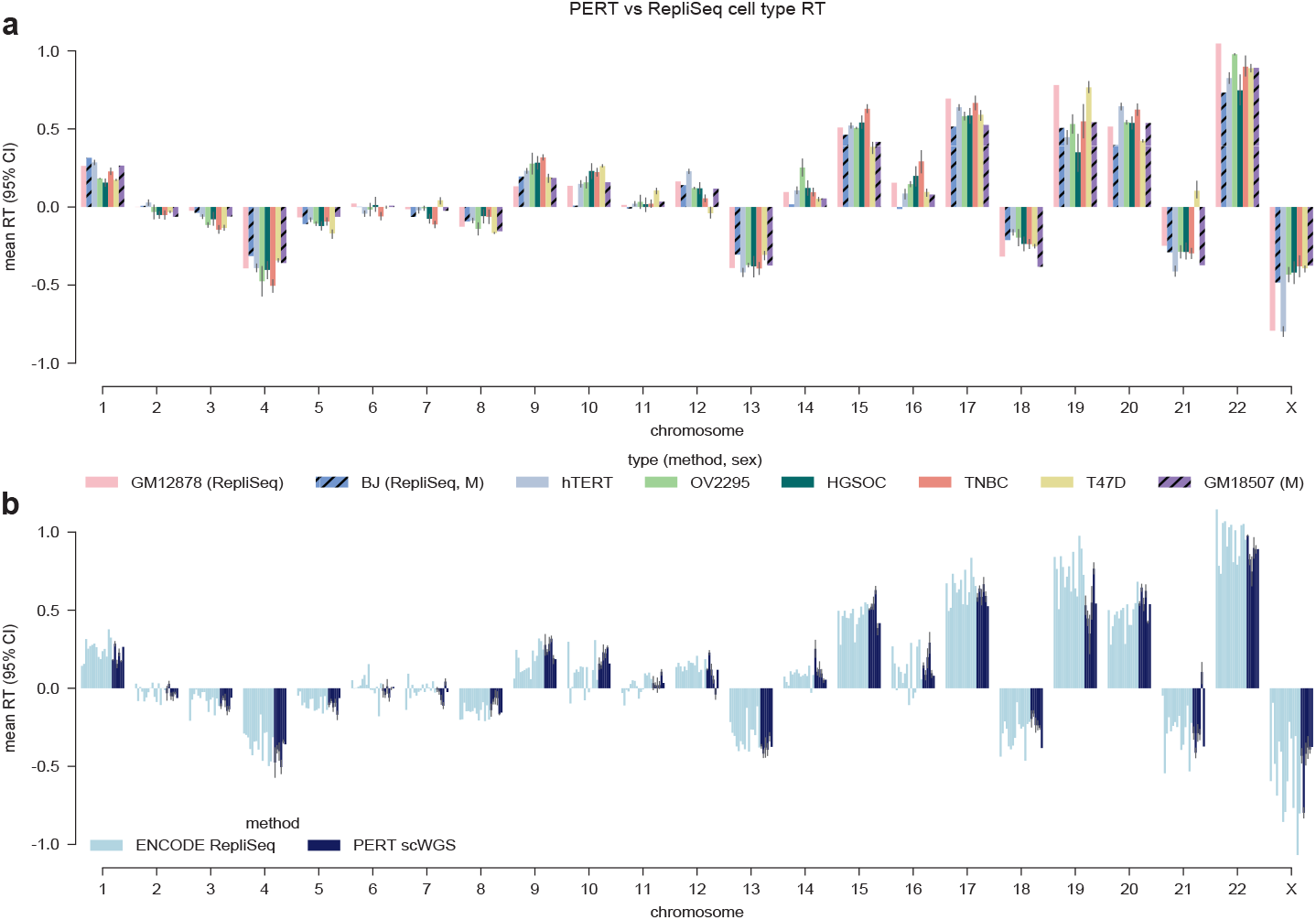
Per-chromosome cell type RT profiles of PERT vs RepliSeq data. Mean RT across cell types and chromosomes. Error bars represent the 95% confidence intervals over the per-chromosome mean RT when multiple clones are present. **a)** Cell types shown in Fig. 3f with colors corresponding to cell type. **b)** Full set of PERT and RepliSeq cell types where each cell type is colored by the method from which the RT profile was obtained. The full set of ENCODE RepliSeq cell types (in order) are: MCF7, BG02ES, BJ, GM06990, GM12801, GM12812, GM12813, GM12878, HELAS3, HEPG2, HUVEC, IMR90, K562, SKNSH, NHEK. The full set of PERT cell types match those seen in panel **a**.

**Fig. ED5.**
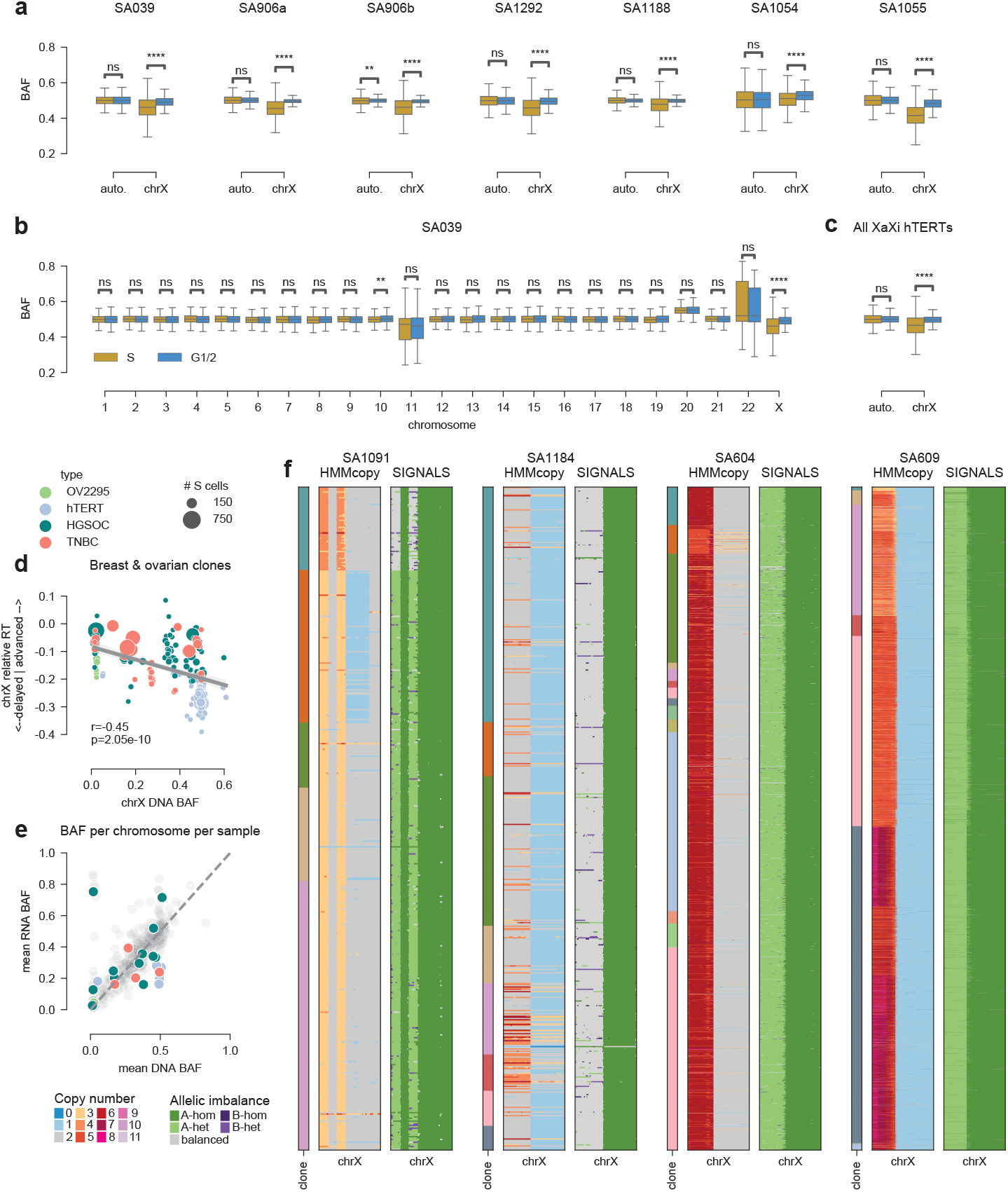
Chromosome X replication timing shifts reflect X-inactivation and reactivation status. **a-c)** B-allele frequencies of S-phase vs G1/2-phase cells (determined via PERT) across the genome for hTERT samples. A/B haplotype block labels are identical across all hTERT samples. **a)** Per-sample comparison of autosomes vs chrX. **b)** Per-chromosome comparison for sample SA039. **c)** Aggregate comparison of autosomes vs chrX for all hTERT samples with XaXi genotype. **d)** chrX B-allele frequency vs relative RT for all clones in the metacohort with *>* 10 S-phase cells. **e)** Mean DNA vs RNA BAF per chromosome per sample for breast and ovarian samples in the metacohort. All chrX points are colored by their sample type. All autosomes arms are colored light grey. The dashed y=x line illustrates 1:1 relationship between gene dosage and transcription. **f)** HMMcopy CN and SIGNALS allelic imbalance states in chrX for the four samples with Xq LOH but not Xp LOH. Clone IDs are annotated on the left-hand side of each sample.

**Fig. ED6.**
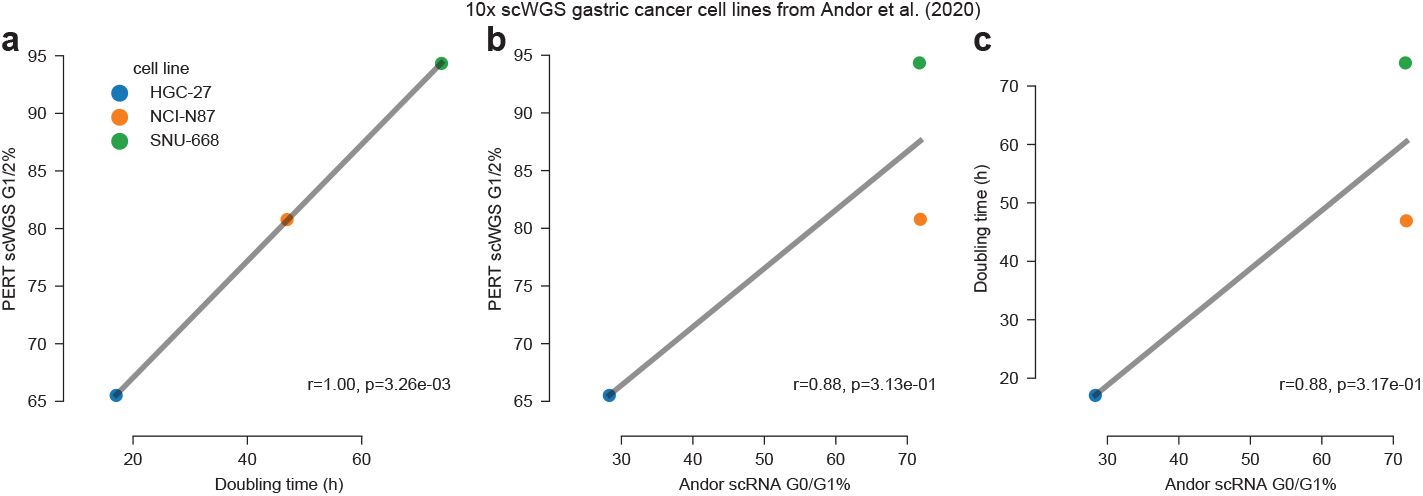
Relationship between cell cycle fractions and doubling time in gastric cancer cell lines sequenced with 10X scWGS platform. PERT-derived fraction of G1/2-phase cells in 10X scWGS libraries of each cell line compared to the corresponding **a)** doubling time (hours) and **b)** fraction of G0/1-phase cells in the 10X scRNA libraries. **c)** Comparison of scRNA G0/1-phase cells to doubling time. Data was derived from Andor et al 2020 [17].

**Fig. ED7.**
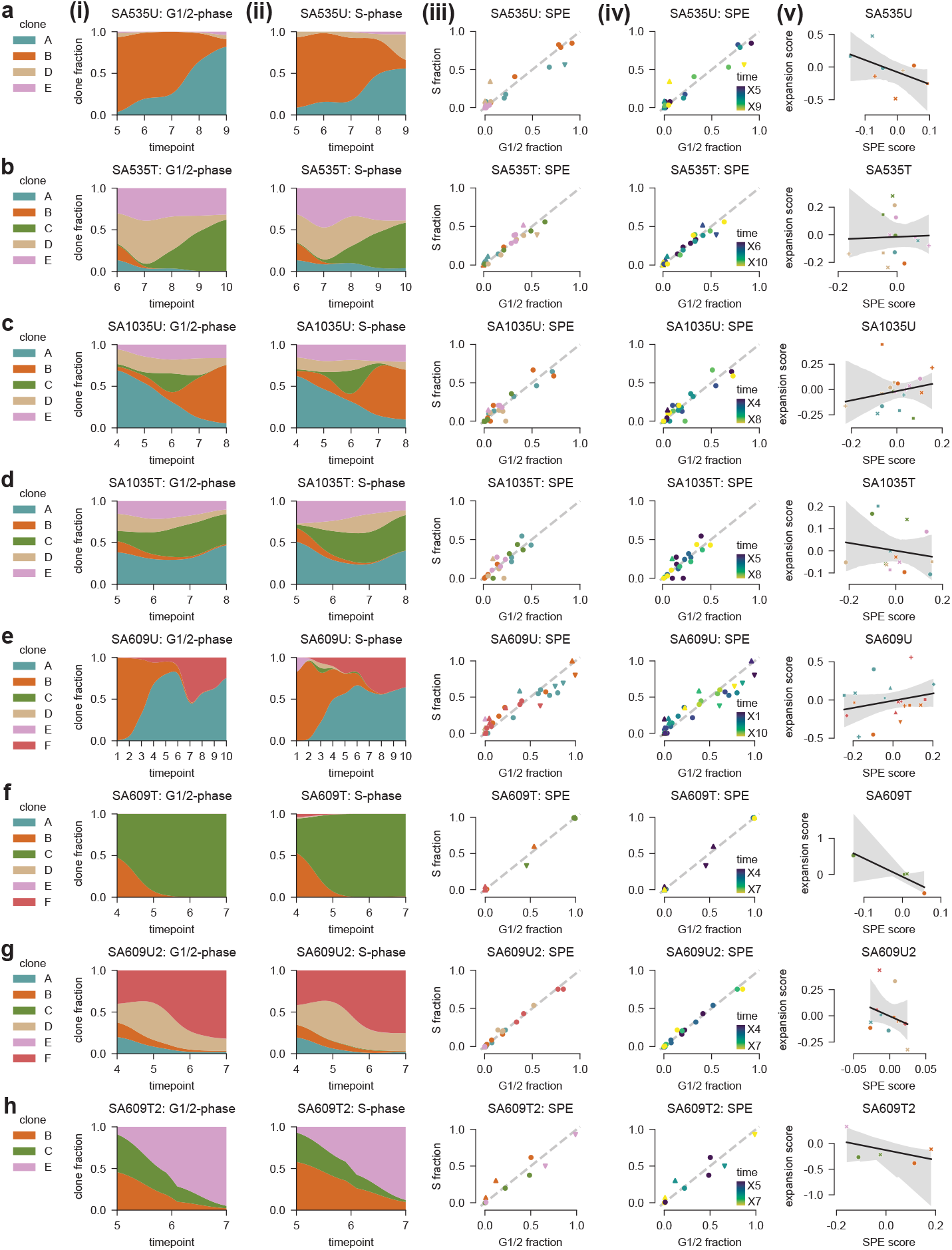
Clone cell cycle phase enrichment and fitness across all TNBC PDX samples. **i)** Relative fraction of each clone within G1/2- and **ii)** S-phase populations. **iii-iv)** Comparison of each clone’s fraction in S-vs G1/2-phase populations at each timepoint. Dashed gray line represents equal prevalence in both phases. Triangles denote clone and timepoint combinations significantly (*p*_*adj*_ *<* 0.01) enriched or depleted for S-phase cells via hypergeometric test. Distance from the dashed gray line represents each point’s continuous SPE score. **v)** Relationship between SPE and clone expansion between timepoints *t* and *t* + 1 for each clone and timepoint combination with *>* 10 G1/2-phase cells. Lines represent linear regression fits with shaded areas representing 95% confidence intervals. Point colors represent the clone ID and the shapes represent the timepoint. **a-h)** Each row corresponds to a unique sample.

**Fig. ED8.**
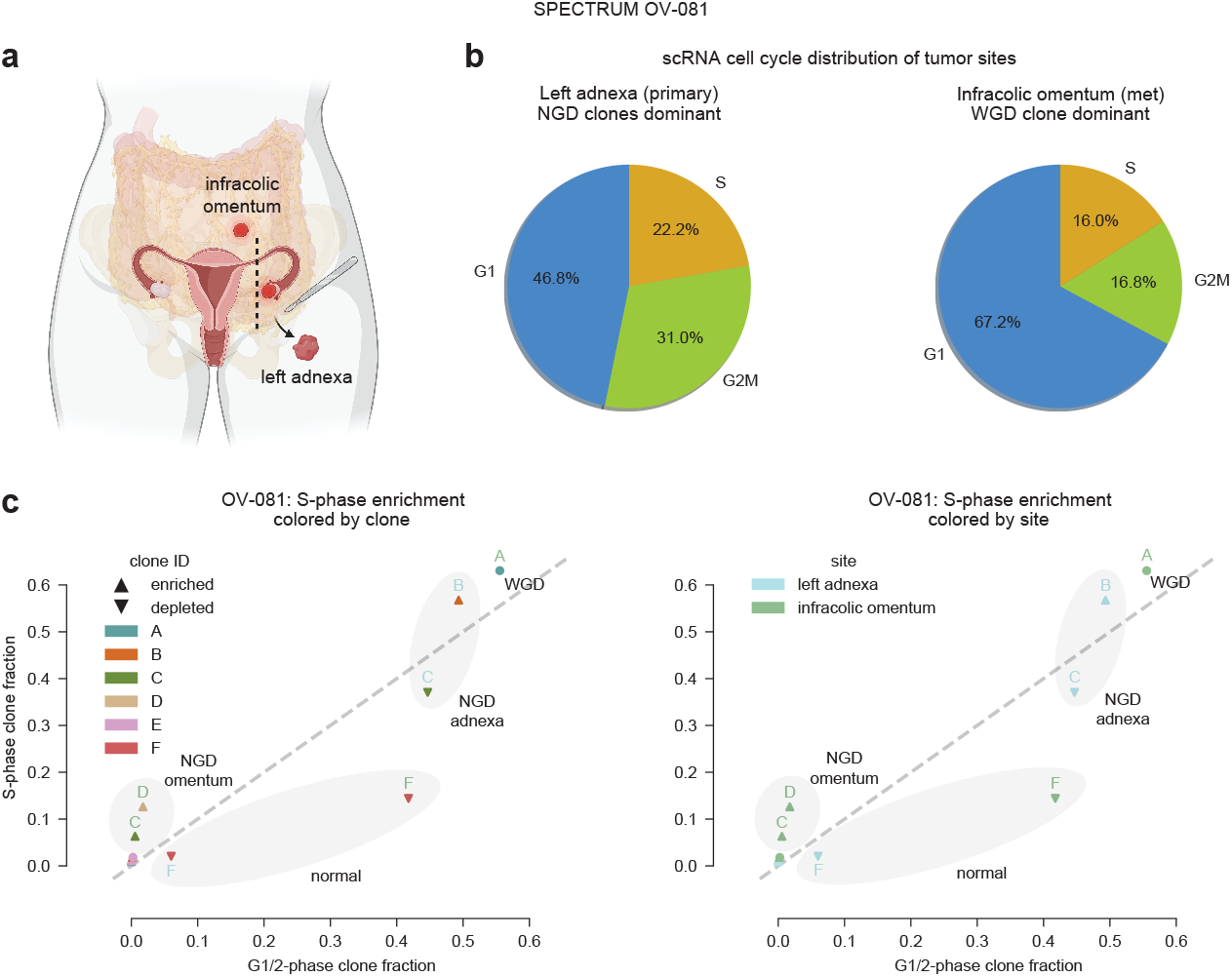
Whole-genome doubled clone in SPECTRUM patient OV-081 proliferates slower than the non-genome doubled clones and faster than normal cells. **a)** Anatomical sites of two samples collected from SPECTRUM patient OV-081 during primary debulking surgery prior to any treatment. **b)** Cell cycle phase distribution of scRNA tumor cells at each biopsy site. Cell cycle phases were determined by Seurat [62]. **c)** Clone fraction in S-vs G1/2-phase scWGS populations for each OV-081 clone within each site. Points are colored by clone on the left and site on the right. Points with upward pointing triangles are significantly (hypergeometric *p*_*adj*_ *<* 0.01) enriched for S-phase cells relative to other clones in the same site; points with downward pointing triangles are significantly depleted for S-phase cells. Points are annotated by their ploidy/tumor status and their site.

## Notes

### Summary of Updates

Additional analysis of human tumors has been added to this manuscript, resulting in new biological insights. All figures and supplemental files have been updated accordingly.

