## Supplemental notes and figures for "Single-cell DNA replication dynamics in genomically unstable cancers"

#### Supplementary Notes

##### Note 1: Evaluation of performance on simulated data

We benchmarked PERT's accuracy at inferring somatic CN, replication states, and cell cycle phase through quantitative simulation experiments. We simulated parameter sweeps over the number of clones, the cell-specific CNA rate, replication stochasticity, read depth overdispersion, GC bias coefficients, and RT profiles (Table S1). A representative example of true and PERT inferred CN and scRT profiles can be seen for dataset P5.8 which has 3 clones and a cell-specific CNA rate of 0.02, resulting in a replication state accuracy (bins correctly assigned as replicated or unreplicated) of 96.5% (Fig. 1c). We benchmarked PERT against the DLP+ cell cycle classifier presented in Laks *et al* [1] for per-cell phase accuracy and Kronos [2] for per-bin scRT accuracy as they are the only available softwares for predicting cell cycle and scRT states, respectively. The Laks classifier achieved similar performance to the initial version (AUC=0.92) on a train-test split after removing low importance sequence-level features not available in our simulated data, with its most important features remaining high correlations and slopes between per-bin GC content and read depth (Fig. S1).

PERT outperformed the Laks classifier and Kronos in all simulated datasets. The performance gap between PERT and Kronos was significant ($p_{adj} < 10^4$) for all parameter combinations and increased as a function of cell CNA rate, number of clones, and noise $\lambda$ (Fig. S2). PERT's replication and CN accuracies decayed at high cell CNA rate and $\lambda$ values but remained robust to extreme replication stochasticity ($\alpha$), GC bias ($\beta_\mu$), and clone number (Figs. S2a-r). PERT accurately estimated the fraction of replicated loci in all cells and thus achieved 93% phase accuracy across all simulated datasets with $\lambda = 0.75$, performing significantly better ($p_{adj} < 10^4$) than the Laks classifier in all parameter sweeps (Fig. S2s-ab). Importantly, PERT's cell cycle phase accuracy was robust to fluctuations in GC bias slope ($\beta_{\mu,0}$), indicating that PERT's performance will not fluctuate between DLP+, 10x, and other scWGS modalities with unique GC bias relationships (Fig. S2ab). Together this analysis demonstrates that PERT provides significant improvement upon existing methods to infer scRT states, and thus cell cycle phase, from scWGS data – particularly in cases where CNAs arise with subclonal structure.

#### Note 2: Examining scRT errors

We investigated scRT errors in representative dataset P5.8 to ensure that the bias of errors made sense in the context of the methods of PERT and Kronos. Kronos errors came at bins with subclonal or cell-specific CNAs whereas PERT errors arose from cell-specific CNAs not shared between S- and G1/2-phase cells (Fig. S3a-c). Such modes of failure make sense as the Kronos method assumes that all S-phase possess the same somatic copy number profile as the G1/2-phase pseudobulk whereas PERT leverages prior information from all G1/2-phase cells when predicting the somatic copy number profiles of S-phase cells. Another source of error was confusion between very early and very late S-phase cells with PERT making fewer said mistakes than Kronos (Fig. S3d,e). Mixing up very early and very late S-phase cells is an expected failure case since cells at both extremes begin to resemble G1/2-phase cells with flat profiles, making it harder to distinguish whether the cell should be mostly replicated or mostly unreplicated. This analysis demonstrates that PERT is working as intended and it is most sensitive to 1) very rare CNAs and 2) very early/late S-phase cells.

#### Note 3: PERT as a data-driven S-phase classifier in unsorted libraries

We compared PERT's phase predictions to the Laks *et al* classifier for unsorted DLP+ data of genetically engineered mammary epithelial 184-hTERT cell lines which contain many clonal, subclonal, and cell-specific CNAs [3] (Fig. 3a, Additional File 1). Sample SA1292 (TP53-/-, BRCA1+/- hTERT) is shown as a representative example (Fig. S4a-h). While the S-phase predictions were largely concordant between PERT and the Laks classifier, PERT recovered an additional 417 G1/2-phase cells which were filtered out as low-quality by the Laks classifier (Fig. S4i). The recovered Laks=LQ, PERT=G1/2 cells clustered with other G1/2-phase cells of the same clone in PCA embedding space and had orthogonal cell cycle features consistent with G1/2-phase cells, suggesting that these cells are indeed G1/2-phase and not low quality (Fig. S4j-l). When comparing PERT vs Laks predicted phases across all hTERT samples, the majority of Laks=G1/2 (13941/15042, 92.7%) and Laks=S (3264/5102, 64.0%) cells retained the same PERT phase but 34.6% (1766/5102) of Laks=S and 52.3% (1810/3464) of Laks=LQ cells were called as PERT=G1/2 (Fig. S4m). PERT detected >5x fewer LQ cells than Laks (3464 to 591) with many of the recovered cells having erroneously high ploidy (true ploidy is N=2) and thus higher breakpoint counts according to the input CN states (Fig. S4n). Reducing the number of LQ cells enables PERT to increase the yield of scWGS libraries, such as SA1292, which had many G1/2-phase cells excluded from previous analysis which increases the power of downstream sequence-level analyses such as SNV and complex SV calling which must be done at the pseudobulk level. This evidence suggests that PERT's direct modeling of how DNA replication, somatic CN, and library-level biases combine to produce observed read count provides high-fidelity cell cycle prediction across diverse data.

#### Note 4: Relationship between S-phase enrichment and clonal expansion in untreated time-series cell lines

To better understand the context in which we only saw a modest positive correlation between S-phase enrichment (SPE) and observed clonal expansion between subsequent timepoints from the untreated TNBC PDX models (Fig. 6a-f), we similarly analyzed untreated time-series hTERT samples with and without TP53 ablation [4] (Fig. S5a-d). Much like the untreated PDX samples, there was only a slight positive correlation between clone SPE and expansion for this data (Fig. S5e-h). We hypothesize that this correlation is noisy due to confounders such as serial passaging between scWGS time-points, destructive sequencing, and parent-child relationships between clones which introduce artificial evolutionary bottlenecks that distort the relationship between true proliferation rate and observed clone expansion rate.

#### Note 5: Additional effects of cisplatin on replication in time-series TNBC PDX data

We investigated additional replication properties of the TNBC PDXs outside of their relationship between S-phase enrichment, clonal expansion, and treatment status (Fig

6a-f). We found that RT profiles were unique between PDXs (Pearson r 0.82-0.87) and highly conserved between on- and off-treatment groups of the same PDX (Pearson r 0.94-0.99) (Fig. S6a), suggesting that there are patient-specific RT programs and that cisplatin does not disrupt genome-wide RT coordination. Additionally, we found on-treatment SA1035 S-phase cells to be earlier in S-phase than off-treatment SA1035 S-phase cells ($p_{adj} < 10^4$); however the on- vs off-treatment distributions had no significant difference for the other two PDXs ($p_{adj} < 10^2$) (Fig. S6b). Finally, on-treatment samples had >2x higher S-phase fraction (SPF) than off-treatment samples in SA1035 and SA535 (Fig. S6c). The synthesis of all this supporting data leads us to believe that cisplatin might be generating more fork stalling in SA1035 and SA535 than in SA609.

### Tables

**Table 1** Parameters values for simulated datasets

| Datatag | # clones | CNA rate | $\lambda$ | $\alpha$ | ENCODE RT | $\beta_\mu$ |
|---|---|---|---|---|---|---|
| D1 | 1 | 0 | 0.75 | 10 | MCF7 | [1.2, 0] |
| D2 | 1 | 0 | 0.75 | 5 | MCF7 | [1.2, 0] |
| D3 | 1 | 0 | 0.75 | 15 | MCF7 | [1.2, 0] |
| D4 | 1 | 0 | 0.75 | 10 | MCF7 | [-1.2, 0] |
| D5 | 1 | 0 | 0.5 | 10 | MCF7 | [1.2, 0] |
| D6 | 1 | 0 | 0.9 | 10 | MCF7 | [1.2, 0] |
| D7 | 1 | 0.02 | 0.75 | 10 | MCF7 | [1.2, 0] |
| D8 | 1 | 0.05 | 0.75 | 10 | MCF7 | [1.2, 0] |
| D9 | 1 | 0 | 0.99 | 10 | MCF7 | [1.2, 0] |
| D10 | 1 | 0 | 0.6 | 10 | MCF7 | [1.2, 0] |
| P1 | 3 | 0 | 0.75 | 10 | MCF7 | [1.2, 0] |
| P1 | 3 | 0 | 0.75 | 10 | MCF7 | [1.2, 0] |
| P2 | 3 | 0.02 | 0.75 | 10 | MCF7 | [1.2, 0] |
| P3 | 3 | 0.05 | 0.75 | 10 | MCF7 | [1.2, 0] |
| P4 | 3 | 0 | 0.75 | 5 | MCF7 | [1.2, 0] |
| P5 | 3 | 0.02 | 0.75 | 5 | MCF7 | [1.2, 0] |
| P6 | 3 | 0.05 | 0.75 | 5 | MCF7 | [1.2, 0] |
| P7 | 3 | 0 | 0.75 | 15 | MCF7 | [1.2, 0] |
| P8 | 3 | 0.02 | 0.75 | 15 | MCF7 | [1.2, 0] |
| P9 | 3 | 0.05 | 0.75 | 15 | MCF7 | [1.2, 0] |
| P10 | 4 | 0.02 | 0.75 | 10 | GM-06990 -12801 -12812 -12813 | [1.2, 0] |
| P11 | 4 | 0.02 | 0.75 | 10 | BJ MCF7 HEPG2 GM12813 | [1.2, 0] |

### Figures

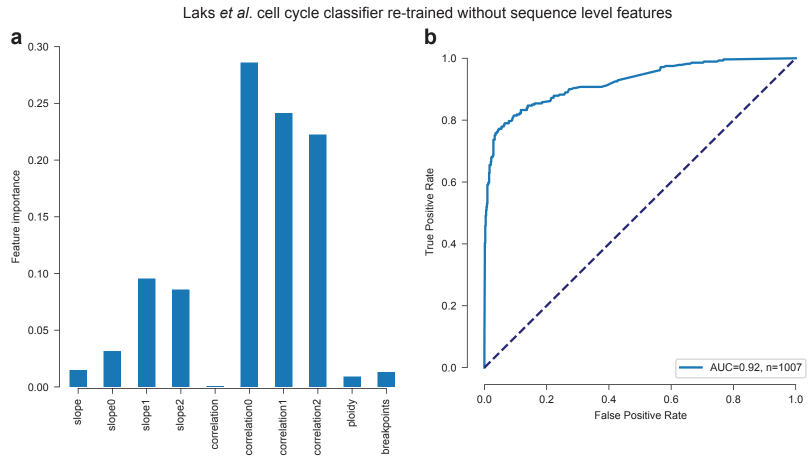


**Fig. S1 Retraining Laks *et al* cell cycle classifier without sequence-level features a)** Feature importance scores and **b)** train-test ROC curve for the DLP+ cell cycle classifier described in Laks *et al* [1]. Features beginning with 'slope' or 'correlation' measure the relationship between GC and various forms of normalized read depth. Features with no suffix represent slopes and correlations between GC and reads per million normalized by integer copy number state. Features with suffix '0' are slopes and correlations between GC and reads per million normalized by integer copy number state and the per-library GC bias curve. Features with the suffix '1' are the same as those with suffix '0' except they use raw read count instead of reads per million prior to normalization. Features with the suffix '2' are the same as those with suffix '0' except they normalize by the GC bias curve of each cell instead of each library.

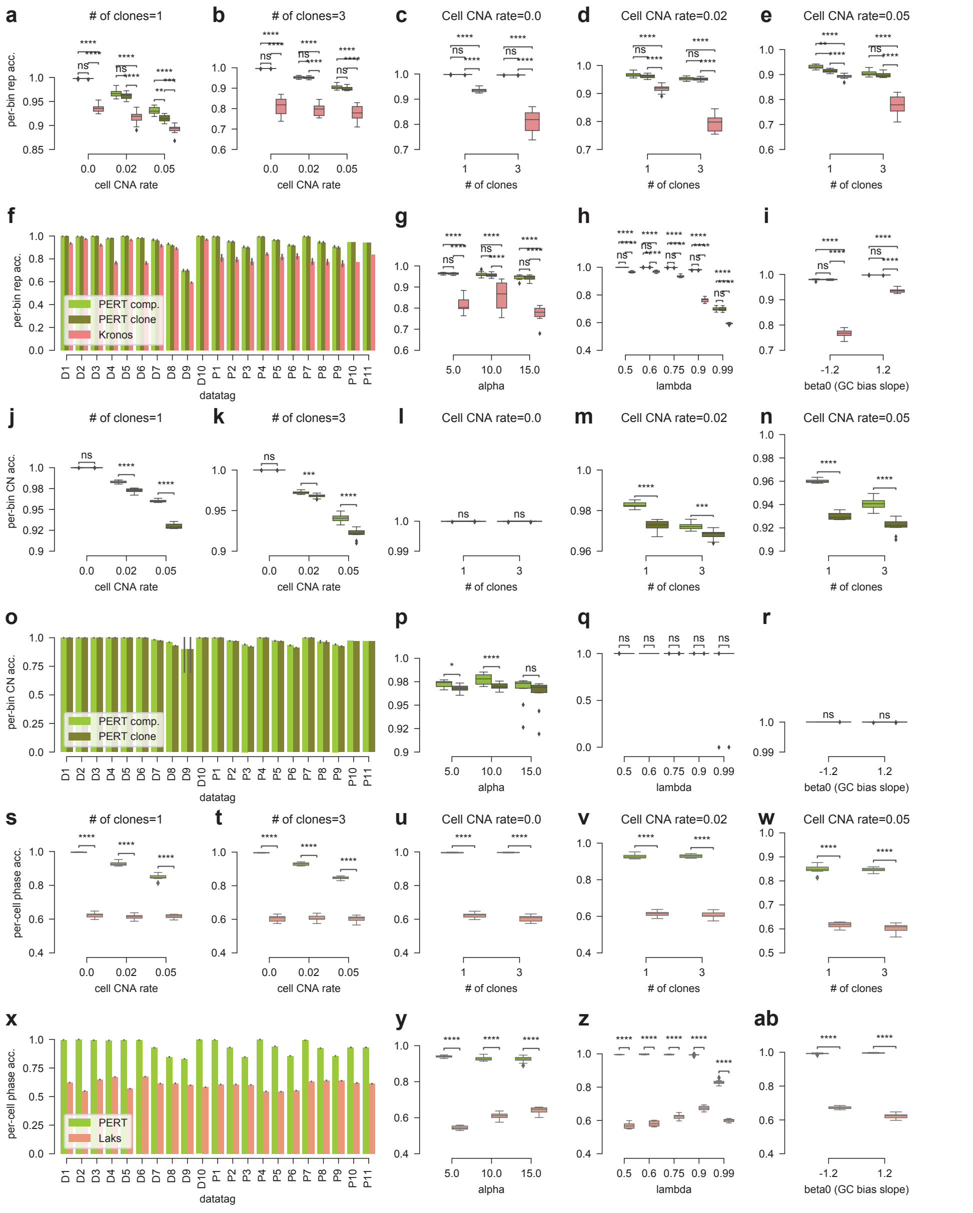


**Fig. S2 Comparison of PERT to Kronos scRT and Laks cell cycle classifier across all simulated datasets. a-i)** Per-bin replication state accuracy for Kronos and PERT with clone and composite CN prior methods in simulated S-phase cells. **a-b)** Parameter sweep across cell-specific CNA rate. **c-e)** Parameter sweep across number of clones. **f)** Accuracy for all unique combinations of simulation parameters (datatag). There are 10 unique datasets for each datatag. **g)** Parameter sweep across the replication stochasticity term $\alpha$. **h)** Parameter sweep across the read depth overdispersion term $\lambda$. **i)** Parameter sweep across the GC bias slope term $\beta_{\mu,1}$. **j-r)** Per-bin somatic copy number state accuracy for PERT with clone and composite CN prior methods in simulated S-phase cells. Ordering of subplots is the same as **a-i**. **s-ab)** Per-cell phase accuracy for the Laks cell cycle classifier and PERT with the composite CN prior in all simulated cells. Ordering of subplots is the same as **a-i**.

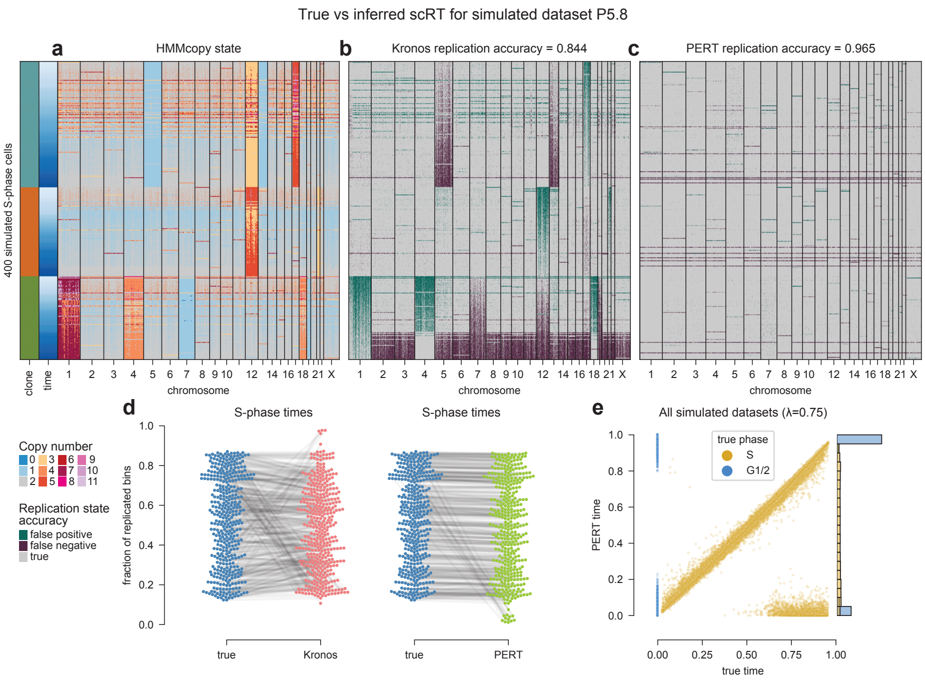


**Fig. S3 Examining scRT errors.** Kronos and PERT scRT errors for true S-phase cells in dataset P5.8. **a)** HMMcopy states for all S-phase cells. **b-c)** Kronos and PERT per-bin replication accuracies. False positives are true unreplicated bins inferred as replicated; false negatives are true replicated bins inferred as unreplicated. Rows in **a-c** are sorted by clone and true S-phase time. **d)** True vs inferred S-phase times for each method. Lines represent the mapping of true to inferred values for the same simulated S-phase cell. **e)** Comparison of true vs PERT inferred S-phase times across all simulated datasets with $\lambda = 0.75$.

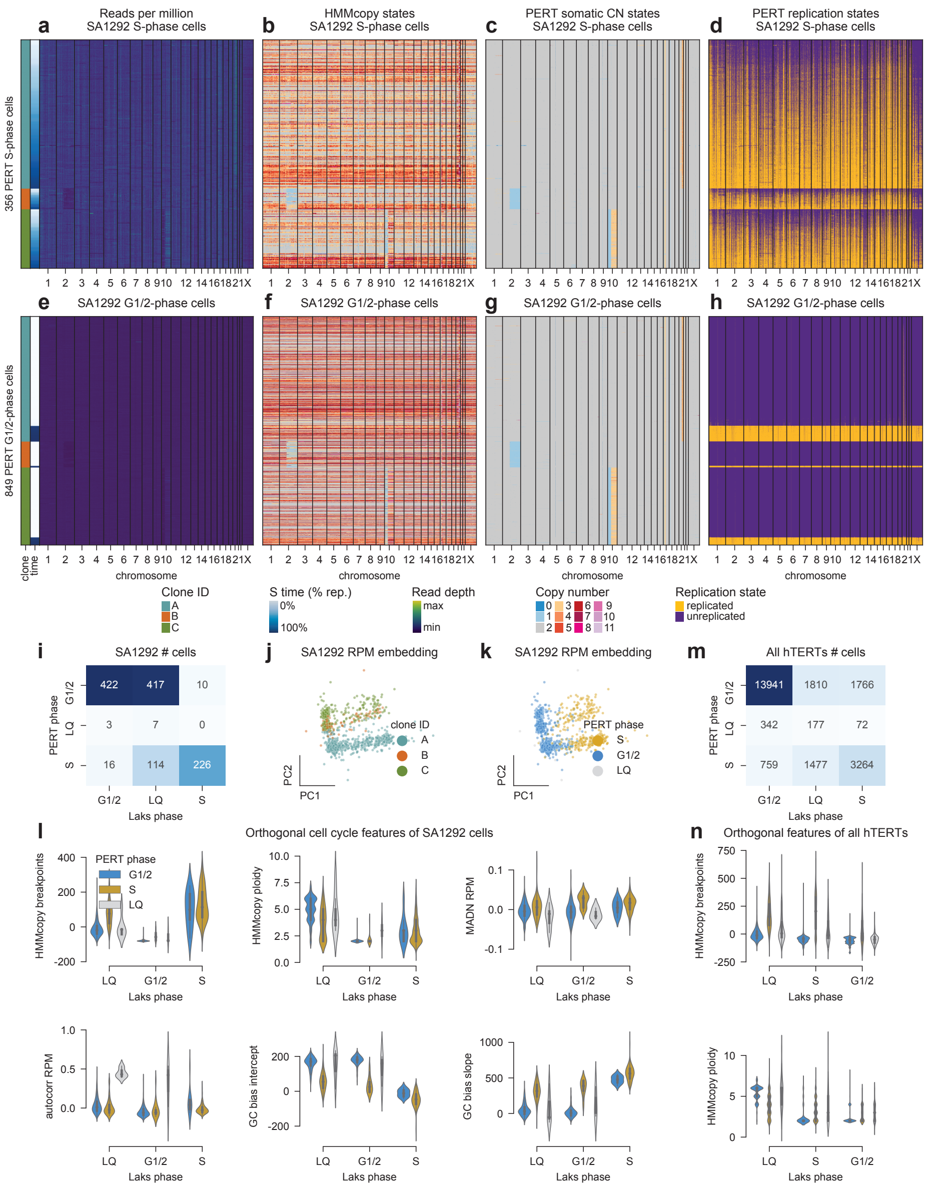


**Fig. S4 PERT phase prediction recovers many low-quality cells. a-l)** PERT output for hTERT TP53-/- BRCA1+/- sample SA1292. **a)** Reads per million, **b)** HMMcopy states [5], **c)** PERT somatic copy number states, and **d)** PERT replication states for cells predicted as S-phase and **e-h)** G1/2-phase by PERT. Rows (cells) are sorted first by clone ID and then fraction of replicated bins. **i)** Confusion matrix of Laks *et al* vs PERT cell cycle phases for sample SA1292. **j-k)** PCA embedding of reads per million where cells are colored by clone ID and PERT phase. **l)** per-cell features correlated with quality and cell cycle phase where each violin corresponds to a unique position of the Laks vs PERT phase confusion matrix. Features include corrected number of HMMcopy copy number breakpoints (subtracted by the mean), HMMcopy ploidies, corrected median absolute deviation of reads per million between neighboring bins (MADN RPM), normalized autocorrelation of reads per million profiles, and slope and intercept coefficients from a linear regression fit between GC and read count. **m-n)** PERT output for all hTERT cell line samples. **m)** Confusion matrix of Laks *et al* vs PERT cell cycle phases. **n)** per-cell features correlated with quality and cell cycle phase where each violin corresponds to a unique position of the Laks vs PERT phase confusion matrix. Features shown are mean-corrected HMMcopy copy number breakpoints and HMMcopy ploidies.

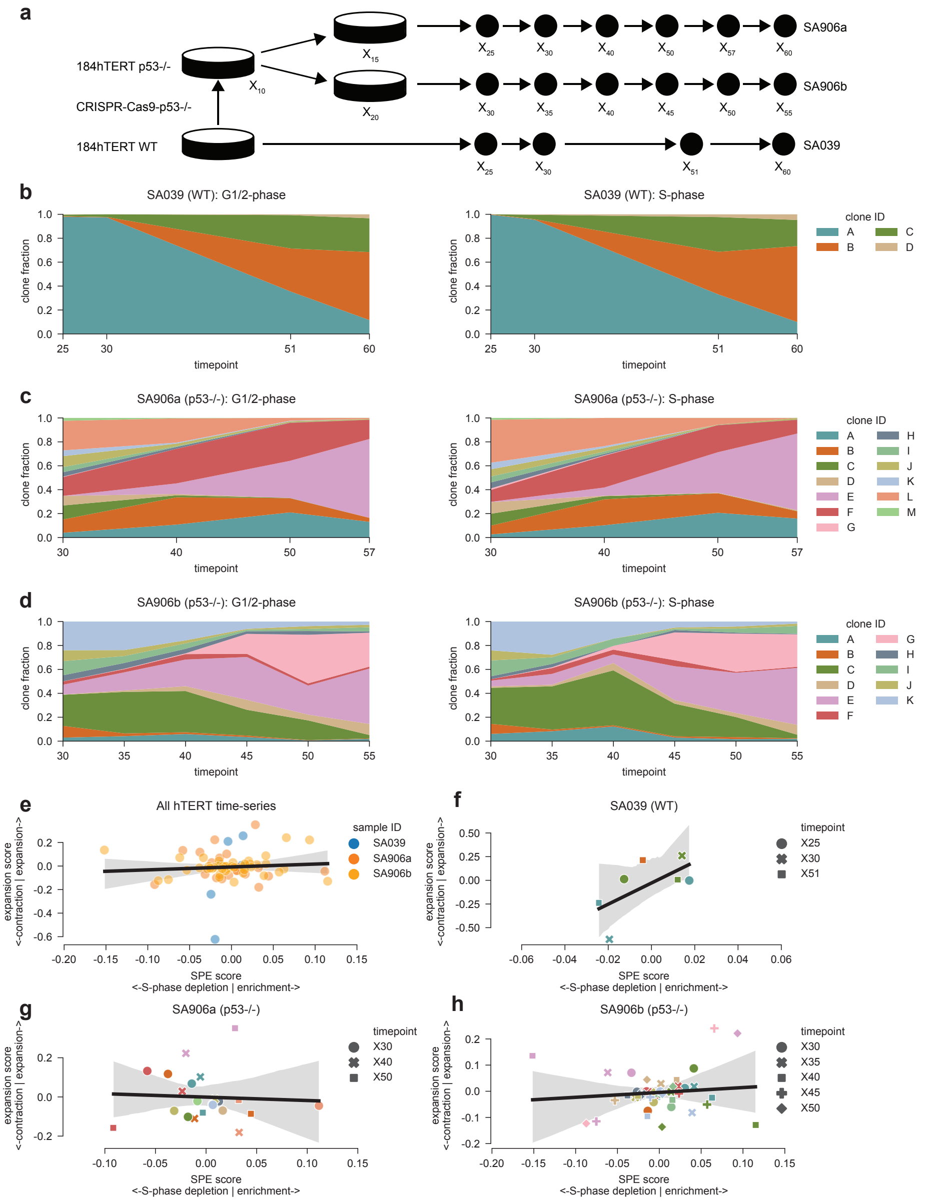


**Fig. S5 Time-series scWGS sampling of untreated hTERT cell lines. a)** Schematic of time-series scWGS sampling for hTERT WT and TP53-/- cell lines. **b)** Relationship between SPE and clone expansion for each clone and timepoint combination with > 10 G1/2-phase cells across all three samples and **c-e)** split by sample. Lines represent linear regression fits with shaded areas representing 95% confidence intervals.

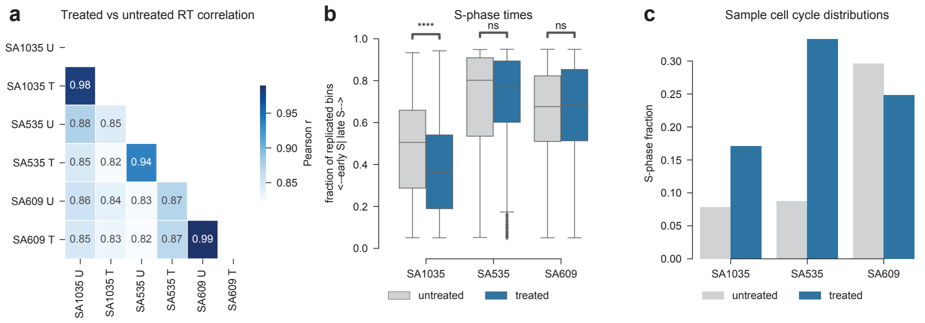


**Fig. S6 Impact of cisplatin on DNA replication dynamics on each TNBC PDX. a)** Pairwise Pearson correlation between treated and untreated sample RT profiles. **b)** Distribution of cell S-phase times between treated and untreated samples. **c)** Total fraction of S-phase cells between treated and untreated samples.
