## Additional File 1 for "Single-cell DNA replication dynamics in genomically unstable cancers"

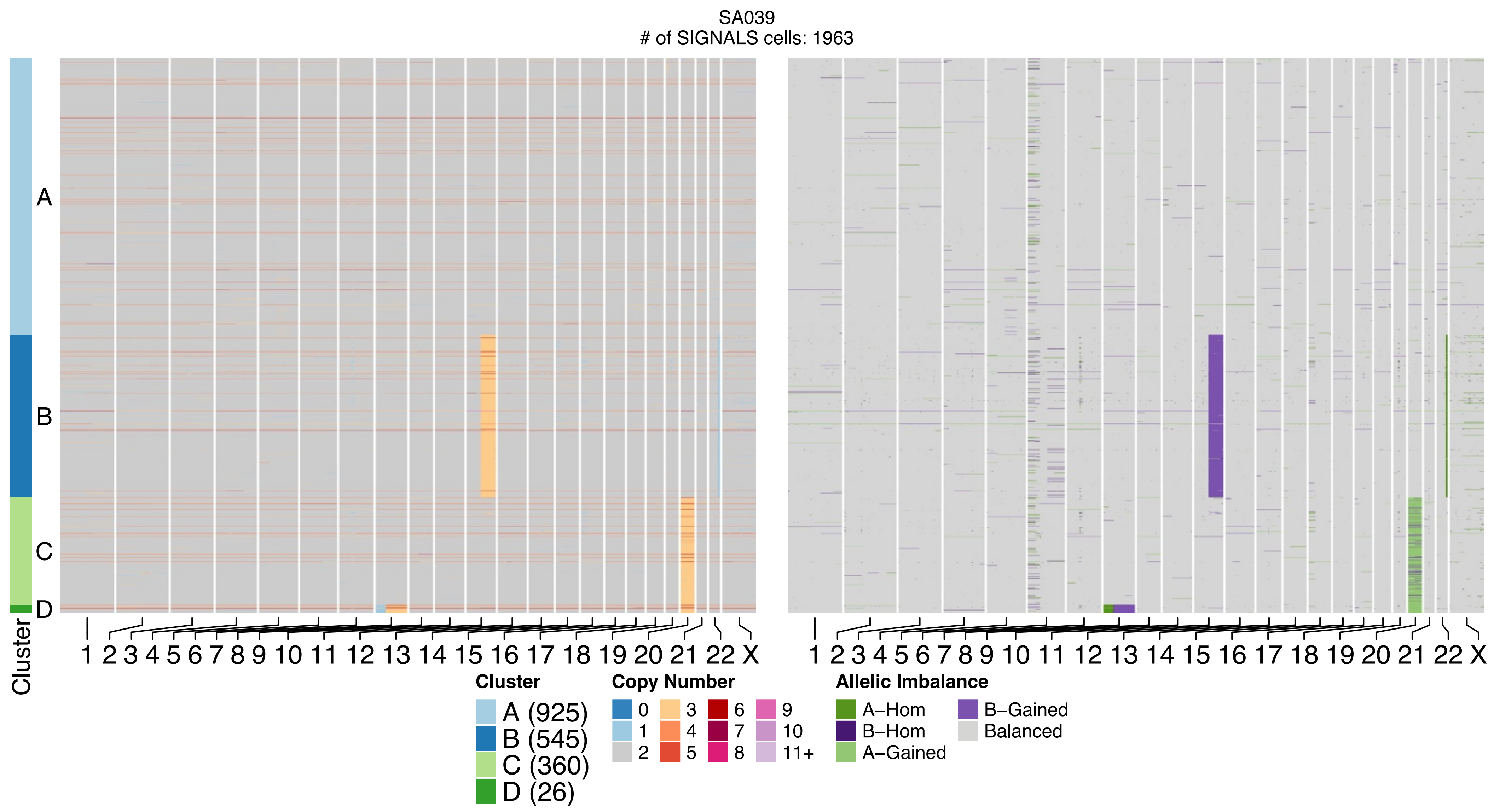
SA039
### of SIGNALS cells: 1963
A
B
C
D
Cluster
1 2 3 4 5 6 7 8 9 10 11 12 13 14 15 16 17 18 19 20 21 22 X
Cluster
A (925)
B (545)
C (360)
D (26)
Copy Number
0 1 2 3 4 5 6 7 8 9 10 11+
Allelic Imbalance
A–Hom
B–Hom
A–Gained
B–Gained
Balanced

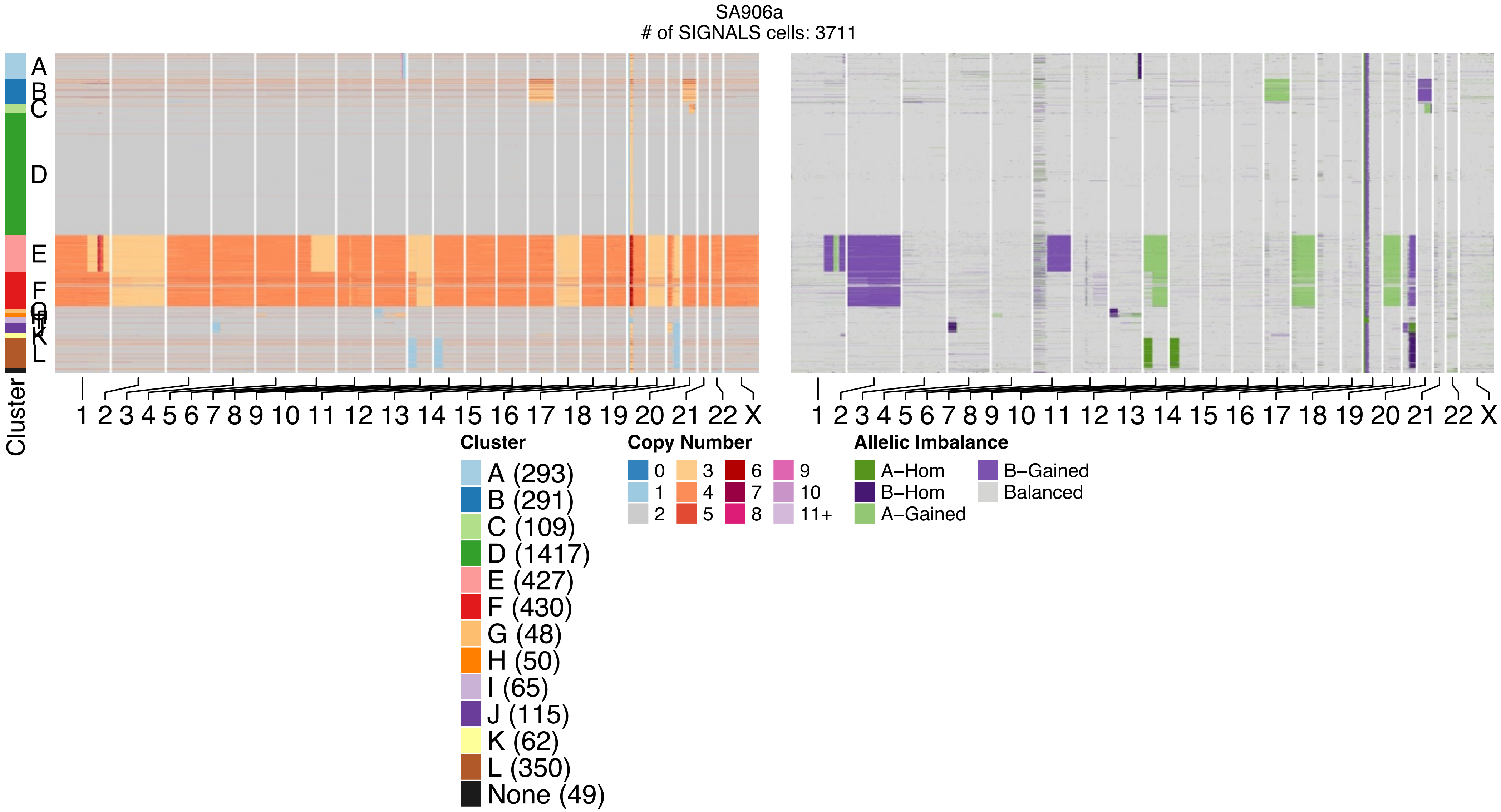

SA906a
### of SIGNALS cells: 3711
A
B
C
D
E
F
G
H
I
J
K
L
Cluster
1 2 3 4 5 6 7 8 9 10 11 12 13 14 15 16 17 18 19 20 21 22 X
1 2 3 4 5 6 7 8 9 10 11 12 13 14 15 16 17 18 19 20 21 22 X
Cluster
A (293)
B (291)
C (109)
D (1417)
E (427)
F (430)
G (48)
H (50)
I (65)
J (115)
K (62)
L (350)
None (49)
Copy Number
0
1
2
3
4
5
6
7
8
9
10
11+
Allelic Imbalance
A–Hom
B–Hom
A–Gained
B–Gained
Balanced

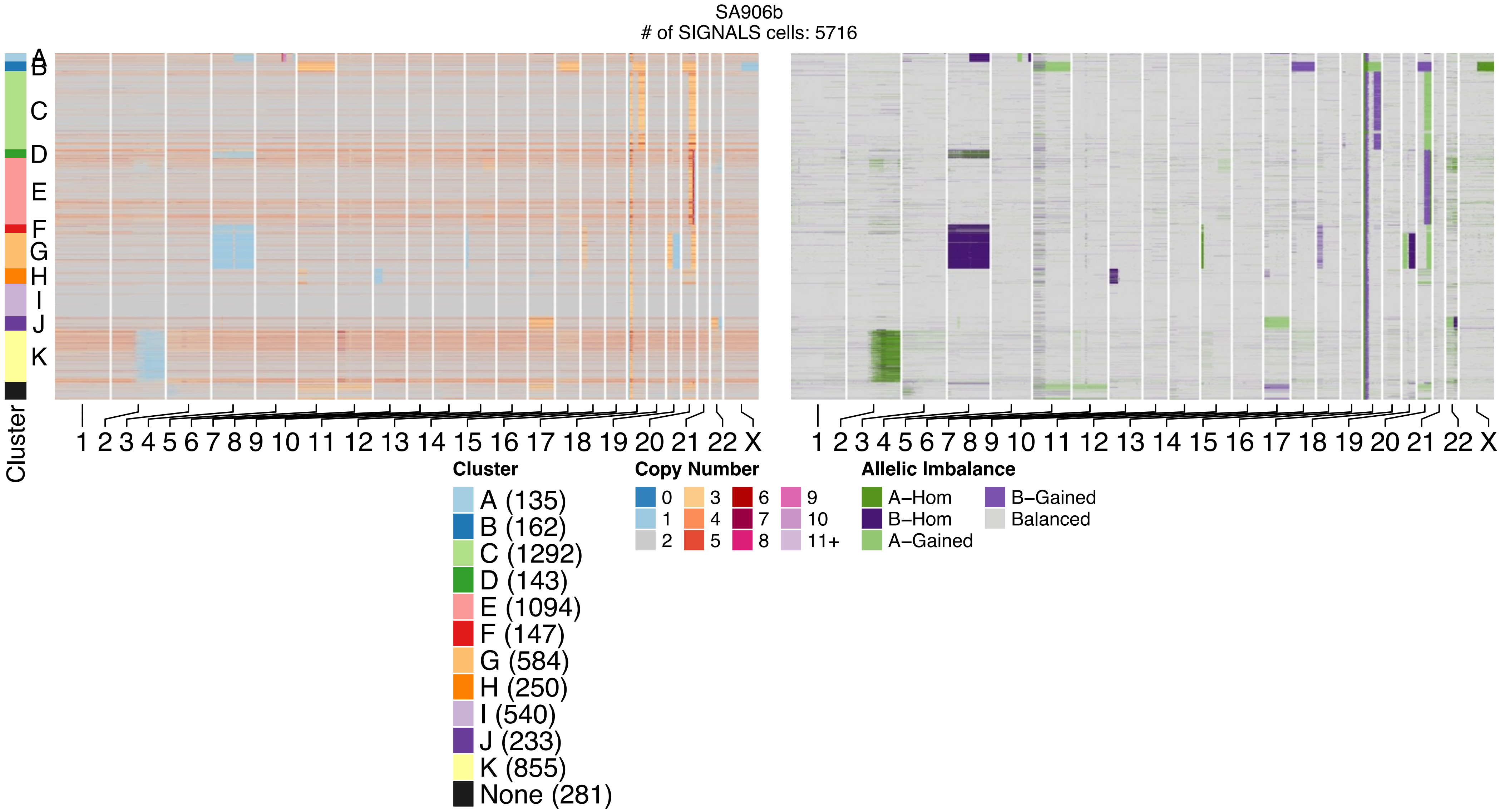
SA906b
### of SIGNALS cells: 5716
A
B
C
D
E
F
G
H
I
J
K
Cluster
1 2 3 4 5 6 7 8 9 10 11 12 13 14 15 16 17 18 19 20 21 22 X
1 2 3 4 5 6 7 8 9 10 11 12 13 14 15 16 17 18 19 20 21 22 X
Cluster
A (135)
B (162)
C (1292)
D (143)
E (1094)
F (147)
G (584)
H (250)
I (540)
J (233)
K (855)
None (281)
Copy Number
0
1
2
3
4
5
6
7
8
9
10
11+
Allelic Imbalance
A−Hom
B−Hom
A−Gained
B−Gained
Balanced

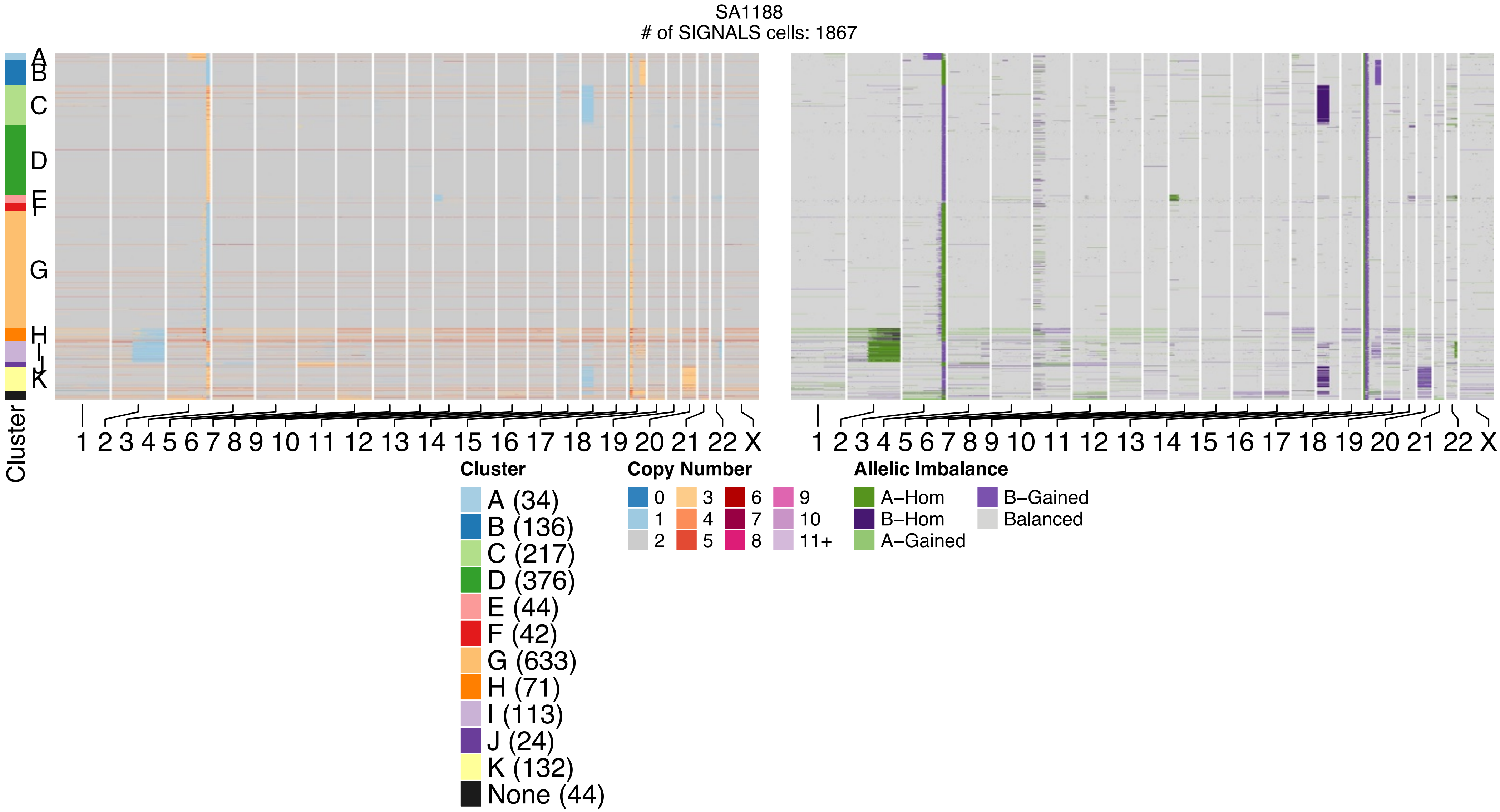
SA1188
### of SIGNALS cells: 1867
Cluster
A
B
C
D
E
F
G
H
I
J
K
1 2 3 4 5 6 7 8 9 10 11 12 13 14 15 16 17 18 19 20 21 22 X
1 2 3 4 5 6 7 8 9 10 11 12 13 14 15 16 17 18 19 20 21 22 X
Cluster
A (34)
B (136)
C (217)
D (376)
E (44)
F (42)
G (633)
H (71)
I (113)
J (24)
K (132)
None (44)
Copy Number
0
1
2
3
4
5
6
7
8
9
10
11+
Allelic Imbalance
A−Hom
B−Hom
A−Gained
B−Gained
Balanced

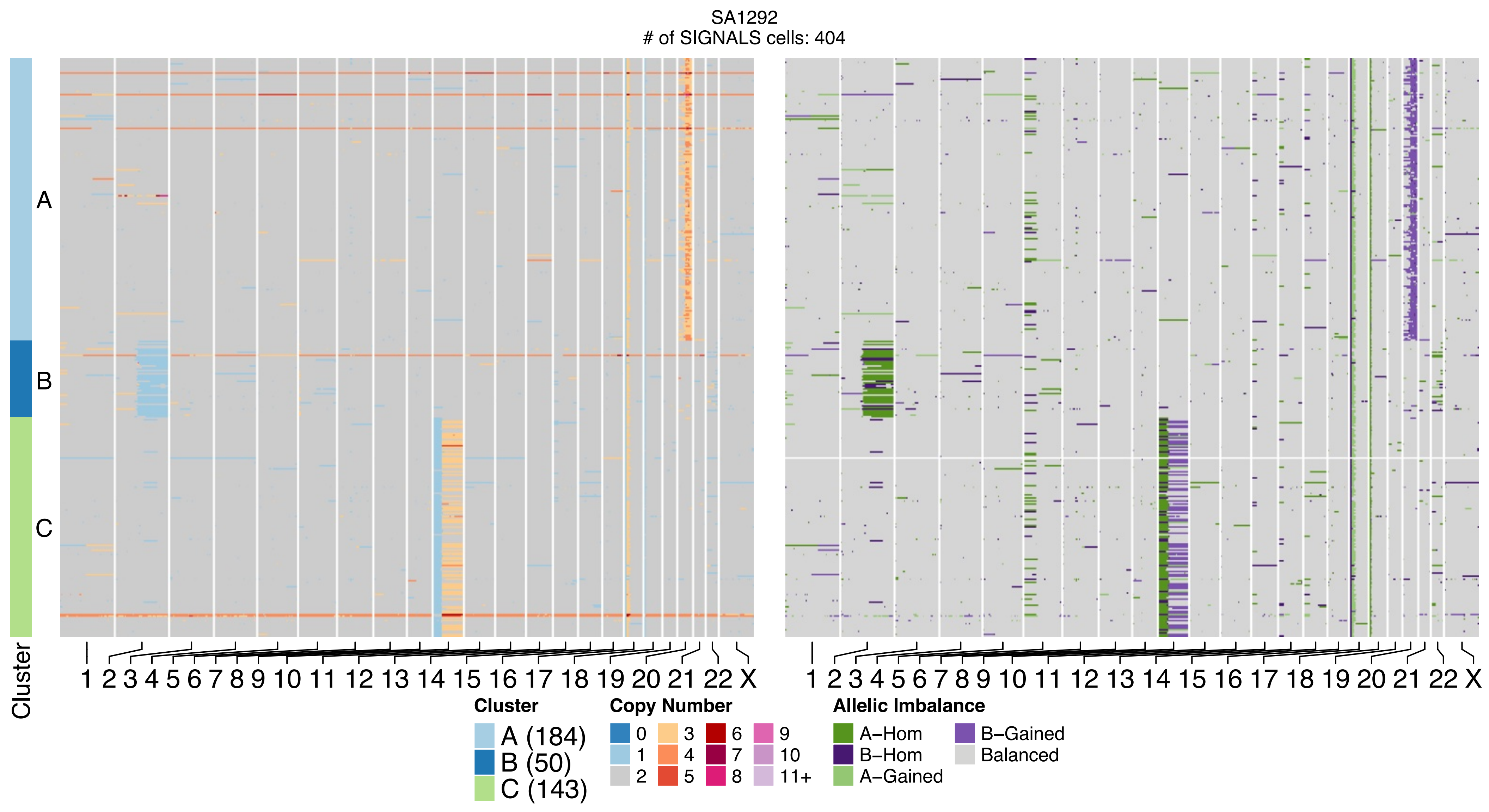
SA1292
### of SIGNALS cells: 404
A
B
C
Cluster
1 2 3 4 5 6 7 8 9 10 11 12 13 14 15 16 17 18 19 20 21 22 X
1 2 3 4 5 6 7 8 9 10 11 12 13 14 15 16 17 18 19 20 21 22 X
Cluster
A (184)
B (50)
C (143)
Copy Number
0 1 2 3 4 5 6 7 8 9 10 11+
Allelic Imbalance
A−Hom
B−Hom
A−Gained
B−Gained
Balanced

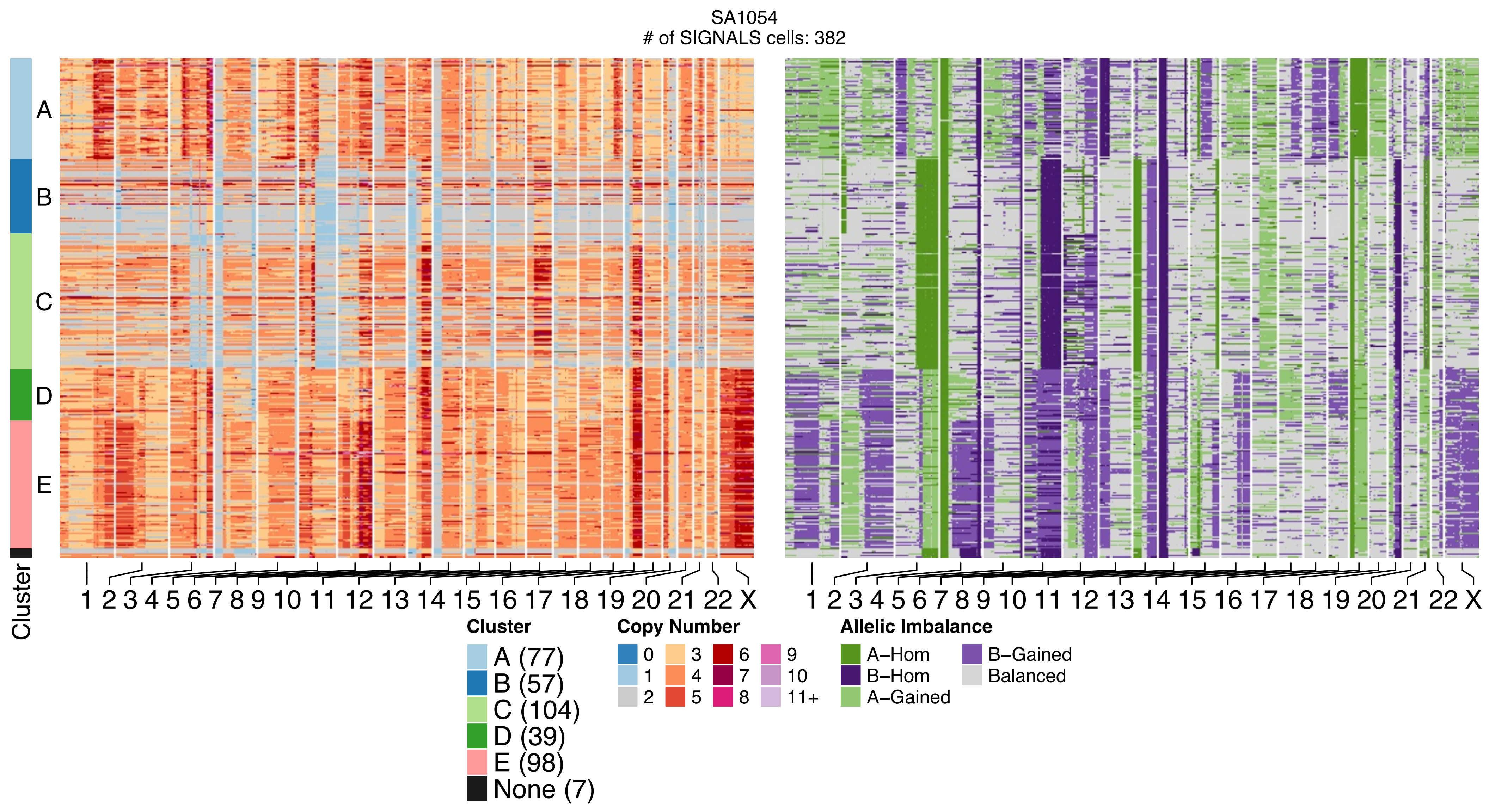
SA1054
### of SIGNALS cells: 382
A
B
C
D
E
Cluster
1 2 3 4 5 6 7 8 9 10 11 12 13 14 15 16 17 18 19 20 21 22 X
1 2 3 4 5 6 7 8 9 10 11 12 13 14 15 16 17 18 19 20 21 22 X
Cluster
A (77)
B (57)
C (104)
D (39)
E (98)
None (7)
Copy Number
0
1
2
3
4
5
6
7
8
9
10
11+
Allelic Imbalance
A–Hom
B–Hom
A–Gained
B–Gained
Balanced

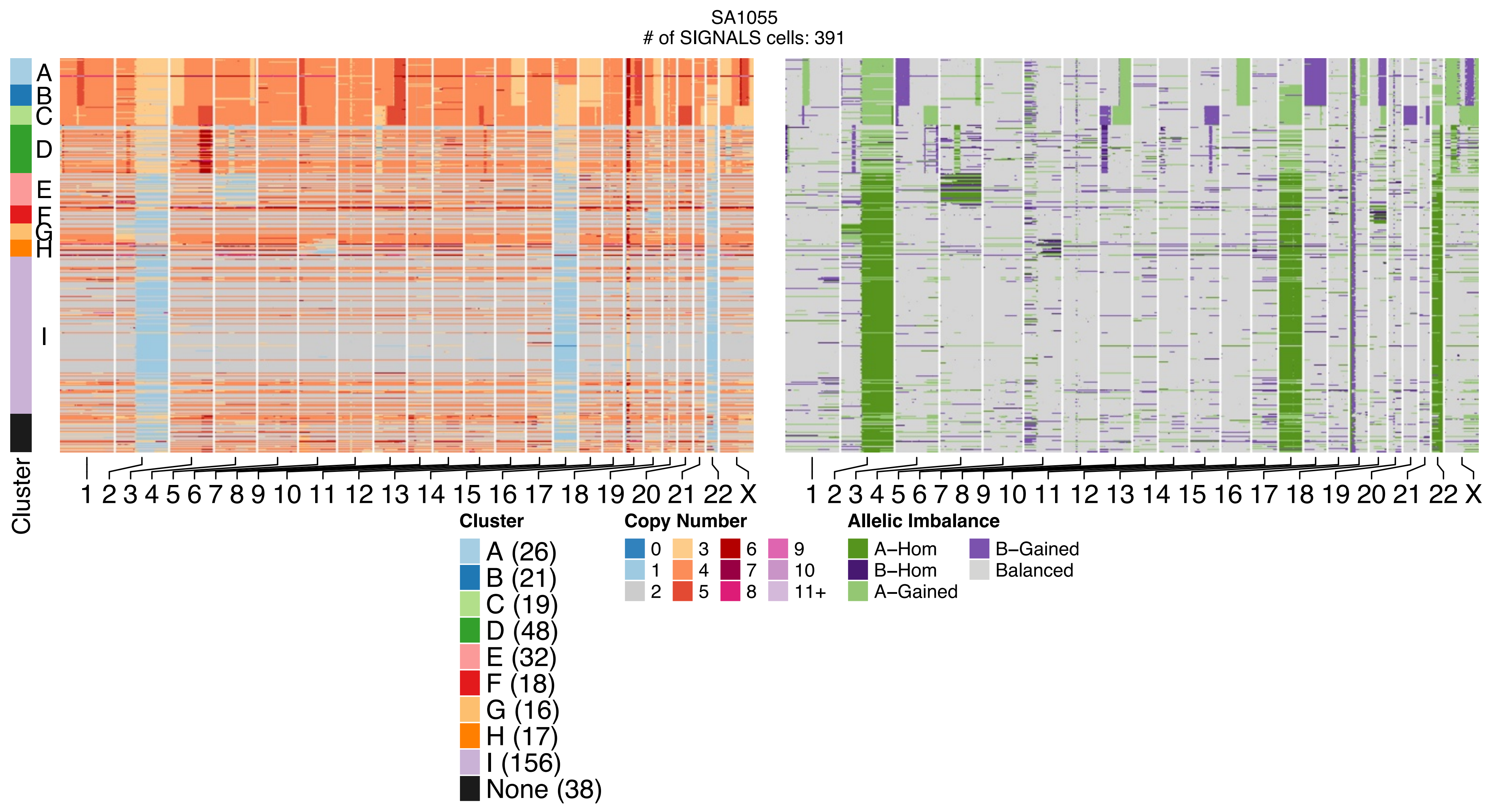
SA1055
### of SIGNALS cells: 391
Cluster
A
B
C
D
E
F
G
H
I
1 2 3 4 5 6 7 8 9 10 11 12 13 14 15 16 17 18 19 20 21 22 X
1 2 3 4 5 6 7 8 9 10 11 12 13 14 15 16 17 18 19 20 21 22 X
Cluster
A (26)
B (21)
C (19)
D (48)
E (32)
F (18)
G (16)
H (17)
I (156)
None (38)
Copy Number
0
1
2
3
4
5
6
7
8
9
10
11+
Allelic Imbalance
A–Hom
B–Hom
A–Gained
B–Gained
Balanced

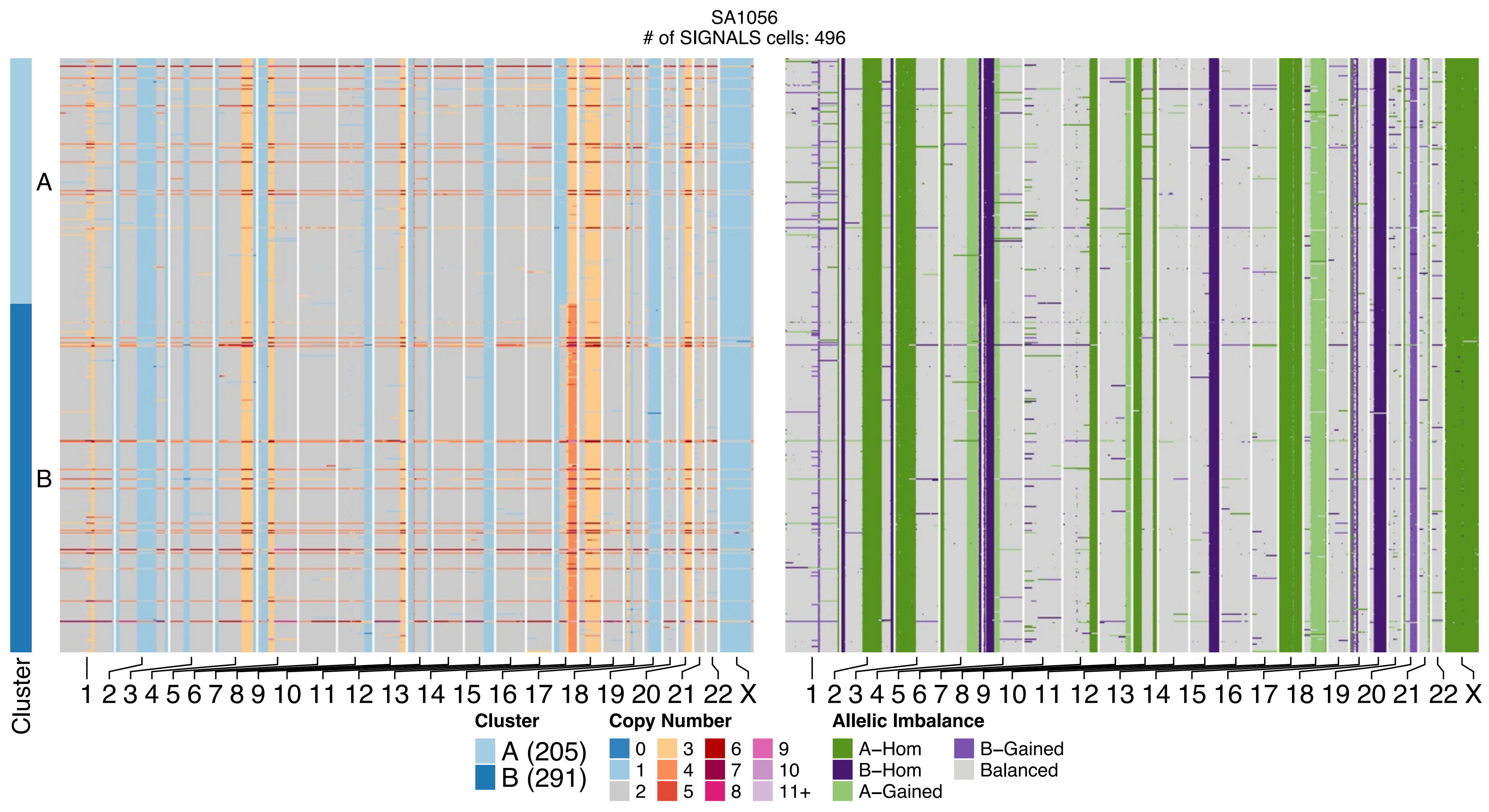
SA1056
### of SIGNALS cells: 496
A
B
Cluster
1 2 3 4 5 6 7 8 9 10 11 12 13 14 15 16 17 18 19 20 21 22 X
1 2 3 4 5 6 7 8 9 10 11 12 13 14 15 16 17 18 19 20 21 22 X
Cluster
A (205)
B (291)
Copy Number
0
1
2
3
4
5
6
7
8
9
10
11+
Allelic Imbalance
A–Hom
B–Hom
A–Gained
B–Gained
Balanced

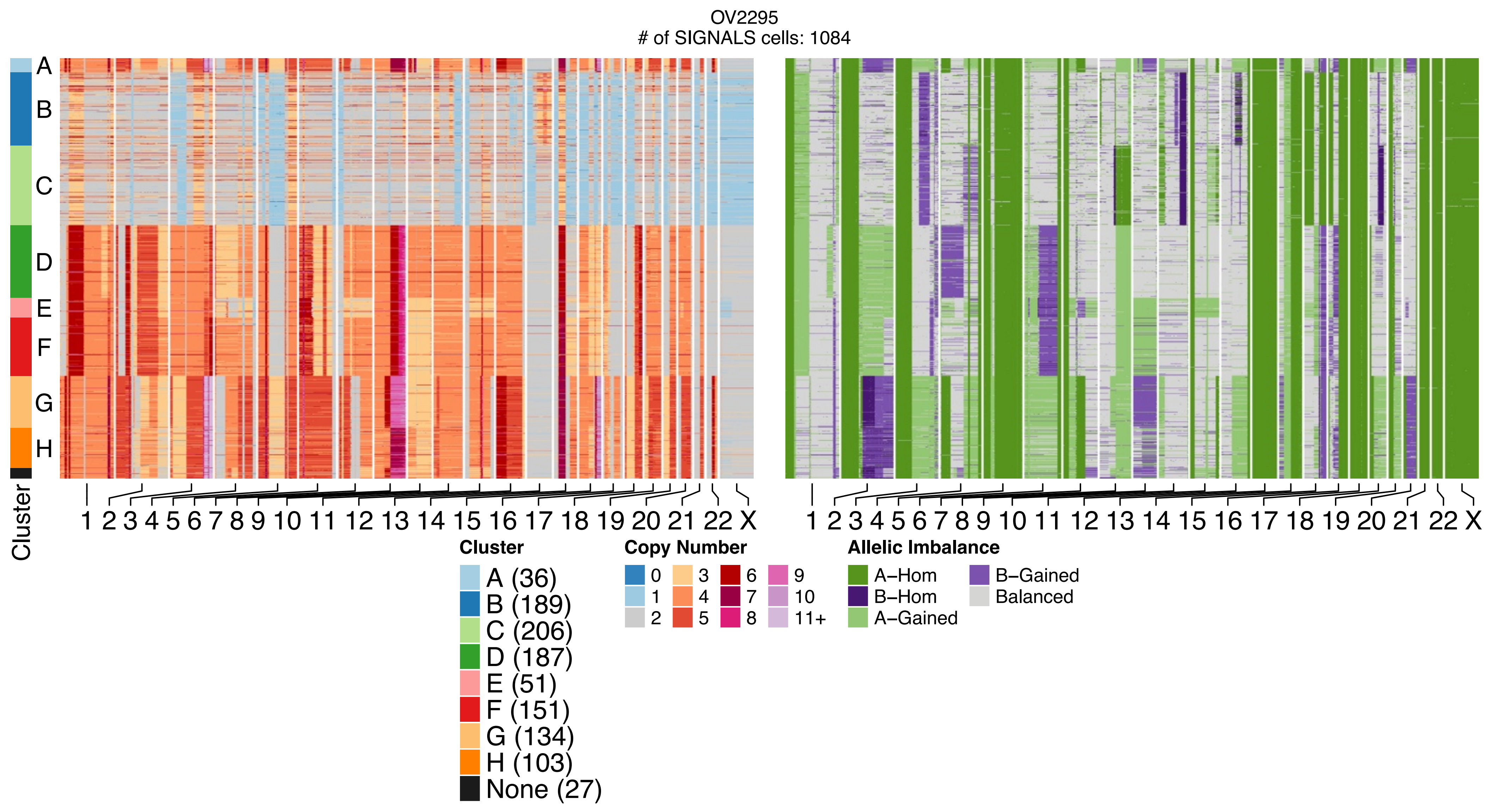
OV2295
### of SIGNALS cells: 1084
A
B
C
D
E
F
G
H
Cluster
1 2 3 4 5 6 7 8 9 10 11 12 13 14 15 16 17 18 19 20 21 22 X
1 2 3 4 5 6 7 8 9 10 11 12 13 14 15 16 17 18 19 20 21 22 X
Cluster
A (36)
B (189)
C (206)
D (187)
E (51)
F (151)
G (134)
H (103)
None (27)
Copy Number
0
1
2
3
4
5
6
7
8
9
10
11+
Allelic Imbalance
A−Hom
B−Hom
A−Gained
B−Gained
Balanced

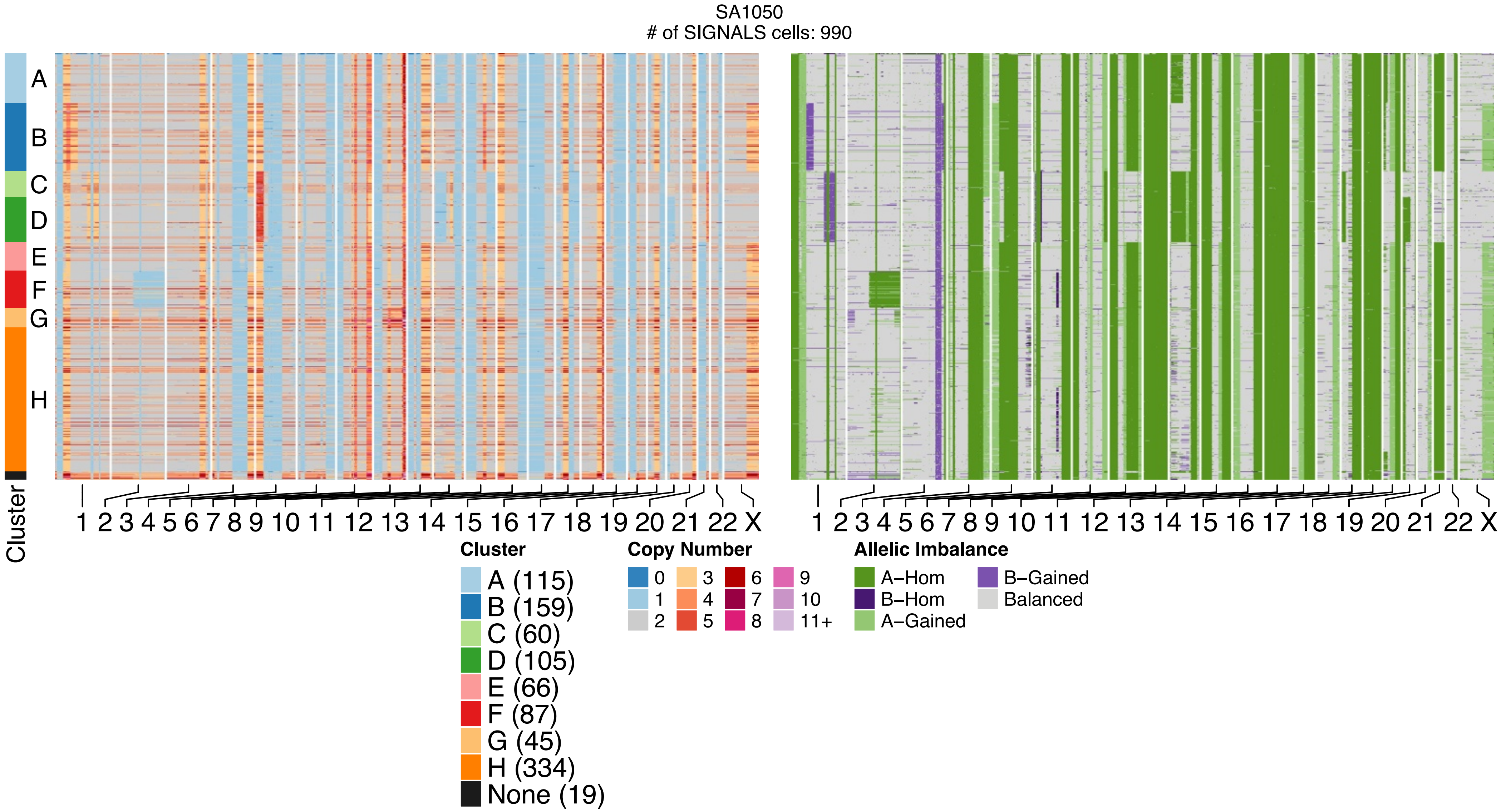
SA1050
### of SIGNALS cells: 990
A
B
C
D
E
F
G
H
Cluster
1 2 3 4 5 6 7 8 9 10 11 12 13 14 15 16 17 18 19 20 21 22 X
1 2 3 4 5 6 7 8 9 10 11 12 13 14 15 16 17 18 19 20 21 22 X
Cluster
A (115)
B (159)
C (60)
D (105)
E (66)
F (87)
G (45)
H (334)
None (19)
Copy Number
0 1 2 3 4 5 6 7 8 9 10 11+
Allelic Imbalance
A−Hom
B−Hom
A−Gained
B−Gained
Balanced

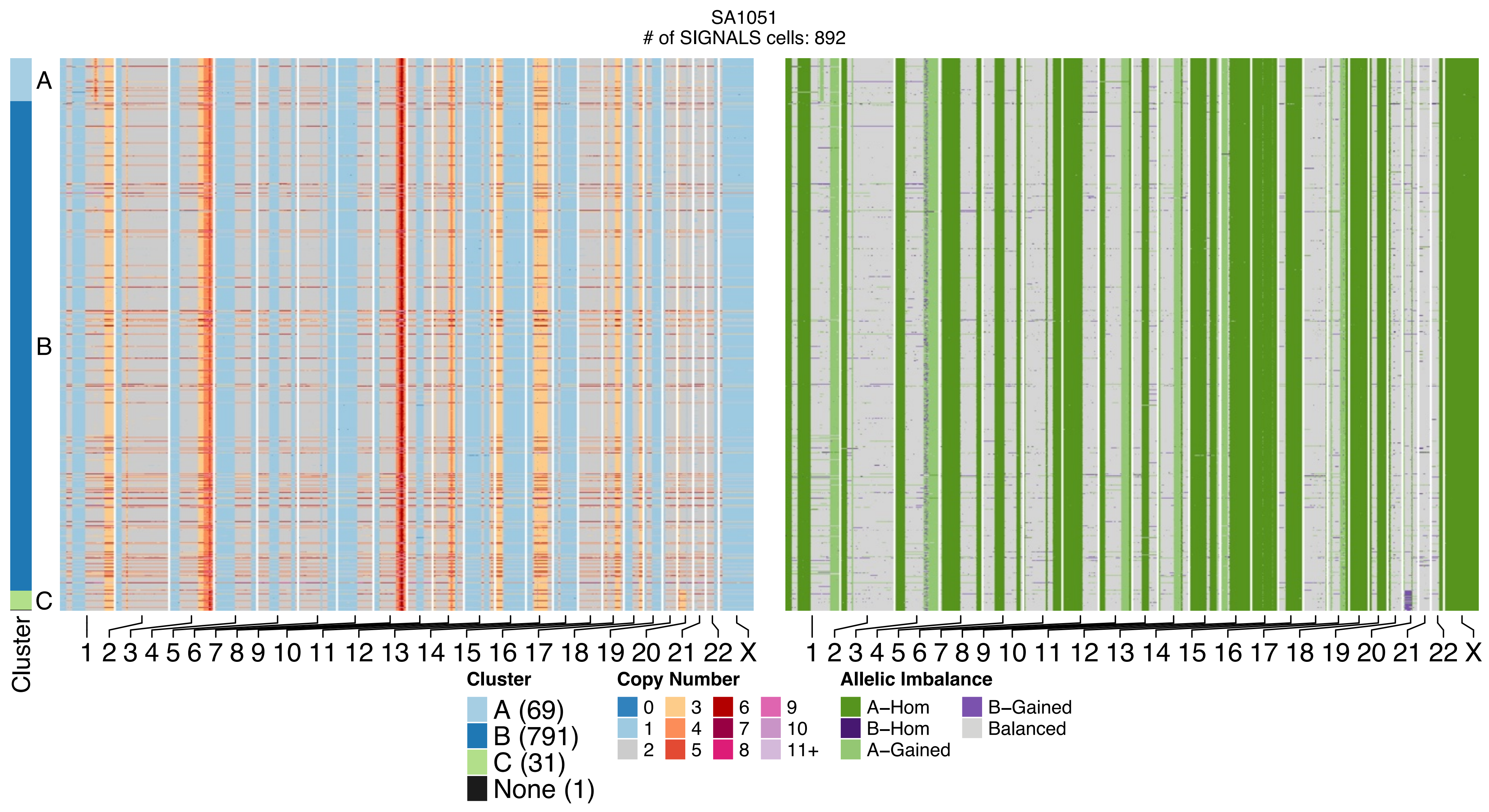

SA1051
### of SIGNALS cells: 892
A
B
C
Cluster
1 2 3 4 5 6 7 8 9 10 11 12 13 14 15 16 17 18 19 20 21 22 X
1 2 3 4 5 6 7 8 9 10 11 12 13 14 15 16 17 18 19 20 21 22 X
Cluster
A (69)
B (791)
C (31)
None (1)
Copy Number
0 1 2 3 4 5 6 7 8 9 10 11+
Allelic Imbalance
A–Hom
B–Hom
A–Gained
B–Gained
Balanced

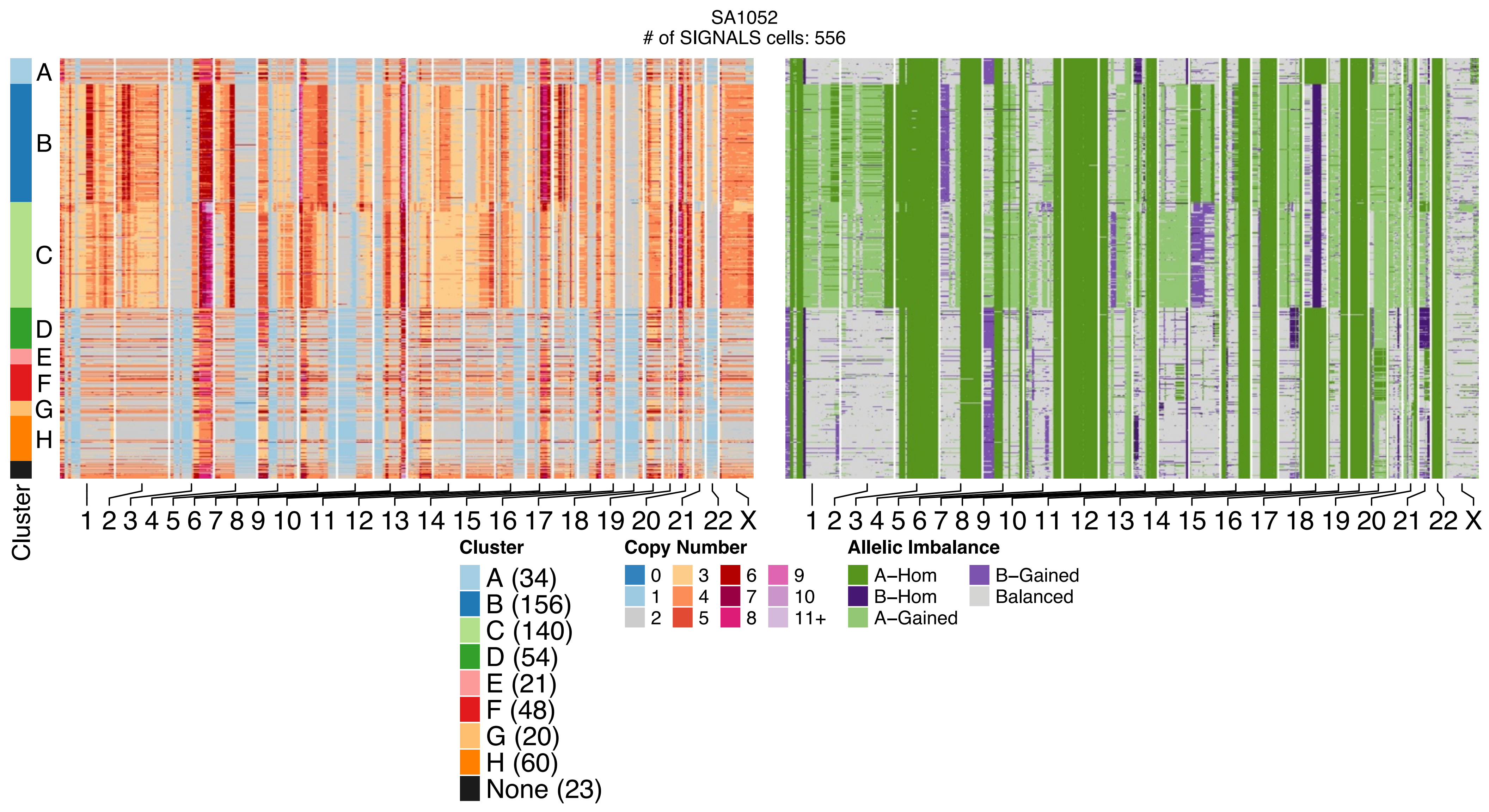
SA1052
### of SIGNALS cells: 556
A
B
C
D
E
F
G
H
Cluster
1 2 3 4 5 6 7 8 9 10 11 12 13 14 15 16 17 18 19 20 21 22 X
1 2 3 4 5 6 7 8 9 10 11 12 13 14 15 16 17 18 19 20 21 22 X
Cluster
A (34)
B (156)
C (140)
D (54)
E (21)
F (48)
G (20)
H (60)
None (23)
Copy Number
0
1
2
3
4
5
6
7
8
9
10
11+
Allelic Imbalance
A–Hom
B–Hom
A–Gained
B–Gained
Balanced

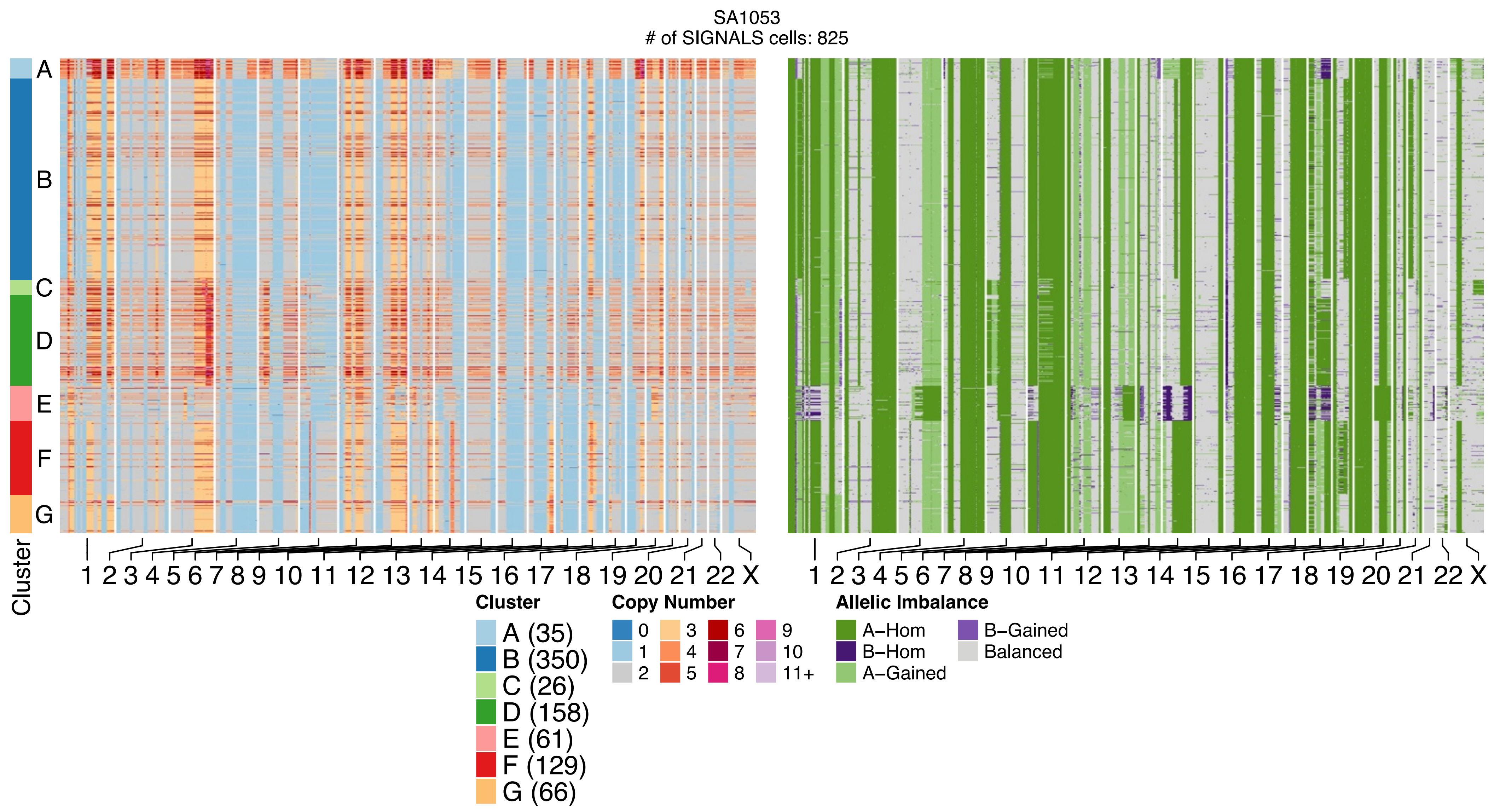
SA1053
### of SIGNALS cells: 825
A
B
C
D
E
F
G
Cluster
1 2 3 4 5 6 7 8 9 10 11 12 13 14 15 16 17 18 19 20 21 22 X
1 2 3 4 5 6 7 8 9 10 11 12 13 14 15 16 17 18 19 20 21 22 X
Cluster
A (35)
B (350)
C (26)
D (158)
E (61)
F (129)
G (66)
Copy Number
0
1
2
3
4
5
6
7
8
9
10
11+
Allelic Imbalance
A–Hom
B–Hom
A–Gained
B–Gained
Balanced

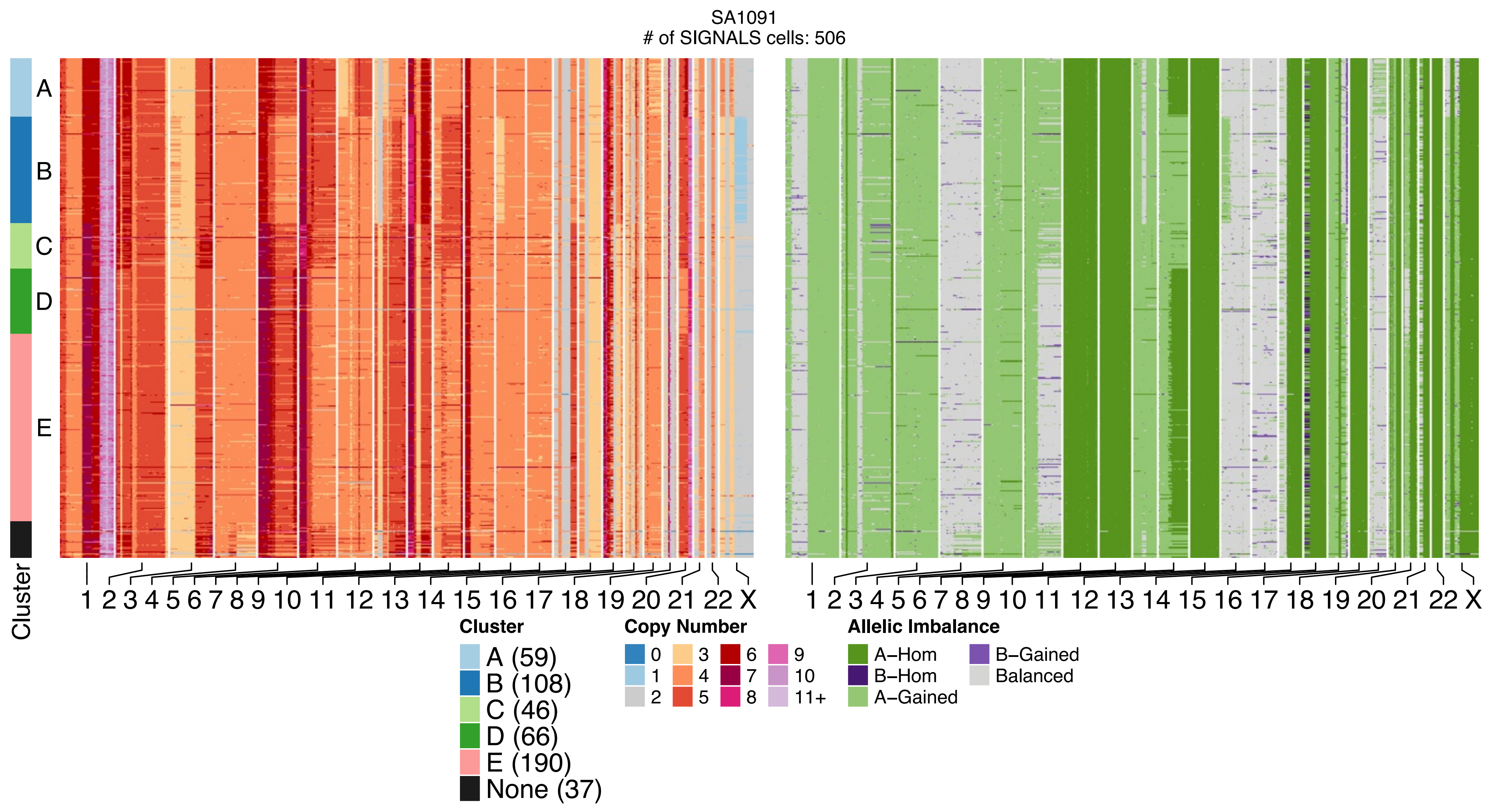

SA1091
### of SIGNALS cells: 506
A
B
C
D
E
Cluster
1 2 3 4 5 6 7 8 9 10 11 12 13 14 15 16 17 18 19 20 21 22 X
1 2 3 4 5 6 7 8 9 10 11 12 13 14 15 16 17 18 19 20 21 22 X
Cluster
A (59)
B (108)
C (46)
D (66)
E (190)
None (37)
Copy Number
0
1
2
3
4
5
6
7
8
9
10
11+
Allelic Imbalance
A–Hom
B–Hom
A–Gained
B–Gained
Balanced

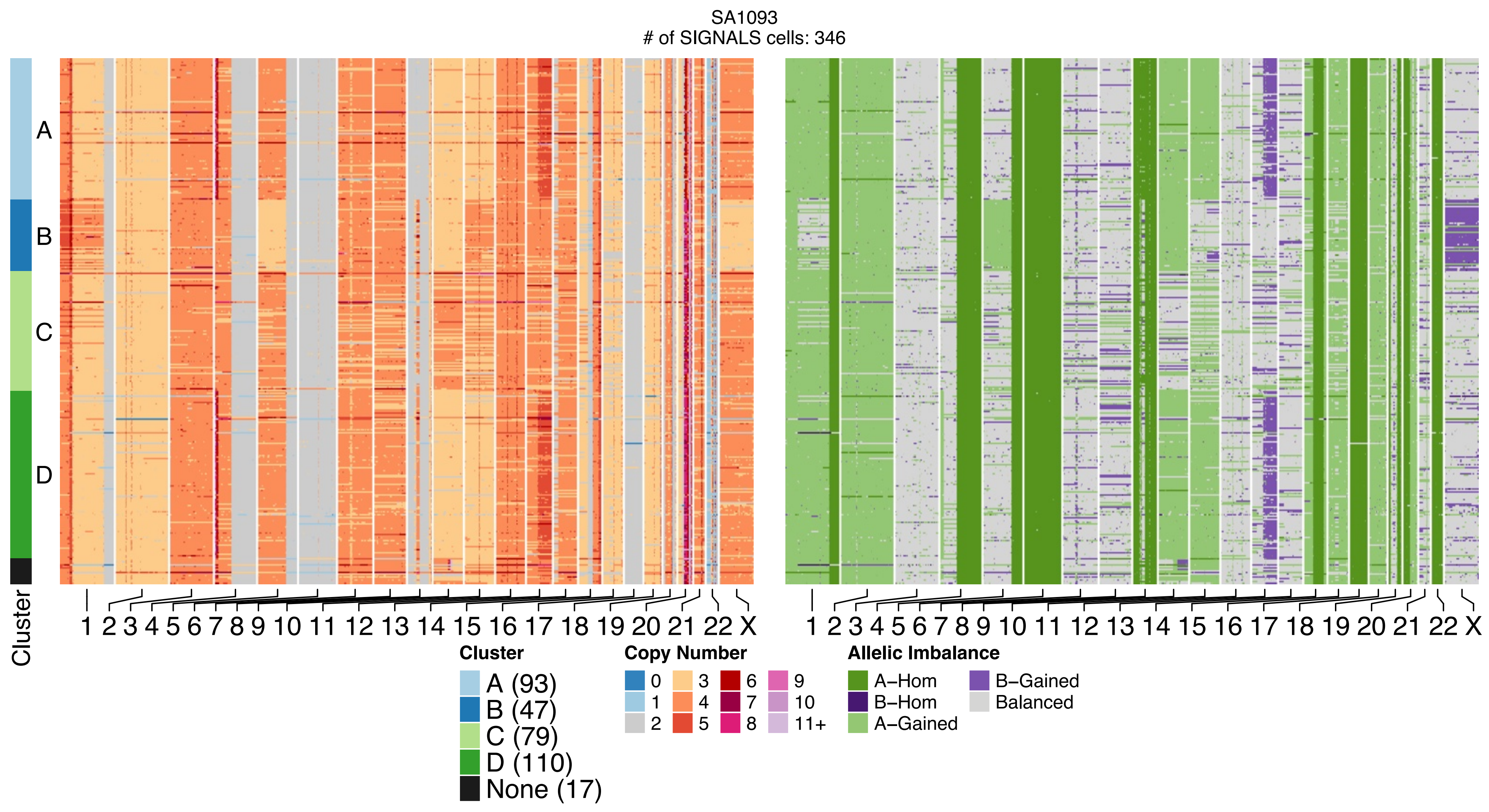
SA1093
### of SIGNALS cells: 346
A
B
C
D
Cluster
1 2 3 4 5 6 7 8 9 10 11 12 13 14 15 16 17 18 19 20 21 22 X
1 2 3 4 5 6 7 8 9 10 11 12 13 14 15 16 17 18 19 20 21 22 X
Cluster
A (93)
B (47)
C (79)
D (110)
None (17)
Copy Number
0
1
2
3
4
5
6
7
8
9
10
11+
Allelic Imbalance
A−Hom
B−Hom
A−Gained
B−Gained
Balanced

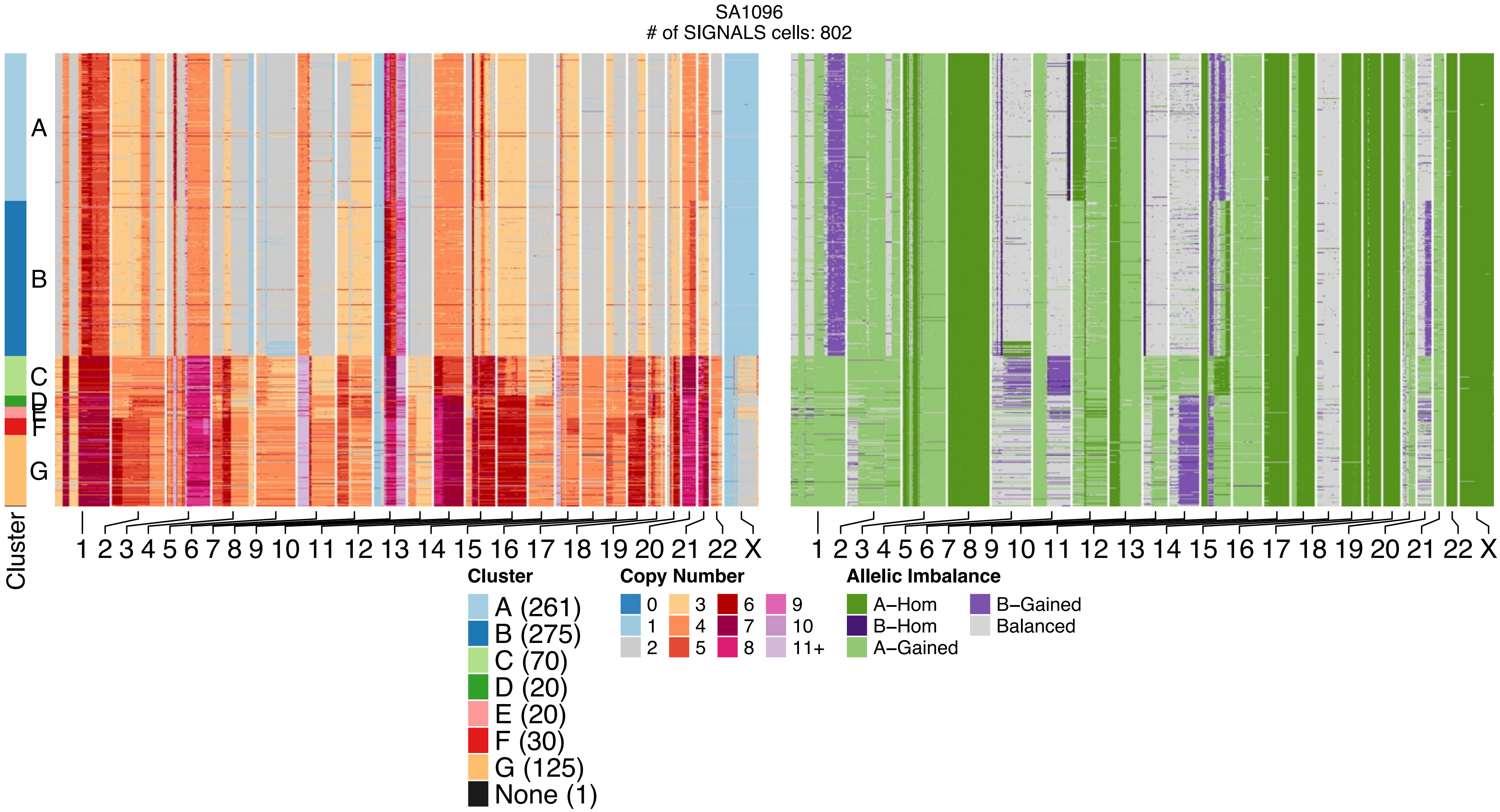

SA1096
### of SIGNALS cells: 802
A
B
C
D
E
F
G
Cluster
1 2 3 4 5 6 7 8 9 10 11 12 13 14 15 16 17 18 19 20 21 22 X
1 2 3 4 5 6 7 8 9 10 11 12 13 14 15 16 17 18 19 20 21 22 X
Cluster
A (261)
B (275)
C (70)
D (20)
E (20)
F (30)
G (125)
None (1)
Copy Number
0
1
2
3
4
5
6
7
8
9
10
11+
Allelic Imbalance
A–Hom
B–Hom
A–Gained
B–Gained
Balanced

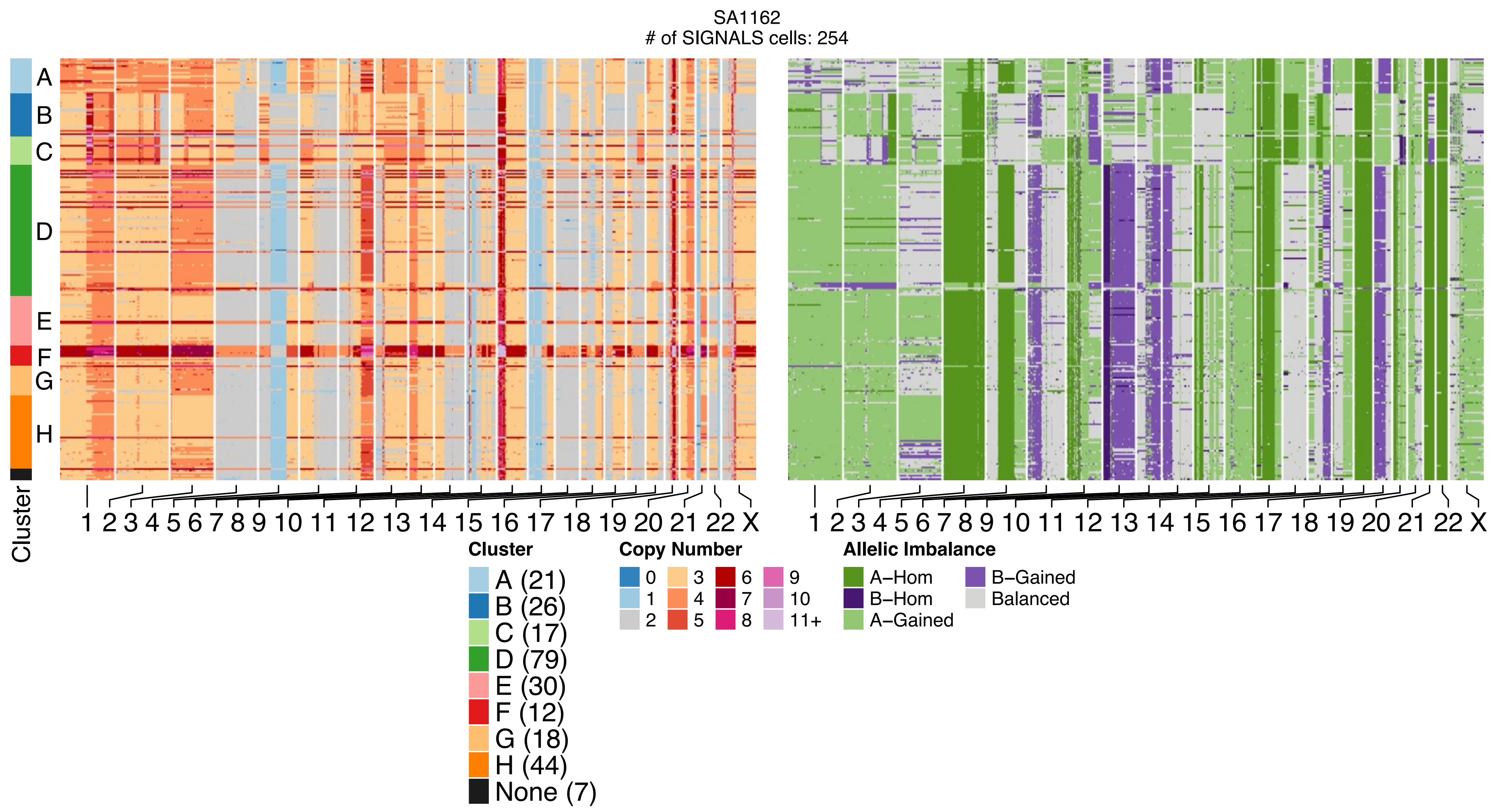

SA1162
### of SIGNALS cells: 254
Cluster
A
B
C
D
E
F
G
H
1 2 3 4 5 6 7 8 9 10 11 12 13 14 15 16 17 18 19 20 21 22 X
1 2 3 4 5 6 7 8 9 10 11 12 13 14 15 16 17 18 19 20 21 22 X
Cluster
A (21)
B (26)
C (17)
D (79)
E (30)
F (12)
G (18)
H (44)
None (7)
Copy Number
0 1 2 3 4 5 6 7 8 9 10 11+
Allelic Imbalance
A−Hom
B−Hom
A−Gained
B−Gained
Balanced

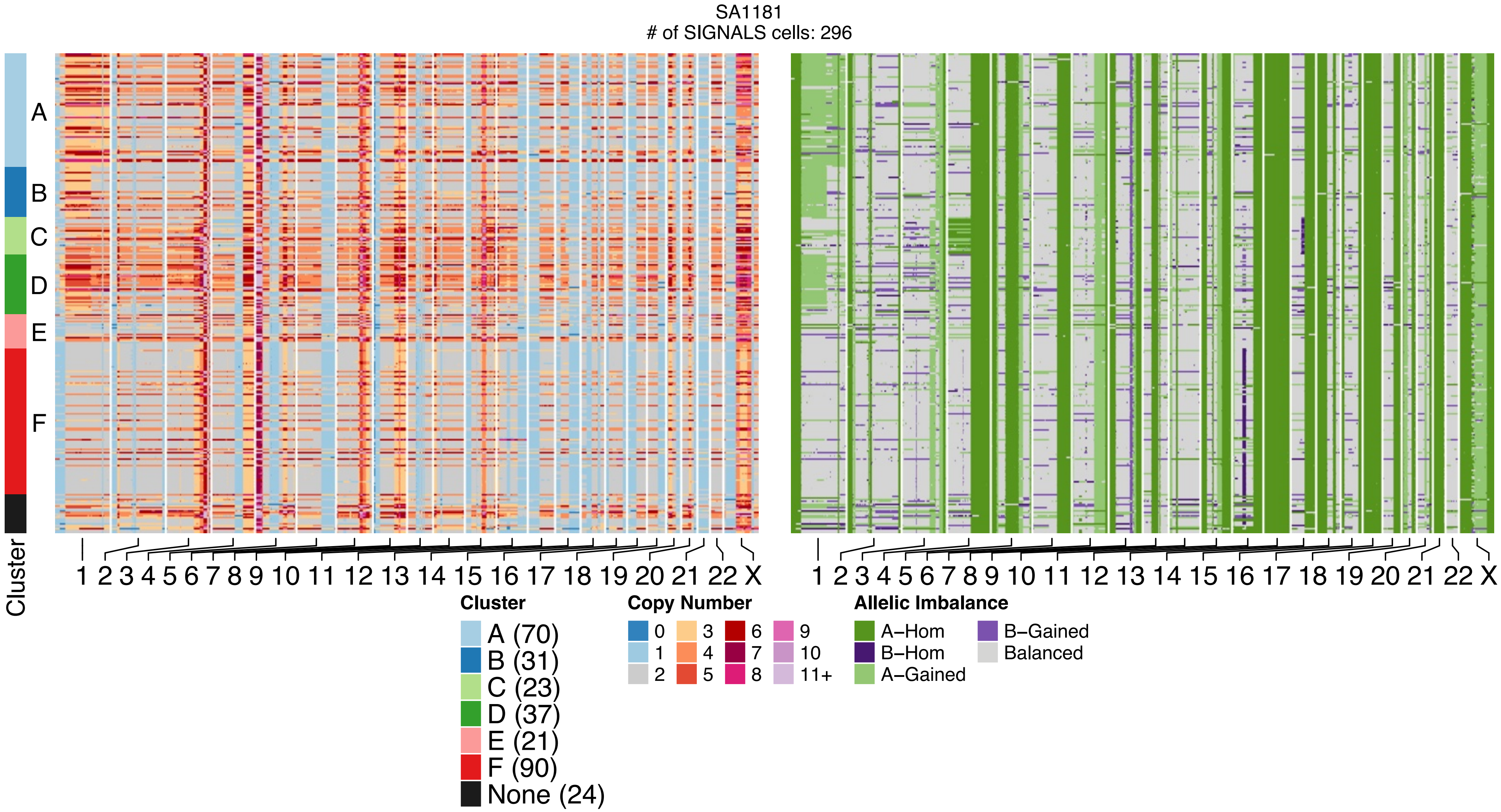
SA1181
### of SIGNALS cells: 296
Cluster
A
B
C
D
E
F
1 2 3 4 5 6 7 8 9 10 11 12 13 14 15 16 17 18 19 20 21 22 X
1 2 3 4 5 6 7 8 9 10 11 12 13 14 15 16 17 18 19 20 21 22 X
Cluster
A (70)
B (31)
C (23)
D (37)
E (21)
F (90)
None (24)
Copy Number
0
1
2
3
4
5
6
7
8
9
10
11+
Allelic Imbalance
A–Hom
B–Hom
A–Gained
B–Gained
Balanced

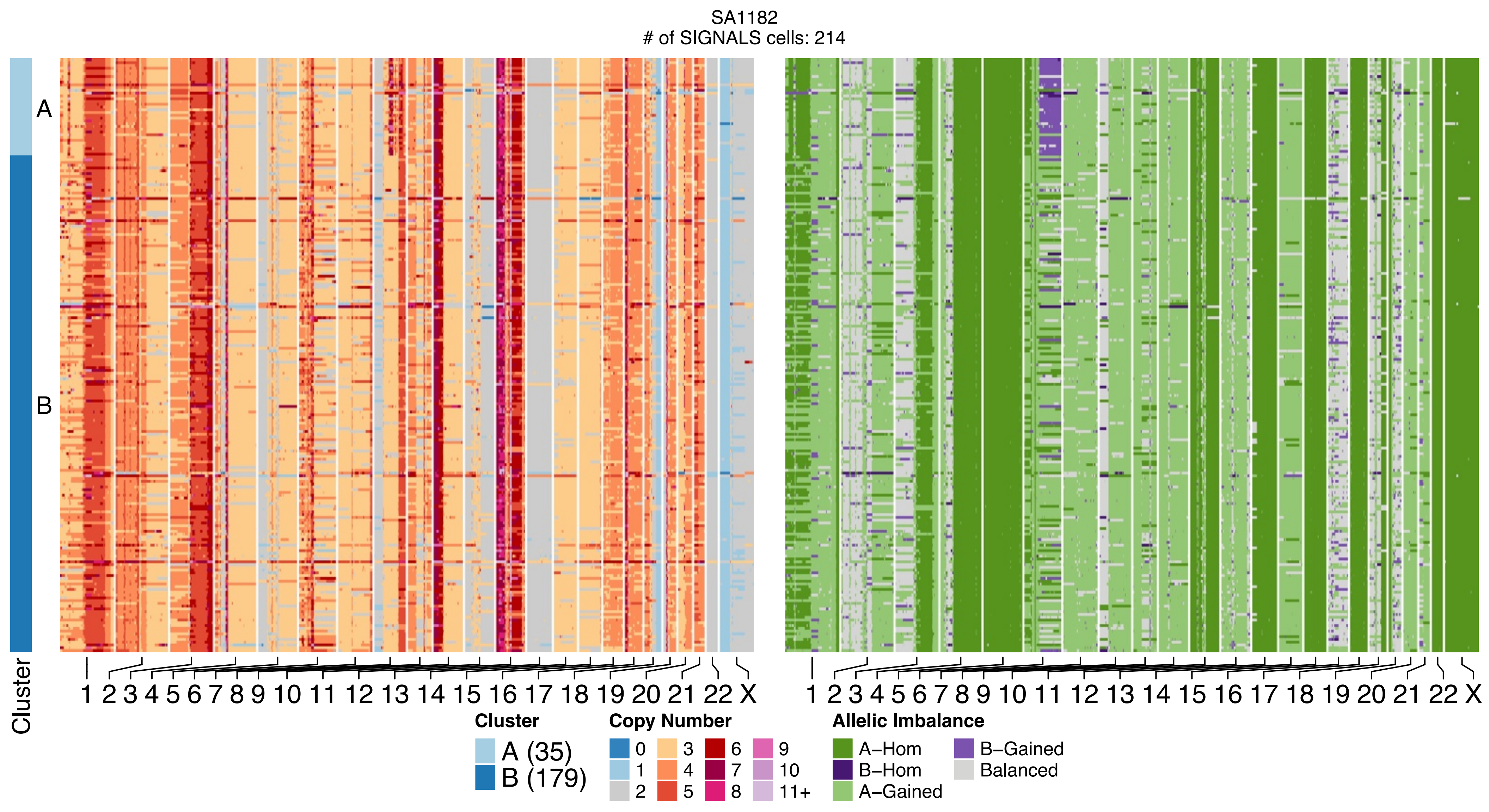
SA1182
### of SIGNALS cells: 214
A
B
Cluster
1 2 3 4 5 6 7 8 9 10 11 12 13 14 15 16 17 18 19 20 21 22 X
1 2 3 4 5 6 7 8 9 10 11 12 13 14 15 16 17 18 19 20 21 22 X
Cluster
A (35)
B (179)
Copy Number
0
1
2
3
4
5
6
7
8
9
10
11+
Allelic Imbalance
A−Hom
B−Hom
A−Gained
B−Gained
Balanced

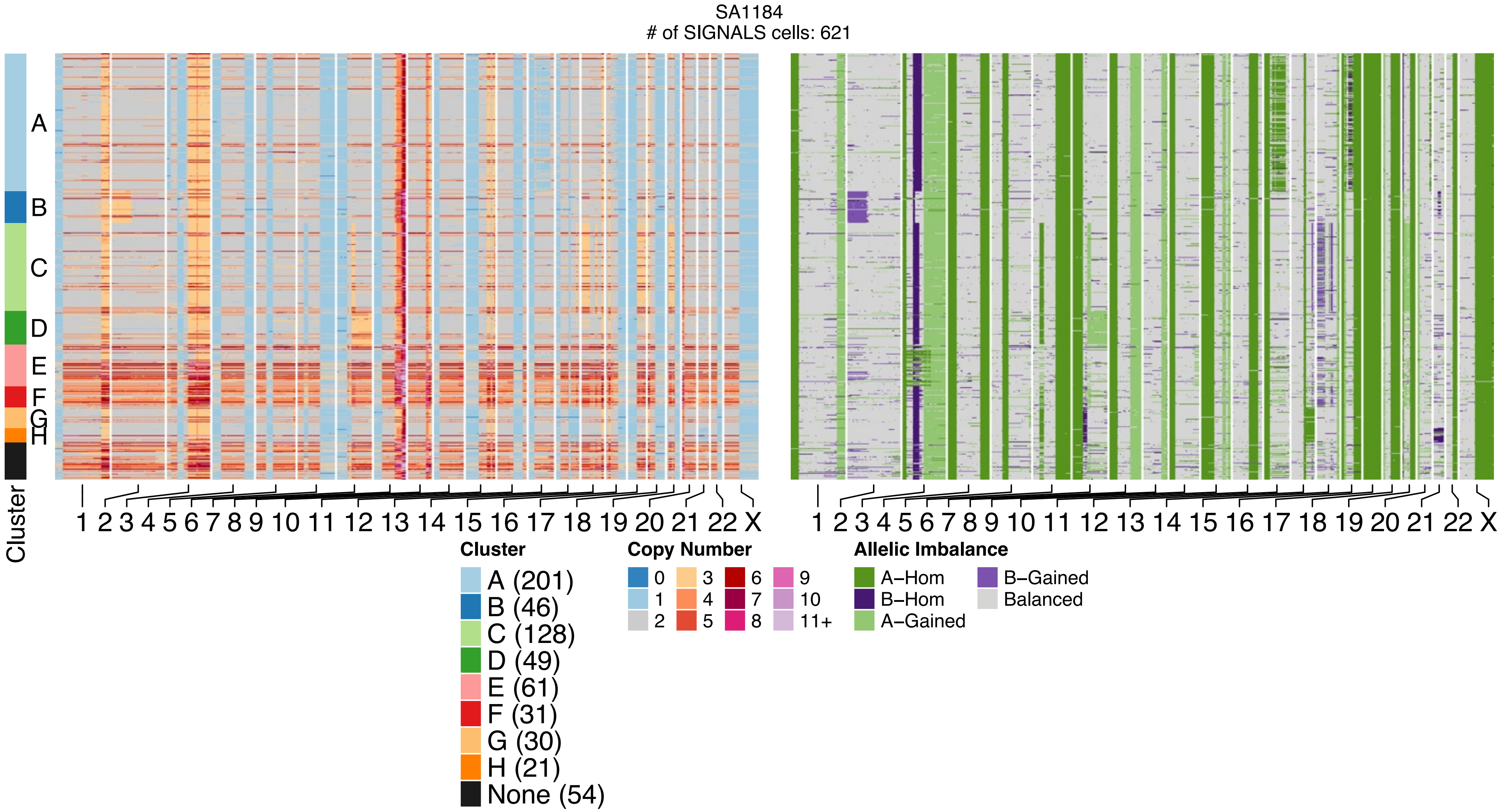
SA1184
### of SIGNALS cells: 621
A
B
C
D
E
F
G
H
Cluster
1 2 3 4 5 6 7 8 9 10 11 12 13 14 15 16 17 18 19 20 21 22 X
1 2 3 4 5 6 7 8 9 10 11 12 13 14 15 16 17 18 19 20 21 22 X
Cluster
A (201)
B (46)
C (128)
D (49)
E (61)
F (31)
G (30)
H (21)
None (54)
Copy Number
0
1
2
3
4
5
6
7
8
9
10
11+
Allelic Imbalance
A−Hom
B−Hom
A−Gained
B−Gained
Balanced

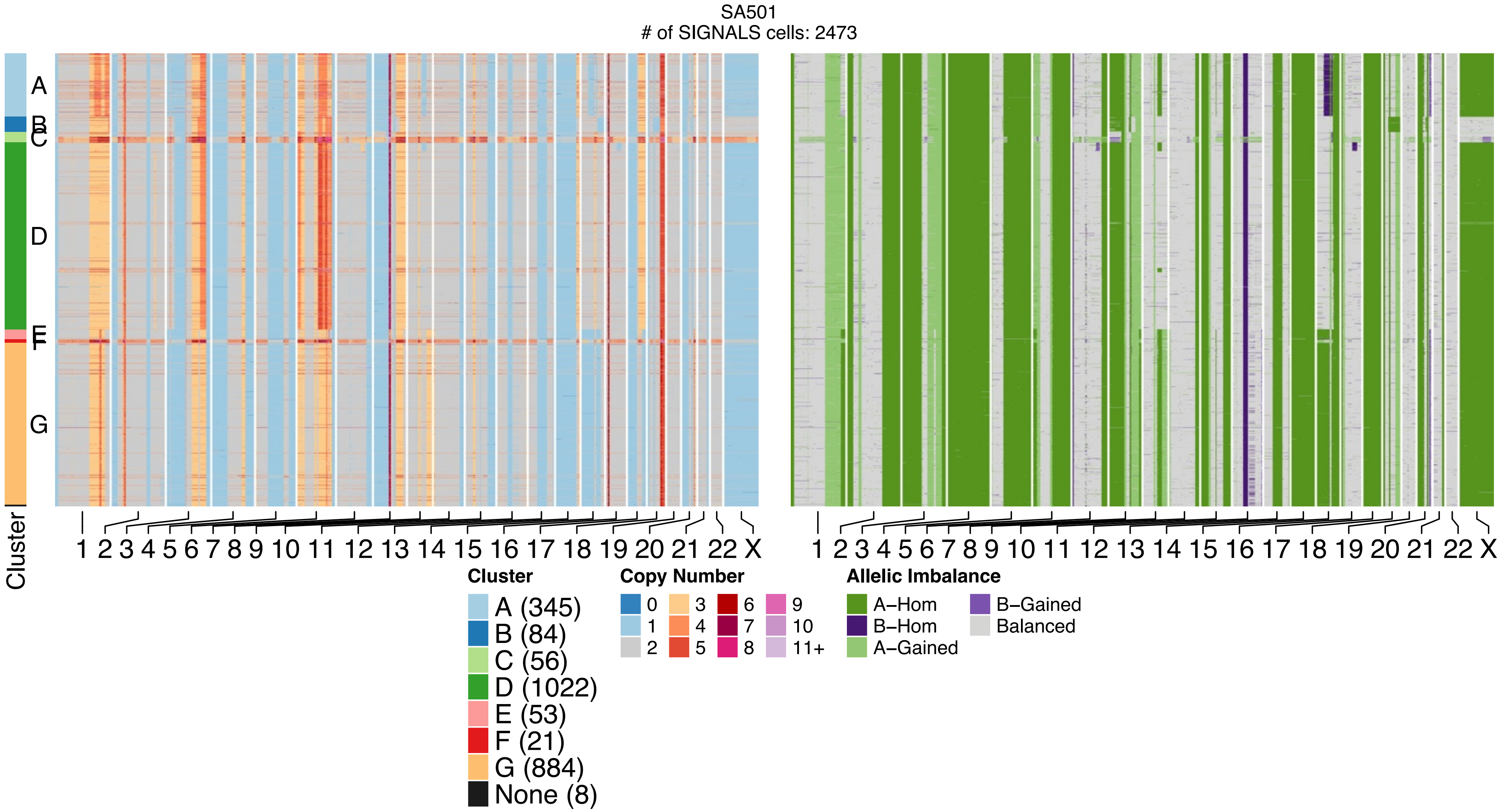
SA501
### of SIGNALS cells: 2473
A
B
C
D
E
F
G
Cluster
1 2 3 4 5 6 7 8 9 10 11 12 13 14 15 16 17 18 19 20 21 22 X
1 2 3 4 5 6 7 8 9 10 11 12 13 14 15 16 17 18 19 20 21 22 X
Cluster
A (345)
B (84)
C (56)
D (1022)
E (53)
F (21)
G (884)
None (8)
Copy Number
0
1
2
3
4
5
6
7
8
9
10
11+
Allelic Imbalance
A–Hom
B–Hom
A–Gained
B–Gained
Balanced

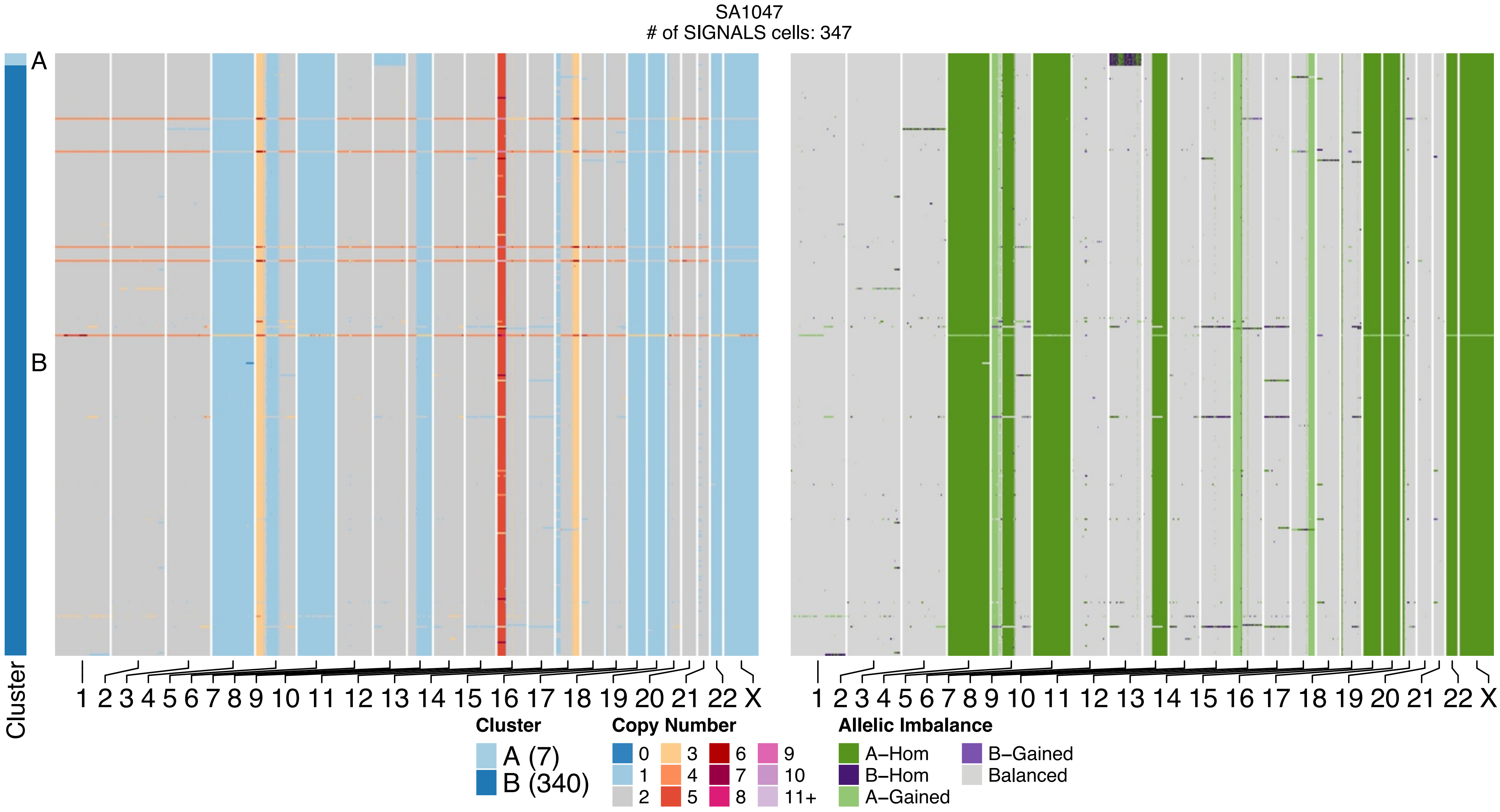
SA1047
### of SIGNALS cells: 347
A
B
Cluster
1 2 3 4 5 6 7 8 9 10 11 12 13 14 15 16 17 18 19 20 21 22 X
1 2 3 4 5 6 7 8 9 10 11 12 13 14 15 16 17 18 19 20 21 22 X
Cluster
A (7)
B (340)
Copy Number
0
1
2
3
4
5
6
7
8
9
10
11+
Allelic Imbalance
A–Hom
B–Hom
A–Gained
B–Gained
Balanced

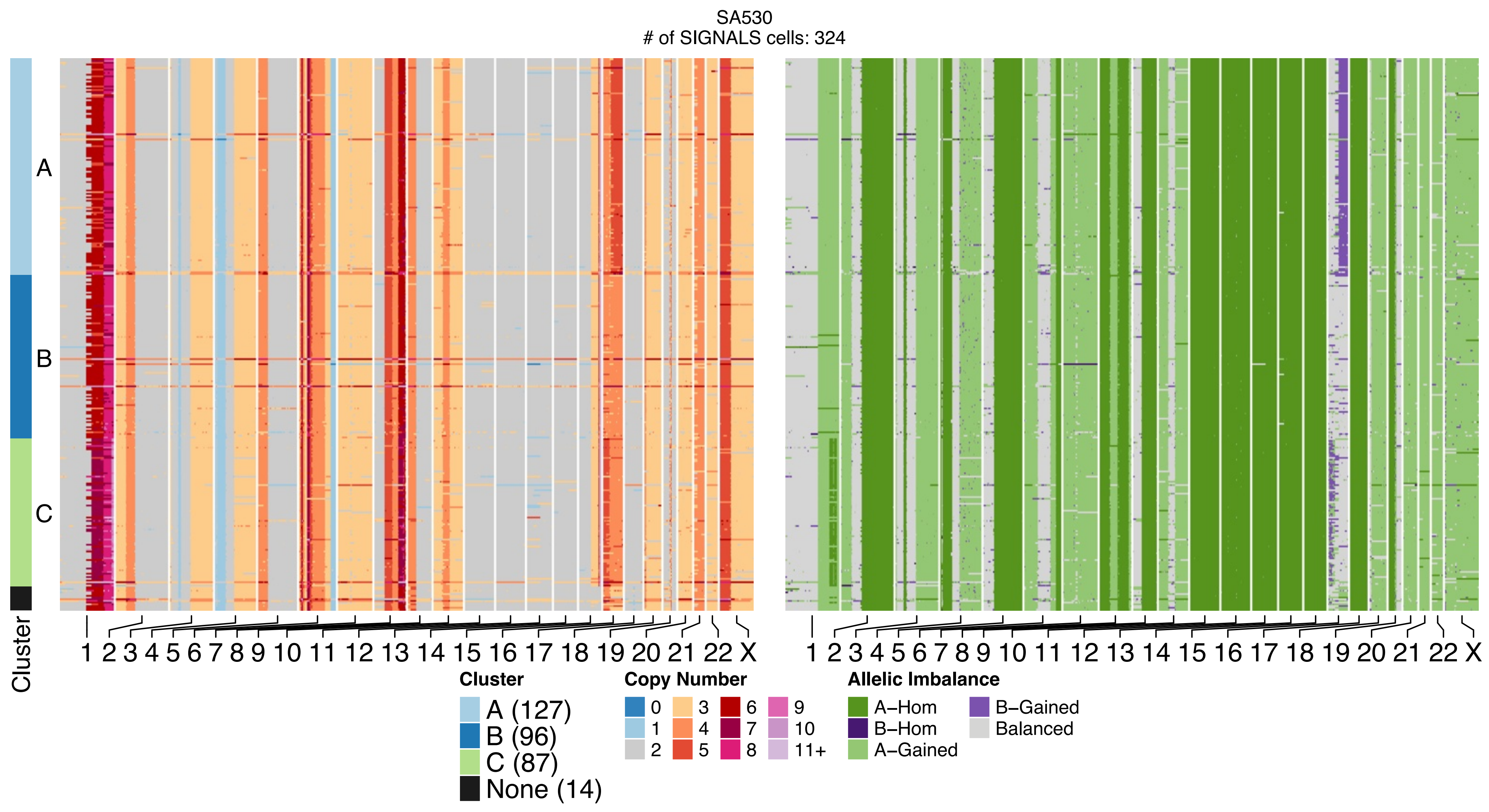

SA530
### of SIGNALS cells: 324
A
B
C
Cluster
1 2 3 4 5 6 7 8 9 10 11 12 13 14 15 16 17 18 19 20 21 22 X
1 2 3 4 5 6 7 8 9 10 11 12 13 14 15 16 17 18 19 20 21 22 X
Cluster
A (127)
B (96)
C (87)
None (14)
Copy Number
0
1
2
3
4
5
6
7
8
9
10
11+
Allelic Imbalance
A−Hom
B−Hom
A−Gained
B−Gained
Balanced

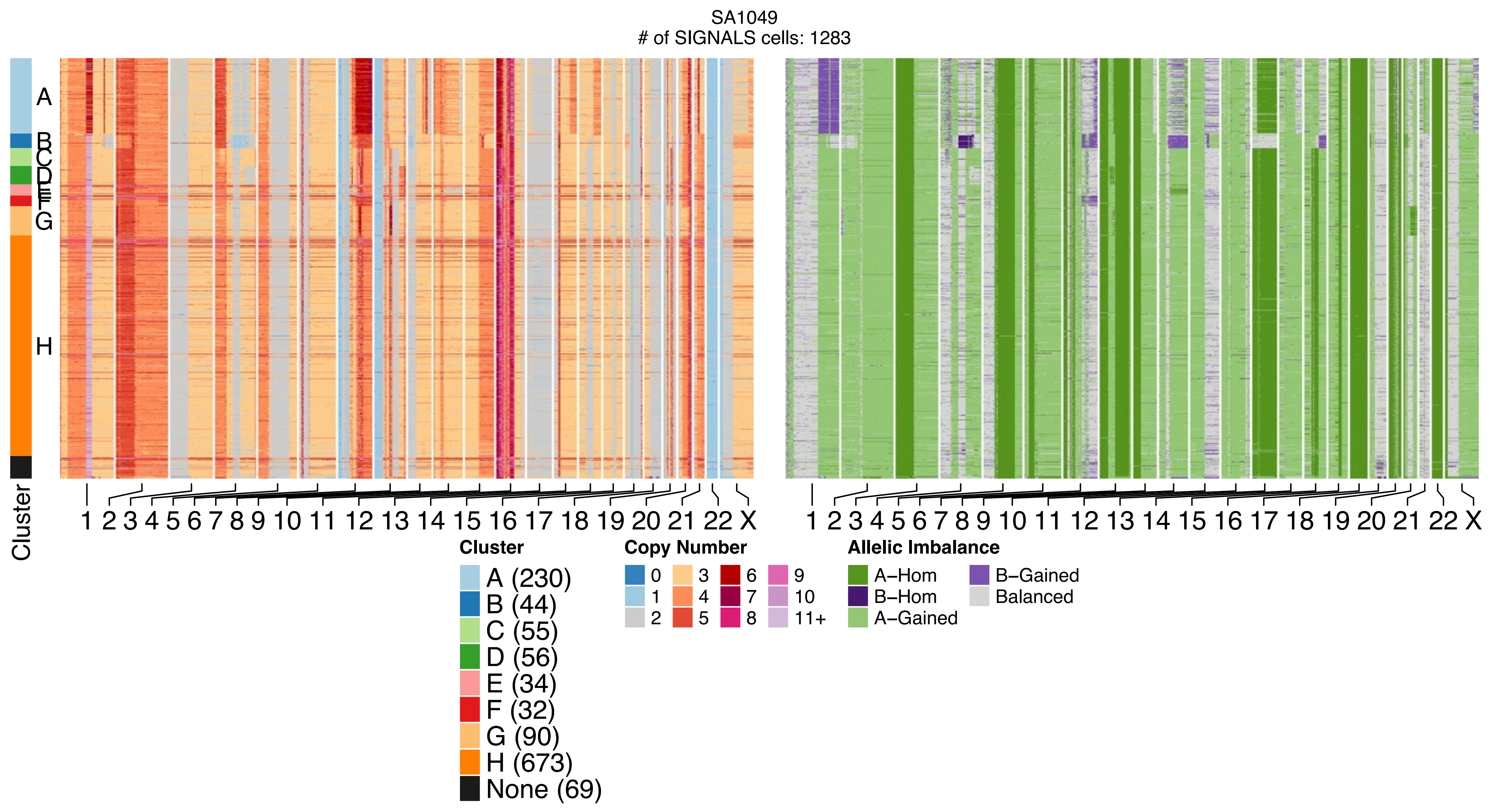
SA1049
### of SIGNALS cells: 1283
Cluster
A
B
C
D
E
F
G
H
1 2 3 4 5 6 7 8 9 10 11 12 13 14 15 16 17 18 19 20 21 22 X
Cluster
A (230)
B (44)
C (55)
D (56)
E (34)
F (32)
G (90)
H (673)
None (69)
Copy Number
0
1
2
3
4
5
6
7
8
9
10
11+
Allelic Imbalance
A−Hom
B−Hom
A−Gained
B−Gained
Balanced

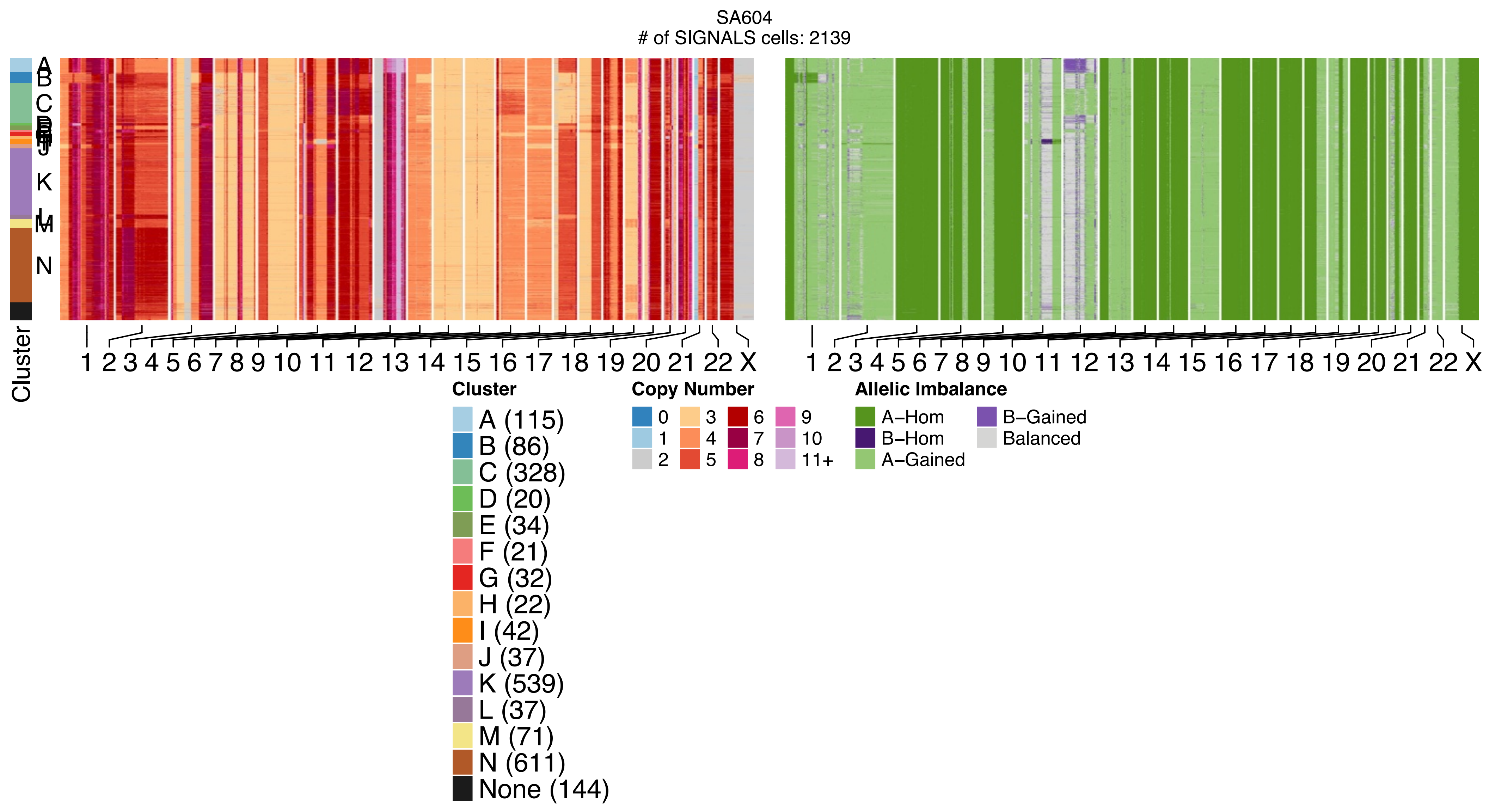
SA604
### of SIGNALS cells: 2139
Cluster
A
B
C
K
L
M
N
1 2 3 4 5 6 7 8 9 10 11 12 13 14 15 16 17 18 19 20 21 22 X
1 2 3 4 5 6 7 8 9 10 11 12 13 14 15 16 17 18 19 20 21 22 X
Cluster
A (115)
B (86)
C (328)
D (20)
E (34)
F (21)
G (32)
H (22)
I (42)
J (37)
K (539)
L (37)
M (71)
N (611)
None (144)
Copy Number
0
1
2
3
4
5
6
7
8
9
10
11+
Allelic Imbalance
A−Hom
B−Hom
A−Gained
B−Gained
Balanced

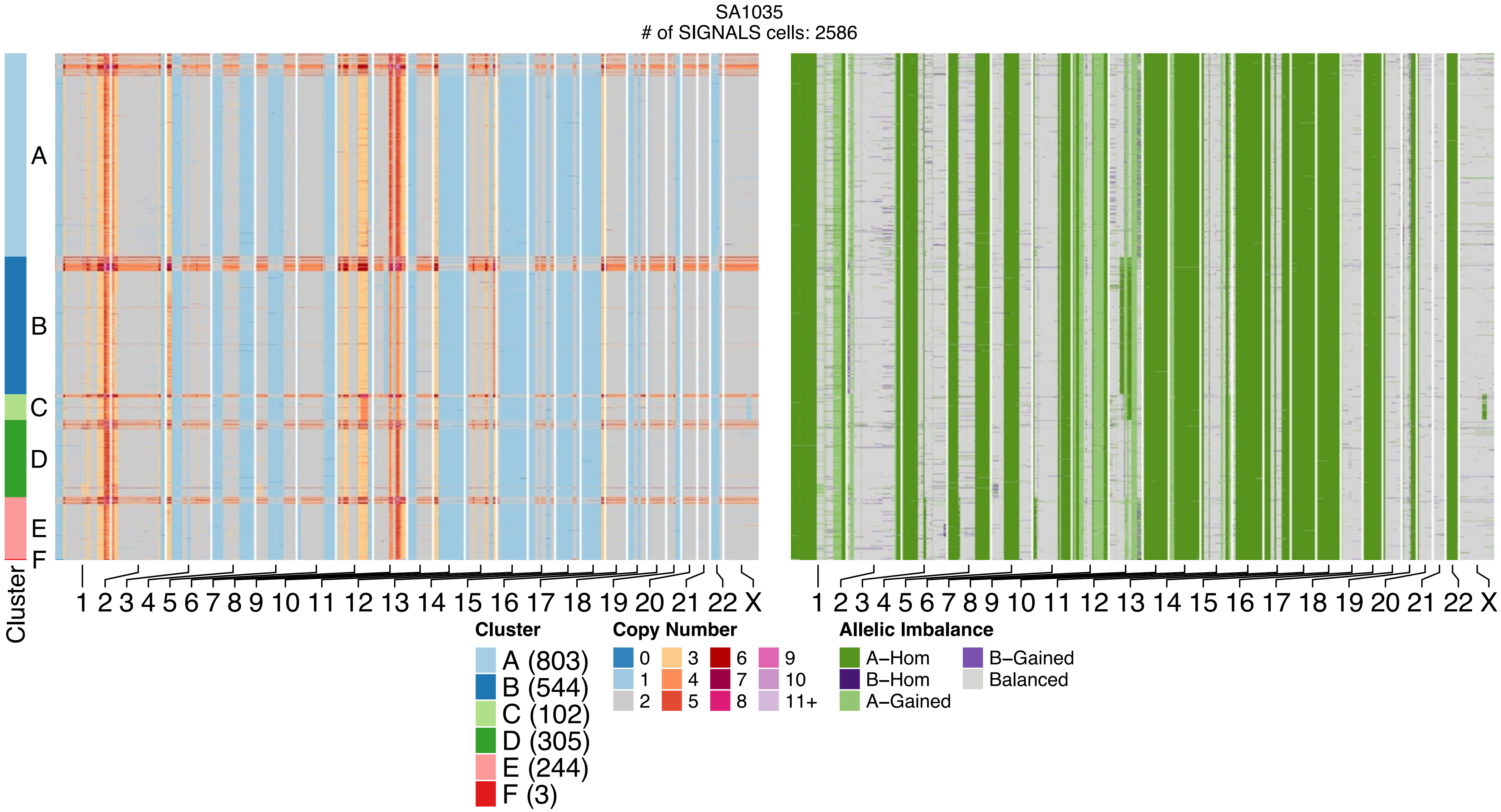
SA1035
### of SIGNALS cells: 2586
A
B
C
D
E
F
Cluster
1 2 3 4 5 6 7 8 9 10 11 12 13 14 15 16 17 18 19 20 21 22 X
1 2 3 4 5 6 7 8 9 10 11 12 13 14 15 16 17 18 19 20 21 22 X
Cluster
A (803)
B (544)
C (102)
D (305)
E (244)
F (3)
Copy Number
0
1
2
3
4
5
6
7
8
9
10
11+
Allelic Imbalance
A–Hom
B–Hom
A–Gained
B–Gained
Balanced

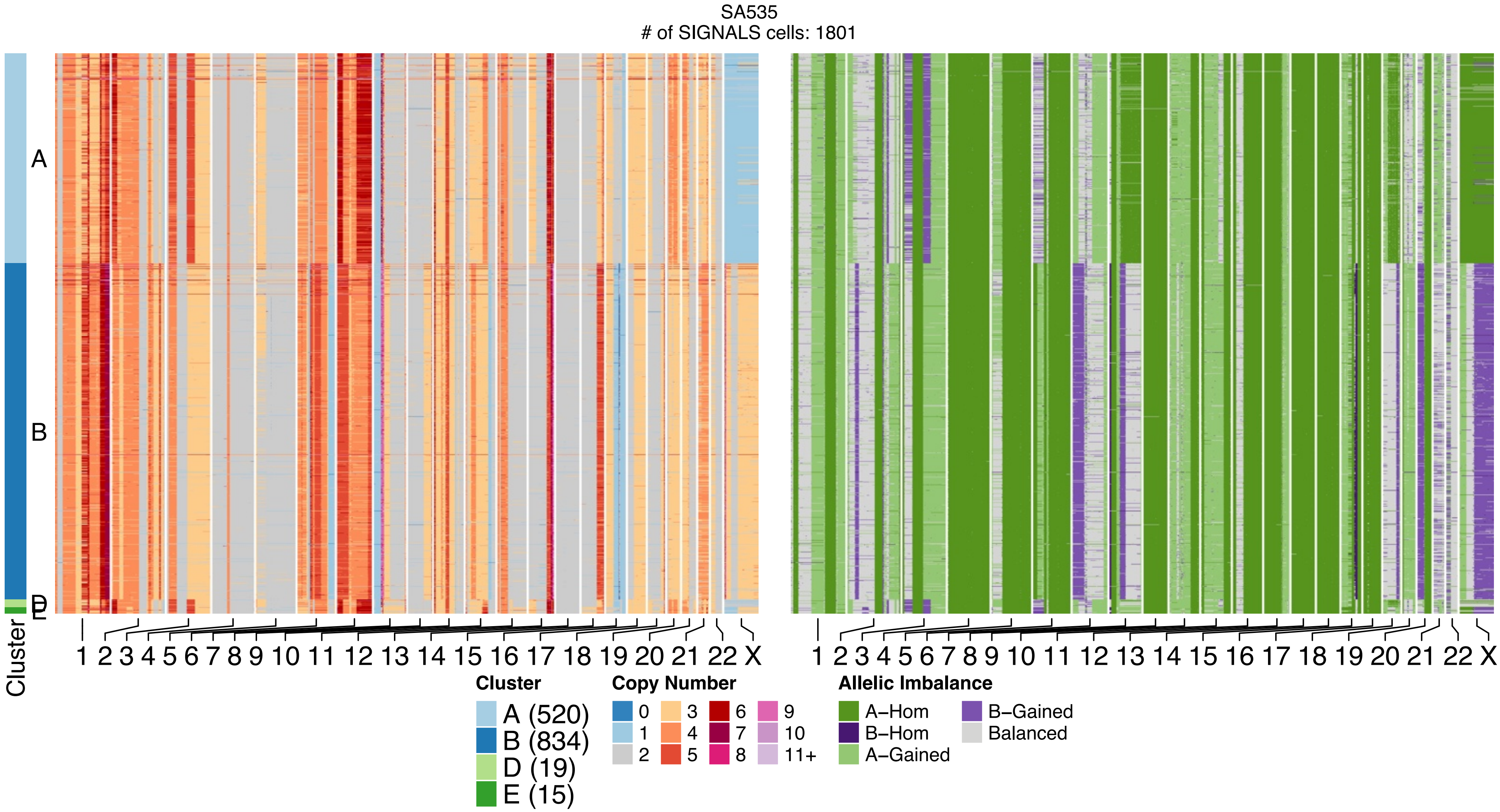

SA535
### of SIGNALS cells: 1801
A
B
D
E
Cluster
1 2 3 4 5 6 7 8 9 10 11 12 13 14 15 16 17 18 19 20 21 22 X
1 2 3 4 5 6 7 8 9 10 11 12 13 14 15 16 17 18 19 20 21 22 X
Cluster
A (520)
B (834)
D (19)
E (15)
Copy Number
0
1
2
3
4
5
6
7
8
9
10
11+
Allelic Imbalance
A–Hom
B–Hom
A–Gained
B–Gained
Balanced

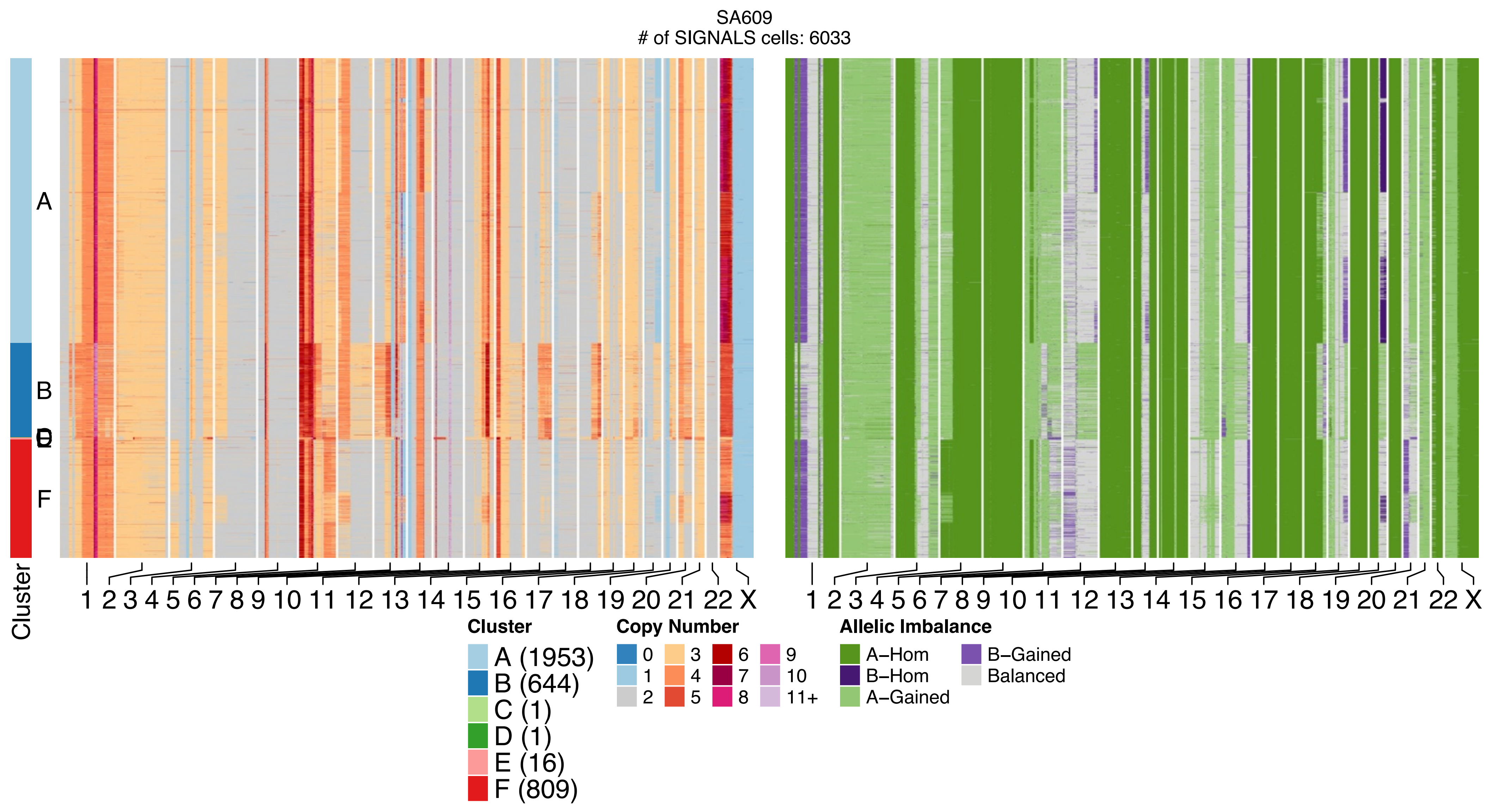
SA609
### of SIGNALS cells: 6033
A
B
D
E
F
Cluster
1 2 3 4 5 6 7 8 9 10 11 12 13 14 15 16 17 18 19 20 21 22 X
1 2 3 4 5 6 7 8 9 10 11 12 13 14 15 16 17 18 19 20 21 22 X
Cluster
A (1953)
B (644)
C (1)
D (1)
E (16)
F (809)
Copy Number
0
1
2
3
4
5
6
7
8
9
10
11+
Allelic Imbalance
A–Hom
B–Hom
A–Gained
B–Gained
Balanced
