## Supplementary figures and images for "Single-cell DNA replication dynamics in genomically unstable cancers"

### Additional File 2

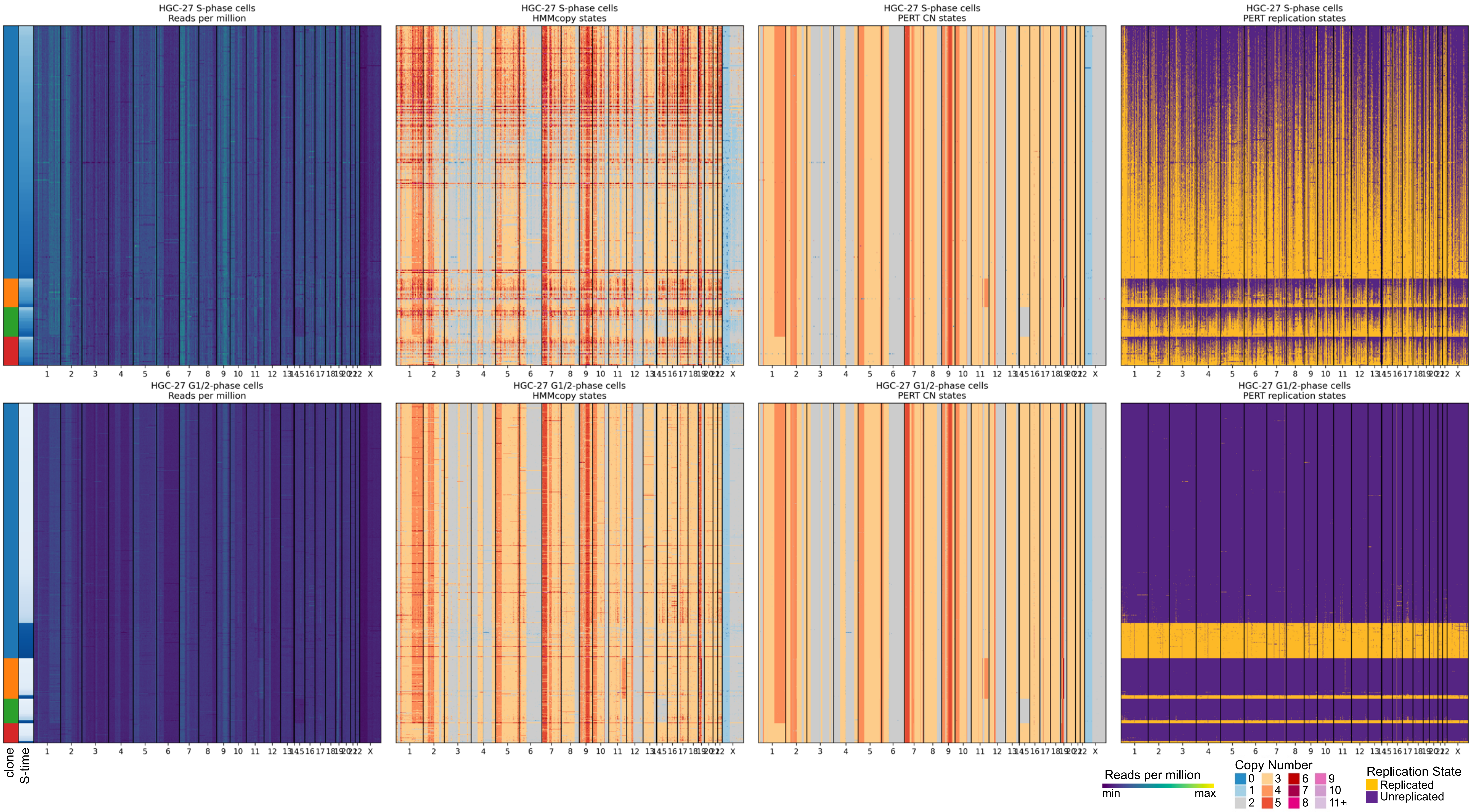

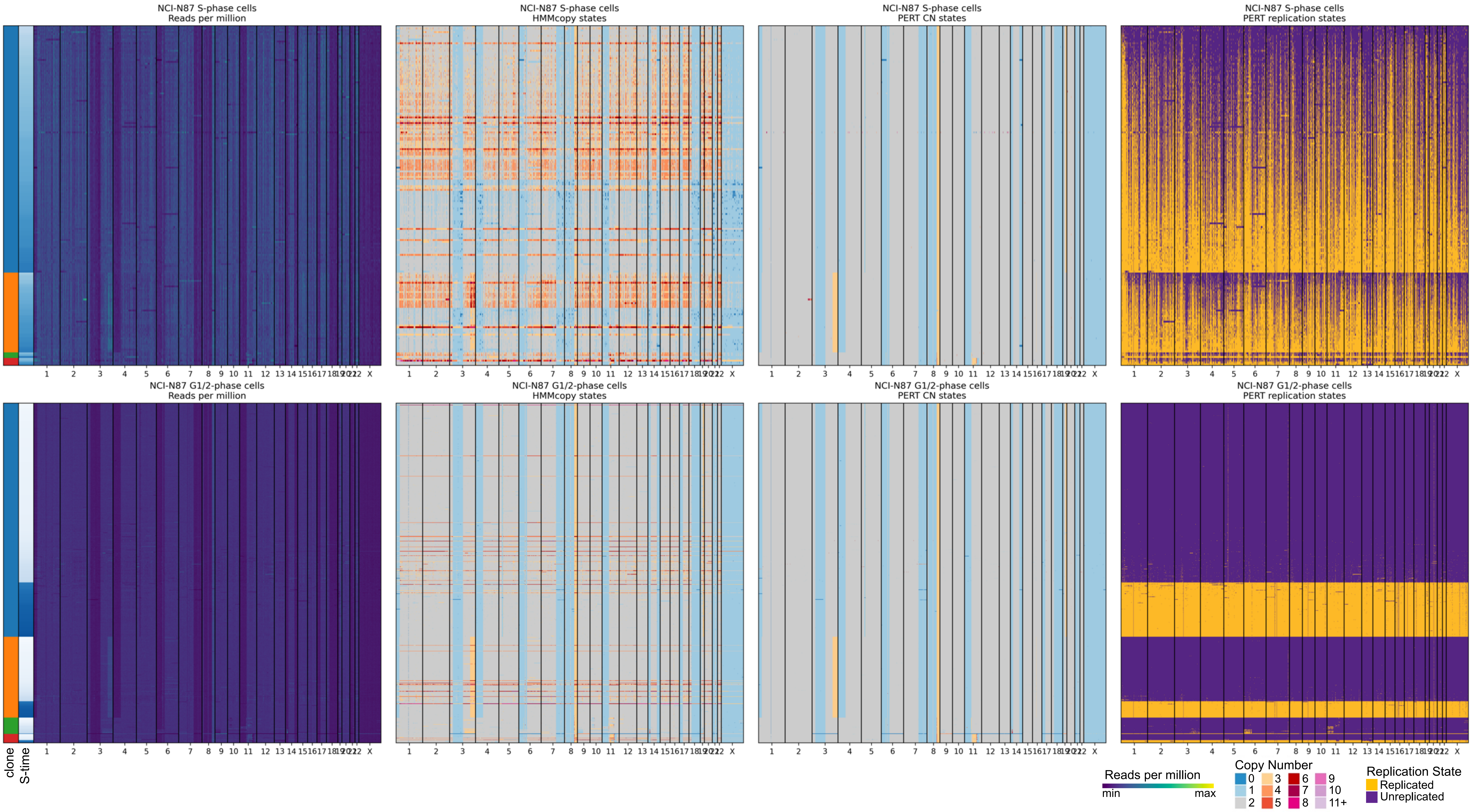

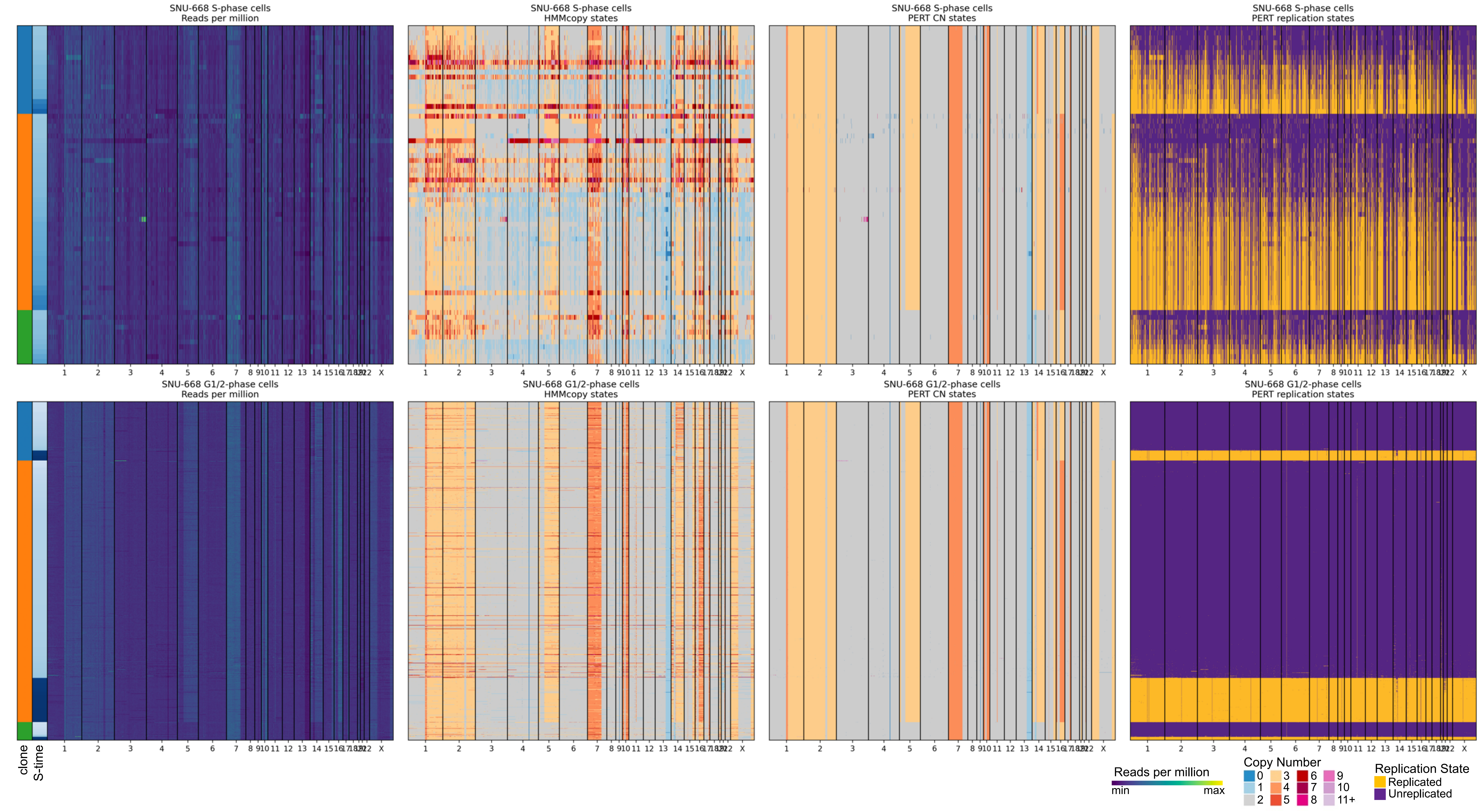

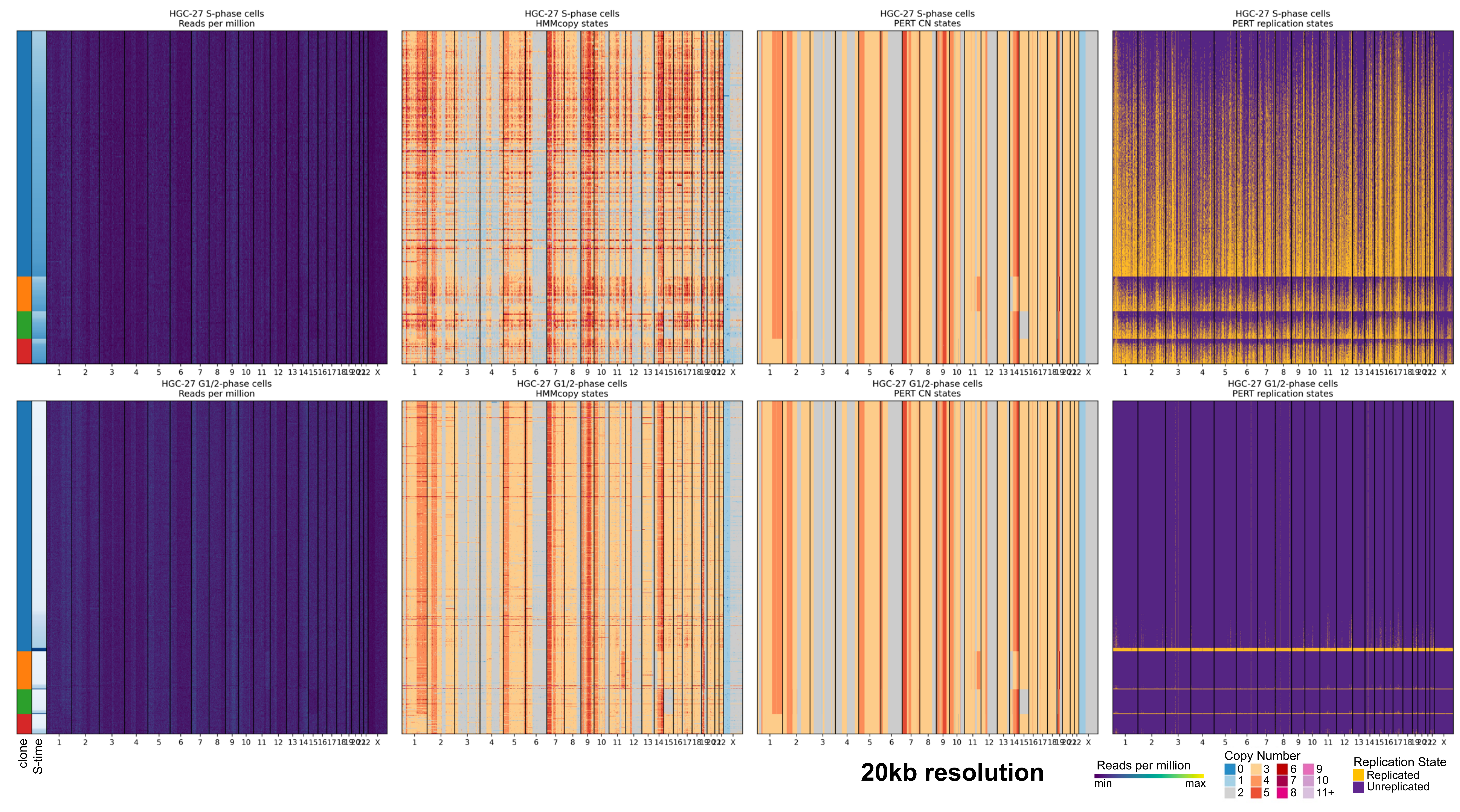

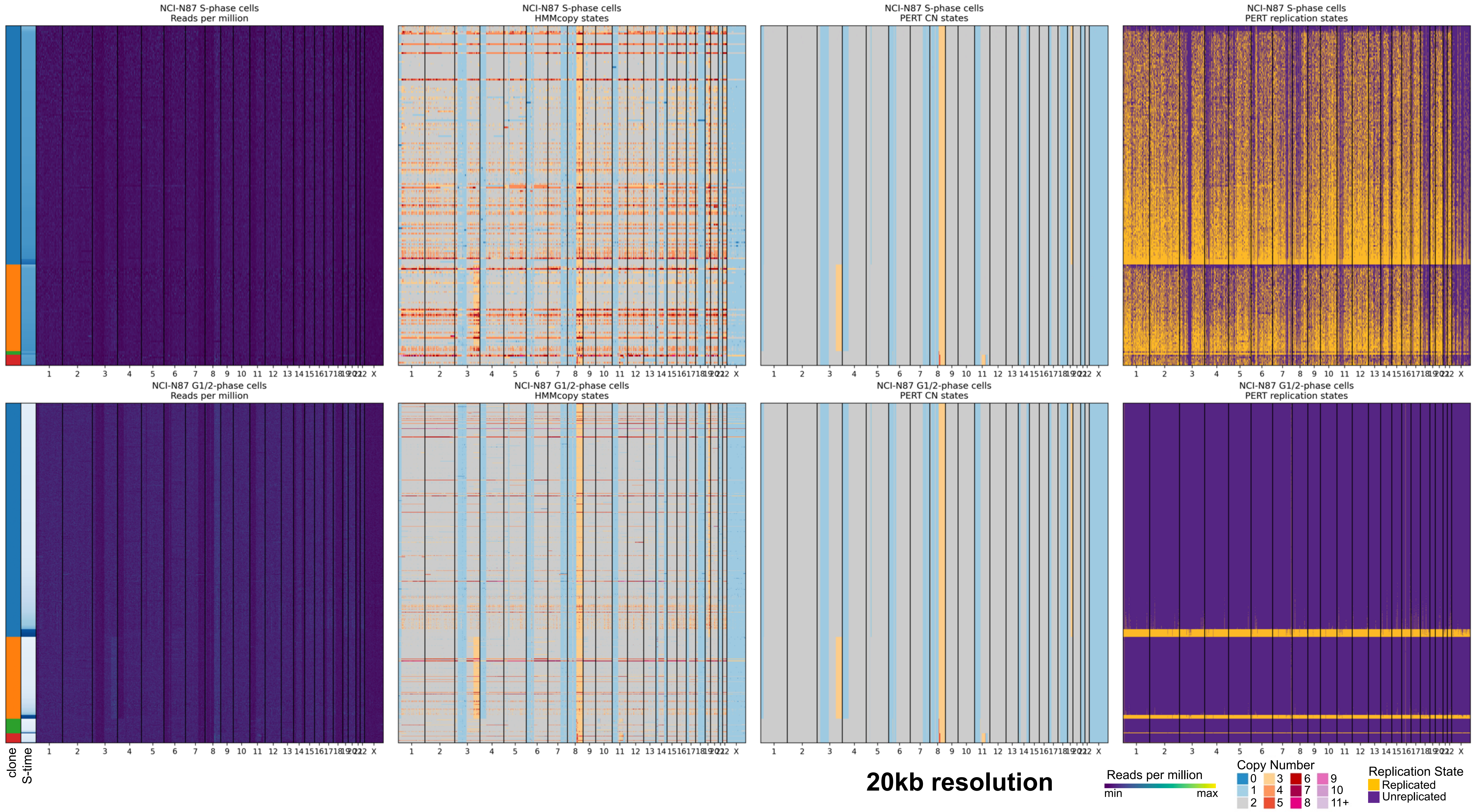
